# Comparative single cell analysis of transcriptional bursting reveals the role of genome organization on *de novo* transcript origination

**DOI:** 10.1101/2024.04.29.591771

**Authors:** UnJin Lee, Cong Li, Christopher B. Langer, Nicolas Svetec, Li Zhao

## Abstract

Spermatogenesis is a key developmental process underlying the origination of newly evolved genes. However, rapid cell type-specific transcriptomic divergence of the *Drosophila* germline has posed a significant technical barrier for comparative single-cell RNA-sequencing (scRNA-Seq) studies. By quantifying a surprisingly strong correlation between species- and cell type-specific divergence in three closely related *Drosophila* species, we apply a new statistical procedure to identify a core set of 198 genes that are highly predictive of cell type identity while remaining robust to species-specific differences that span over 25-30 million years of evolution. We then utilize cell type classifications based on the 198-gene set to show how transcriptional divergence in cell type increases throughout spermatogenic developmental time. After validating these cross-species cell type classifications using RNA fluorescence in situ hybridization (FISH) and imaging, we then investigate the influence of genome organization on the molecular evolution of spermatogenesis vis-a-vis transcriptional bursting. We first show altering transcriptional burst size contributes to pre-meiotic transcription and altering bursting frequency contributes to post-meiotic expression. We then report global differences in autosomal vs. X chromosomal transcription may arise in a developmental stage preceding full testis organogenesis by showing evolutionarily conserved decreases in X-linked transcription bursting kinetics in all examined somatic and germline cell types. Finally, we provide evidence supporting the cultivator model of *de novo* gene origination by demonstrating how the appearance of newly evolved testis-specific transcripts potentially provides short-range regulation of neighboring genes’ transcriptional bursting properties during key stages of spermatogenesis.

**Significance Statement:** Understanding the divergence and evolution of novel genes and expression is an essential question in evolutionary biology. However, rapid transcriptomic divergence in tissues such as testis has significantly hindered comparative single-cell RNA-Seq studies and understanding of the rapidly evolving components of the cell types. Here, we provide a novel strategy that does not rely on preexisting marker genes for cell type identity to overcome the challenge. We found that genome organization affects expressional evolution through transcriptional bursting dynamics. Our results also support the cultivator model of de novo gene origination—a model emphasizing the genomic environment in shaping novel gene origination—by illustrating how newly evolved transcripts alter the transcriptional bursting kinetics of neighboring genes.

## Introduction

Gametogenesis is an essential process in all sexually reproducing species in nature, ensuring genetic material is successfully transferred between generations. Male gametogenesis, spermatogenesis, is a specialized developmental process that exhibits several key features. First, cell differentiation from stem cell to fully formed gamete is remarkably conserved across animals. Second, despite the necessary maintenance of gametic function, testis-expressed genes tend to evolve very rapidly, with some genes being newly originated. Third, although cell types in the testis exhibit conserved features, expression patterns of genes in each cell type vary drastically between species. Because of this, the expression dynamics of testis-expressed genes are an important issue for both evolutionary biology and developmental biology.

While the transcriptomes of highly conserved cell types in testis are likely to remain stable over large evolutionary timescales, the rapid evolution of spermatogenesis in *Drosophila* presents significant challenges for comparative analyses of single-cell transcriptomic data, even among closely related species. For instance, many seemingly conserved marker genes from *D. melanogaster* (1, 2) are lost or not annotated in other *Drosophila* species, which means even marker genes, are not evolutionarily conserved. Additionally, the rate of gene expression changes drastically varies in *Drosophila* (3, 4), making it difficult to transfer cell types across different species. Our interest lies not only in the conserved transcriptional programs of cell types across species but also in complete gene expression patterns, including species-specific genes. The application of standard techniques (5–8) often fails to reproduce previous cell-type assignments in *D. melanogaster* samples, generating a large number of species-specific cell type clusters that likely do not reflect true phenotypic differences.

Within approximately 30 million years of evolution in *melanogaster* species group, the process of germline differentiation has undergone many phenotypic changes (9–11). Alternatively, a large degree of spermatogenesis (Figure 1A) has remained unaltered through similar evolutionary time scales (12, 13), including gross anatomical phenotypes such as organ structure or the progression of various cell-type specific changes (14, 15). We therefore hypothesized that a high degree of species-specific gene expression evolution within each cell type is the primary factor driving technical difficulties in clustering and identifying cell type differences within comparative studies of *Drosophila* spermatogenesis (Figure S1).

**Figure 1:**
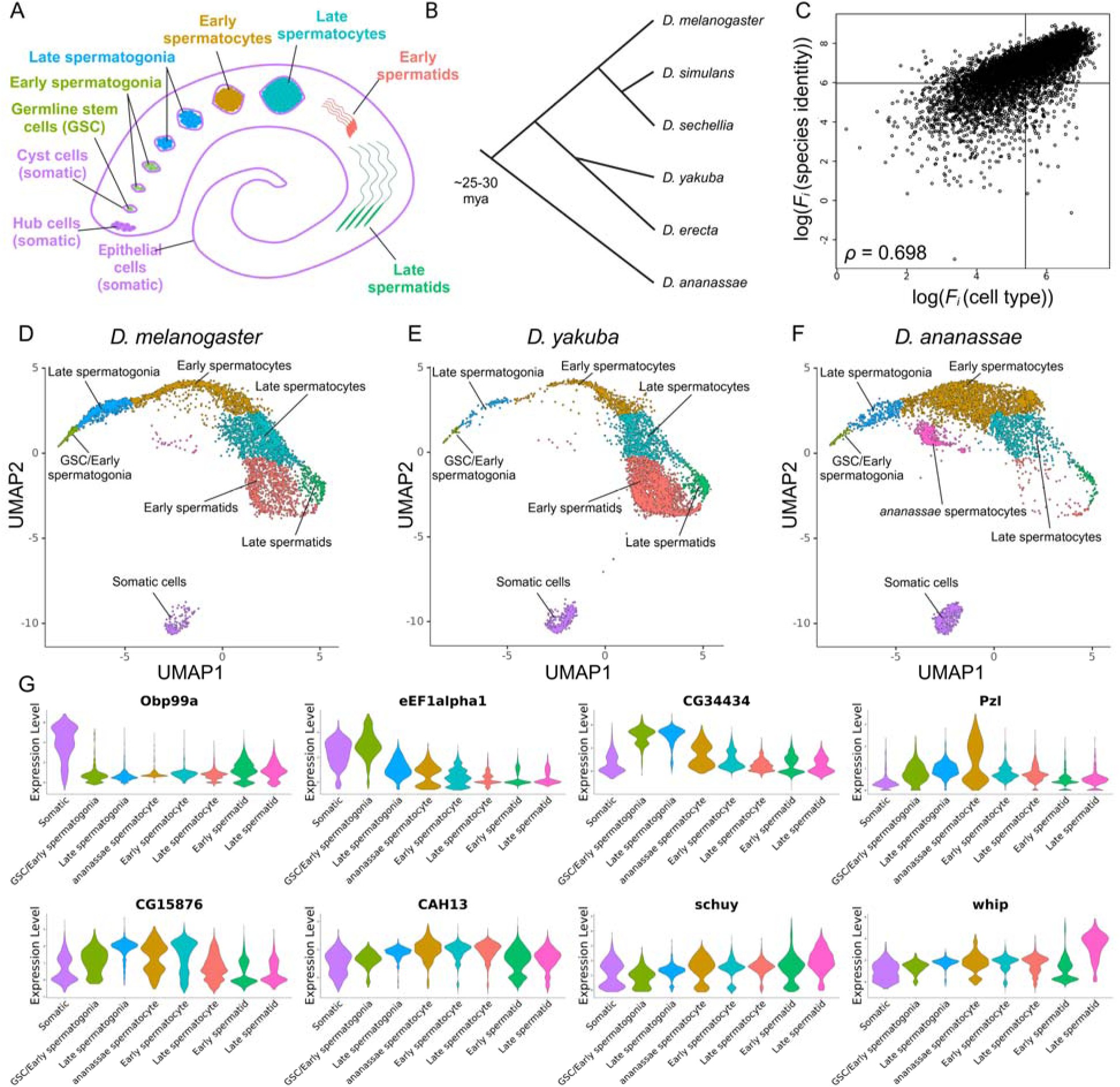
Identification of 198-gene list allows for cell type classification across species in a rapidly diverging process. **(A)** Illustration of spermatogenesis from germline stem cells to late spermatids. **(B)** Phylogeny of species used in this study with estimated divergence times. **(C)** ANOVA log(F-statistic) values from scRNA-Seq data using previously identified *D. melanogaster* cell types and cross-species labels for the 3 species show high correlation resulting in a large degree of species-specific cell type divergence. Drawn lines correspond to median *F*- statistic values for each (independent) marginal distribution. **(D-F)** UMAP projections resulting from the 198-gene list reveal a progression of intermediate cell types spanning germline stem cells to late spermatids for the 3 species. **(G)** Violin plots from the most differentially expressed gene within each cell type.

Transcription is an intrinsically random process which may be modeled stochastically (16, 17). Single-molecule observations of transcriptional activity are intrinsically “bursty,” consisting of short bouts of transcription that produce discrete quantities of mRNAs (“on” state) followed by periods of inactivity (“off” state) (18–20). Two key parameters controlling the transcriptional kinetics of different genes are burst size (or burst amplitude), which reflects the number of transcripts produced per burst, and burst frequency, which reflects the number of bursts that occur per unit time (18, 21, 22). Importantly, the analysis of transcriptional bursting kinetics offers unique insight into understanding how certain genes are regulated. The kinetics underlying burst size is a property of core promoter elements, driven by the stoichiometry of RNA polymerase II (pol II) availability, while the kinetics of burst frequency is influenced primarily by enhancer elements, modulated by the probability of transcriptional initiation (23–26). Transcriptional bursting properties have provided a novel perspective into the molecular underpinnings of sex-biased transcription burst amplitude in male and female embryos of *D. melanogaster* (27). Recent work has shown that the upregulation of male X chromosome genes is primarily mediated by a higher pol II initiation rate and burst amplitude (27), paving the way for further studies of dosage compensation in spermatogenesis.

Despite this, the molecular mechanisms underlying down-regulation of the X chromosome remain under-studied (28). While prior work has suggested that expression of genes on the X chromosome is globally down-regulated during spermatogenesis in *Drosophila* (29), it is not possible to determine anything beyond the general reduction of expression for X-linked genes (28, 30). Alternatively, previous single-cell RNA-Seq (scRNA-Seq) data has proposed that there is some degree of non-canonical dosage compensation (31, 32). A decrease in X chromosome expression beyond that which would be expected by dosage has also led others to suggest that the entire *Drosophila* X chromosome is inactivated (33).

Beyond the X chromosome, our interest in the testis is also related to the observation that the testis is a hotspot for tissue-specific expression of newly evolved genes (34, 35), including novel genes and transcripts (e.g., non-coding RNAs) arising *de novo* from non-genic, transcriptionally inactive sequences. While the origination of *de novo* protein-coding genes has been relatively well-studied (36–38), the study of the evolutionary mechanisms underlying the origination of *de novo* transcripts such as lncRNA in fruit flies has been lacking (36, 39, 40). This gap highlights the importance of comprehensively identifying these RNA genes in multiple *Drosophila* species(41). Such *de novo* transcripts are often expressed in a tissue-specific manner, typically in the adult testis of *Drosophila* (34, 36, 39). Intriguingly, scRNA-Seq data has shown that *de novo* originated genes, unlike recently evolved duplicate genes, are biased toward expression in mid-spermatogenesis (35). It is not yet clear if this pattern is common across multiple *Drosophila* species and what function these transcripts may hold.

In our study, we undertook a comprehensive analysis of spermatogenesis in three *Drosophila* species, aiming to unravel the complex transcriptional dynamics and novel expression evolution that underpin it. We first developed a novel approach to study scRNA-Seq data from rapidly evolutionarily diverging species. We then quantified a strong correlation between species- and cell-type-specific transcriptomic divergence across these species, identifying a core set of genes crucial for predicting cell type identity over approximately 30 million years of evolutionary divergence. After classifying cell types across these species, we analyzed how transcriptional divergence progresses throughout spermatogenic development. We further investigated the effects of genome organization on the molecular evolution of spermatogenesis, focusing on transcriptional bursting. We demonstrated variations in transcriptional burst size and frequency before and after meiosis, examining differences in autosomal versus X chromosomal transcription. Our findings support the cultivator model of *de novo* gene origination (42), showing that newly evolved testis-specific transcripts might play a regulatory role in the transcriptional bursting properties of neighboring genes.

## Results

### A simple ANOVA-based statistical procedure identifies gene lists predictive of cell type but not species identity

We used several commonly available software suites to analyze scRNA-Seq data from testes tissue derived from three species, *D. melanogaster*, *D. yakuba*, and *D. ananassae*, but were unable to obtain high-quality cell type assignments to correlate similar cell types across different species (Figure S1). For example, widely used markers in *D. melanogaster* did not robustly classify cell type identities in species such as *D. ananassae*. We thus hypothesized that species-specific cell type transcriptomic divergence significantly hinders cell type classification. Clear morphological and functional conservation of spermatogenesis across various *Drosophila* species, e.g., mitotic staging or spermatid differentiation, suggests some degree of evolutionary conservation of key genetic elements. In this case, challenges in cross-species clustering could potentially be overcome by focusing on key conserved elements, not the entire transcriptome. We thus developed a simple methodology to identify genes that are both evolutionarily conserved and important in determining cell identity. We hypothesized that these genes could be identified by calculating ANOVA *F*-statistic scores for cell type (where available) and species. To test this strategy, we generated a simulated data set consisting of nine unique cell types evolving in three different species (Supplementary Methods, Figure S2A), to simulate a scenario in which cell type differences are masked by species-level differences. (Figure S2A). A good cell type marker gene, indexed by variable *i*, for multiple species should exhibit a large *F_i_* for cell type and a small *F_i_*for species identity (Figure S2B, bottom-right quadrant). We selected the top 200 genes showing high *F_i_* (cell type) and low *F_i_* (species identity). The genes identified through our methodology effectively captured the most informative genes, preferentially containing cell type information over species identity information (Supplementary Methods, Figure S2C).

Uncorrected simulated data resulted in 27 non-overlapping cell types, as they were clustered based on both cell type *and* species identity (Figure S2D). When standard batch correction techniques, such as Monocle3’s implementation of batchelor via ‘align_cds()’ (6, 43), are applied to remove species-specific effects, UMAP and PCA projections show that these effects have been entirely removed with entirely overlapping cell type assignments despite species-specific differences in cell type (Figure S2E). This stands in contrast to the removal of species-specific changes using these same batch correction techniques in real scRNA-Seq data derived from testis tissue (Figure S1), as remaining species-specific differences hinder the identification of cell types across species in these data sets. This is likely due to the excessively simplistic and additive nature of the simulations in contrast to the more complex, non-linear nature of cell-type specific evolution. Regardless, the application of standard batch correction techniques remained insufficient for cell type identification in real data.

Projection and clustering using the top 200 gene list in the simulated data showed 9 distinct cell type clusters that still maintained species-specific differences (Figure S2F, G). Cell type identities are readily recovered, while species-specific divergence within these cell types is also clearly visualized. These show that our method identifies gene lists that are predictive of cell type but not species identity, which is important for clustering cell types across species. The methodology is also effective across phylogenies of varying degrees of asymmetry (Figure S3, Supplementary Methods: *Performance of H_SQ_ criteria in asymmetric phylogenies*).

### Application of the ANOVA-based statistical procedure identifies 198 gene list in related Drosophila species

We then applied our non-parametric statistical procedure on our scRNA-Seq data set derived from testis tissue in *D. melanogaster*, *D. yakuba*, and *D. ananassae*. The divergence time between *D. melanogaster* and *D. yakuba* has been estimated to be about 7 million years ago while the divergence time between *D. melanogaster* and *D. ananassae* has been estimated to be about 25-30 million years ago (44)(Figure 1B). Prior data and cell type labels from *D. melanogaster* were obtained from (35), while 5000 cells derived from *D. yakuba* and 5000 cells derived from *D. ananassae* were sequenced (Supplementary Methods). Raw data was aligned in ‘cellranger’ to reference genomes and annotations obtained from FlyBase to generate read counts. Further orthology information between the three species’ annotations was obtained using FlyBase (dmel-r6.44) and used to generate “melanogasterized” *D. yakuba* (dyak-r1.05) and *D. ananasssae* (dana-r1.06) genomes. Specifically, each genome was reduced to a core set of single-copy genes that possessed strict one-to-one orthology between the three species. After processing in ‘Monocle3’ (43), data from one-to-one orthologs were combined together and retained. A summary of key processing parameters is presented in Table S1.

The distributions of *F_i_* (cell type) and *F_i_*(species identity) were calculated using our experimentally derived data (Figure 1C, Data S1). Interestingly, the joint distribution of log(*F_i_* (cell type)) and log(*F_i_*(species identity)) showed an exceedingly large correlation (Pearson ρ = 0.698, p < 2×10^-16^, Figure 1C). This result is on par with the correlation found in simulated data (Figure S2B, Pearson ρ = 0.74, p < 2×10^-16^), suggesting a high degree of species-specific cell type evolution in *Drosophila* testis tissue. The medians of the independent distributions *F_i_* (cell type) and *F_i_* (species identity) were chosen as threshold *p_cell-type_* and *p_species-identity_*, respectively (Figure 1C, solid lines). Note that the independent distributions of *F_i_* (cell type) and *F_i_* (species identity) are not fully represented in the joint distribution of *F_i_* (cell type) and *F_i_*(species identity), because some genes were missing *F_i_* (cell type) or *F_i_* (species identity) values. This definition of *p_cell-type_*and *p_species-identity_* produced a new 198-gene (Table S2).

To support the biological significance of the 198-gene set, we performed a gene ontology (GO) analysis using the PANTHER database (45, 46)(Table S3). The eleven statistically significant major categories (FDR < 0.05) were highly concordant with known function during spermatogenesis, including ‘cilium mobility,’ ‘dynein-associated pathways,’ ‘insulin signaling,’ ‘centrosome cycle,’ ‘protein maturation,’ ‘proteolysis,’ and ‘general reproduction.’ Notably, all of these major categories showed over-representation in comparison to what is expected for all genes in *D. melanogaster*, demonstrating that this statistical procedure effectively identifies transcriptionally important, functionally conserved genes in the rapidly evolving *Drosophila* testis transcriptome.

### Cell type clustering of testis scRNA-Seq data reveals increasing transcriptomic divergence throughout spermatogenesis

To validate the expression of the 198-gene set we identified, we visualized both UMAP and PCA projections of all analyzed cells with an overlay of previously identified cell type assignments (35)(Figure S4A). Importantly, cells from all species can be projected onto the same visualization (subsetted by the 198-gene list) without extraneous processing beyond standard single-cell transcriptomics pipelines (Figure 1D-F, Figure S4B). We note that batch correction for species was applied to the simulated 200-gene and to this 198-gene list as part of Monocle3’s standard pipeline. A visual examination of these projections shows striking features concordant with known spermatogenesis function. 1) While the sequence of spermatogenesis was not explicitly encoded into our statistical procedure or in the visualization process, the developmental progression of cell types in the projection follows a known progression from germline stem cells to late spermatids. Despite being a continuous developmental process where each stage does not produce spatially segregated clusters, the stages of previous cell type assignments form relatively distinct boundaries. This eliminates the need for more advanced pseudo-time projection techniques. 2) All somatic cell types cluster separately from germ cell types. 3) We observe a gradual divergence in gene expression between cells of the same cell type as sperm mature. For example, germline stem cells appear to have low expression variance across all species, while spermatids show much larger variance (Figure 1D-F). Note that all these features are readily observed in PCA projections of the same data set (Figure S4B).

Importantly, the application of this methodology allows for recapitulation of previously identified cell type clusters using unsupervised clustering on a unified, pan-species UMAP projection with the clustering resolution set to produce the same number of clusters reported in previous studies (35) (Leiden clustering, resolution = 3 x 10^-4^, 9 clusters). While it is likely to be the result of residual species-specific differences, these cell type assignments show that fewer cells are represented in the earlier germline stem cell, spermatogonia, and spermatocyte stages of *D. yakuba* when compared to *D. melanogaster.* We also observe a correlated accumulation of cells in later spermatid developmental stages. These cell type assignments also reveal the presence of an additional cell cluster/expression present in *D. ananassae* that appears to be absent from *D. melanogaster* and *D. yakuba*. We believe that this cluster is likely a result of transcriptomic divergence (over evolutionary timescales) that persists even after batch correction. Specifically, these cells likely represent a set of spermatocyte cells that demonstrate a stereotypical expression pattern that exists only in *D. ananassae* (see *New spermatocyte cell cluster reveals a unique expression pattern* and Supplementary Materials: *‘ananassae’ cell cluster*). We also observe a low abundance of early and late spermatid stages in *D. ananassae* (Figure 1D-F, Figure S4B). These changes in cell type assignment and visualized patterns among species reflect changes in the expression patterns of the 198-gene list, possible technical biases during data generation, and fundamental processes driving species-specific differences in reproduction (see “*New spermatocyte cell cluster reveals a unique expression pattern*”, Supplementary Materials: *‘ananassae’ cell cluster*). These cell clusters were further investigated using RNA FISH (*RNA in-situ hybridization chain reaction validates cross-species cell type assignments*.)

### High differential expression across cell types in D. melanogaster spermatogenesis does not necessarily indicate differential expression across cell types in closely related species

A cross-species analysis of the five most significantly differentially expressed genes for each cell type/cluster was performed. Unless specifically noted, the phrase “differential expression” indicates analyses of differential expression across cell types that do not take species identity into account. For example, if a gene is said to be differentially expressed in GSC/Early spermatogonia, we mean that the expression of this gene is altered when comparing GSC/Early spermatogonia *for all species* to all other cell types *for all species*. This is achieved by combining GSC/Early spermatogonia from *D. melanogaster*, *D. yakuba*, and *D. ananassae* into one category and comparing these cells to all other cells detected in *D. melanogaster*, *D. yakuba*, and *D. ananassae*.

Interestingly, only one (*MtnA*) of thirteen previously reported markers (*aly*, *zfh1*, *Fas1*, *Hsp23*, *MtnA*, *His2Av*, *aub*, *bam*, *fzo*, *twe*, *soti*, *Dpy-30L2*, *p-cup*) (35) remains as one of the top- 5 differentially expressed genes by cell type (all spp., e.g. all GSC from all spp. considered together, Figure S5). These results suggest that while the 13 markers genes identified in previous *D. melanogaster* studies may be highly differentially expressed across cell types in *D. melanogaster*, these expression patterns are likely to have evolved in a species-biased manner. A lack of one-to-one orthology may also be excluded as a driving factor in these discrepancies, as only one of these thirteen markers (*zfh1*) was removed due to this reason.

We then analyzed all significantly differentially expressed genes for each cell type, testing for both over- and under-expression (Data S2, Table S4). This analysis was performed by comparing cell types without regard to species. Among the twelve previously used markers present in the data set, *Fas1*, *fzo*, and *Dpy-30L2* were not found to be differentially expressed in any of these cell types (all spp.). In the somatic cells, *Hsp23* and *MtnA* remained highly differentially expressed (all spp.) with the correct cell type specificity. Similarly, early spermatogenesis markers did appear to be differentially expressed in GSC/Early spermatogonia cell types, but with highly attenuated signals. For example, of all 1326 significantly differentially expressed genes in GSC/Early spermatogonia, the most differentially expressed classic marker gene was *His2Av*, which was ranked 375 when sorted by log(fold change), while the next closest marker was *aub*, ranked 600. This attenuation of signal appears to continue through later stages of spermatogenesis, but reverses at the late spermatid stage. There, *p-cup* is ranked 18/639 and *soti* is ranked 24/639 respectively. Interestingly, cells demonstrating the “*ananassae*-specific” expression pattern showed significant overexpression of both early and late spermatogenesis markers: *His2Av* (ranked 245/366) and *p-cup* (ranked 73/366), respectively. The attenuation of cell-type information of previously utilized marker genes highlights how cell type-specific evolutionary divergence contributed to the masking of proper cell type assignment between species.

Among the twelve classic marker genes in our data set, only *aly* appears in our 198-gene list, indicating that the remaining *D. melanogaster* marker genes’ ability to predict cell type likely occurs in a species-specific manner. Eight of the twelve marker genes did not appear in the 198-gene list as they showed high levels of species information, while the remaining four marker genes remained relatively similar across species. Of those, two genes (*aub*, *bam*) were excluded for having undefined *F_i_* (cell type) values resulting from little to no variance in certain cell types. This low/no variance occurs due to having uniformly few to no counts within different cell types. Interestingly, the remaining two genes (*Hsp23*, *MtnA*) were not excluded for having evolutionarily diverged expression patterns, but they were excluded for having poor predictive ability for cell type. This appears to contradict the appearance of *MtnA* in an analysis of the most highly differentially expressed genes across cell types (regardless of species, see Data S2). This discrepancy arises from differences in statistical methodologies. *MtnA* expression is found only in a very small number of cells, primarily somatic cells. Differential expression analysis, which compares expression within one cell type vs expression across all other cell types (“one-vs-all”), revealed elevated *MtnA* expression in somatic cells (all spp.) despite its low detection in many other cell types (all spp.). In contrast, ANOVA compares the between-cell type variance across all cell types, where *MtnA*’s limited detection results in a low *F*-statistic value due to minimal variance in most cell types. This highlights the value of performing cell type assignment on genes that show highly specific yet robust expression across cell types. This effect is particularly important in the context of scRNA-Seq technology, where the absence of detection may be as much the result of low read count as it is of biologically low expression. Focusing on quantitative differences within widely expressed genes like those in our 198-gene list, rather than a reliance on non-detection events of relatively lowly expressed genes, may lead to more consistent cell type classification in future comparative scRNA-Seq studies.

### RNA in-situ hybridization chain reaction validates cross-species cell type assignments

While we were able to transfer a reference species’ cell type labels across species both in our simulated data sets and in our three-species data set, it was unclear whether the cell type assignments reflected biological reality. To determine the accuracy of the cell type assignments, we performed *in-situ* hybridization chain reactions (HCR RNA FISH) (47, 48) in whole-mount testis tissue from *D. melanogaster* and *D. ananassae*. Four genes were chosen: *Rbp4*, *B52*, *Pkd2*, and *soti* (Figure 2) to fulfill multiple criteria: 1) highlighting predicted evolutionary divergence between species in common marker genes, 2) demonstrating high cell-type specificity to aid interpretation of subsequent *in-situ* imaging, and 3) providing maximal information regarding our newly detected “*ananassae* spermatocyte” cluster. We note that during imaging, there was a large degree of crosstalk between the fluorophores representing *B52* and *soti*, so images were not amplified using both fluorophores in the same sample.

**Figure 2:**
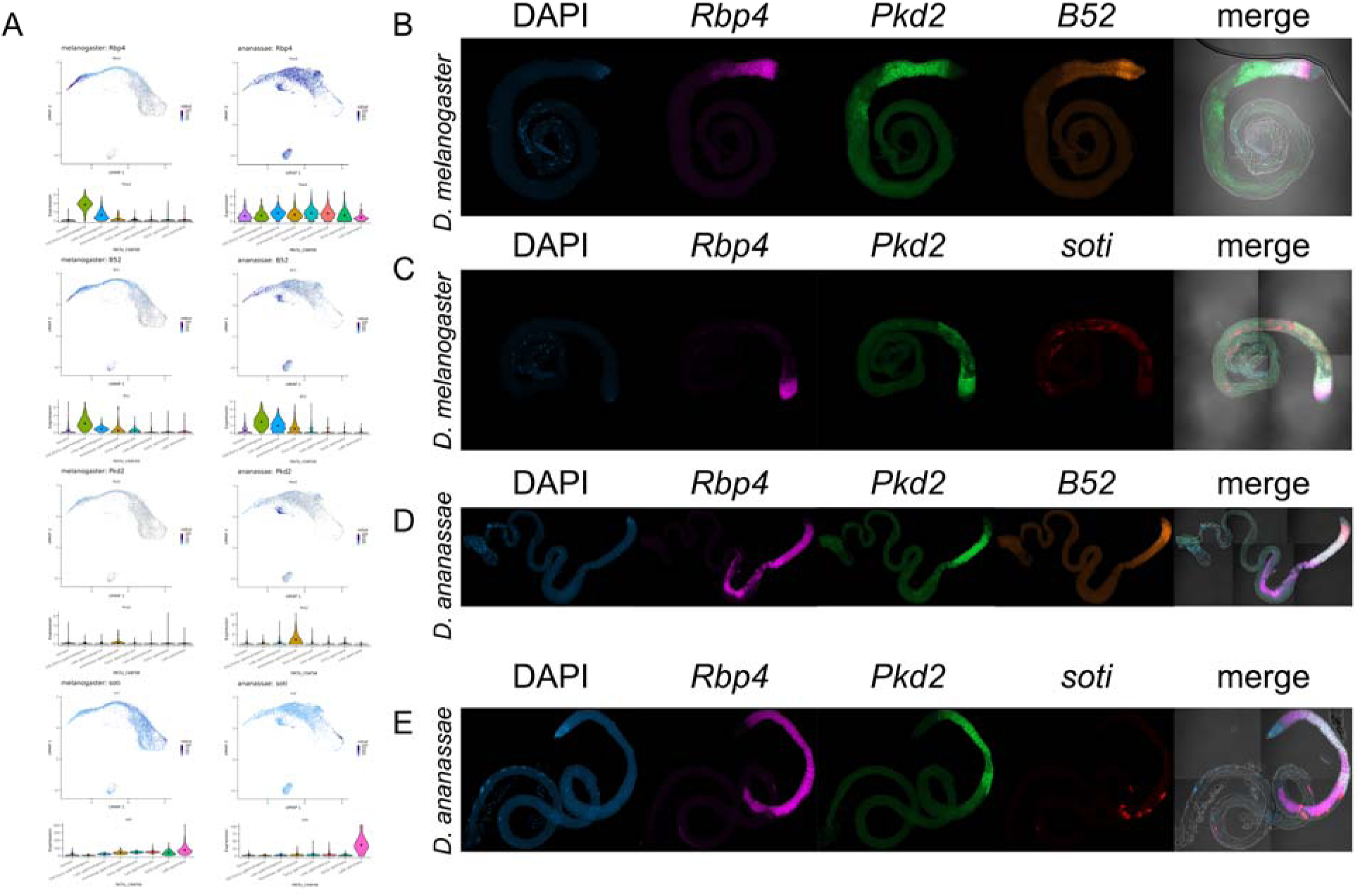
Validation of expression pattern divergence and cell type classification using HCR RNA FISH. **(A)** scRNA-Seq UMAP shows predicted expression patterns in *D. melanogaster* and *D. ananassae* for four genes: *Rbp4*, *B52*, *Pkd2*, and *soti*. **(B-E)** RNA FISH images in *D. melanogaster* **(B, C)** and *D. ananassae* **(D, E)** using these same genes, with DAPI nuclear staining (**B-E**, left column) and overlay of these channels over brightfield microscopy (**B-E**, right column). *Rbp4* is a marker gene used in *D. melanogaster* to indicate germline stem cells/spermatogonia. Our scRNA-Seq cell type classification predicts that *Rbp4* has a broad expression pattern in *D. ananassae*. *B52* is a gene with a restricted expression pattern in *D. melanogaster* germline stem cells/spermatogonia. This expression pattern is predicted to be slightly broader in *D. ananassae*, extending into the new *ananassae* spermatocyte cluster. *Pkd2* is predicted to be primarily restricted to the newly identified *D. ananassae* spermatocyte cluster in both *D. melanogaster* and *D. ananassae*. *soti* is a marker gene used in *D. melanogaster* to indicate spermatid cell types, and our classification predicts that *soti* has both higher overall expression levels and lower cell type specificity in *D. melanogaster* than in *D. ananassae*.

*Rpb4* was chosen as it was a previously employed cell-type marker for *D. melanogaster* GSC/Early spermatogonia cell types in prior publications (1). Our cross-species cell type classification predicted that this marker gene had highly differing expression patterns between *D. melanogaster* and *D. ananassae* (Figure 2A). This prediction was validated, as *Rbp4* showed an unexpectedly wide expression pattern in *D. ananassae* beginning with early spermatogonia and culminating in expression overlap with late spermatids (*soti* expression and DAPI staining) (Figure 2B-E). This is a surprising finding, as *Rpb4*, an RNA-binding protein, was found to be a regulator of mitosis. *Rbp4* interacts with *Fest* to regulate the translation of *Cyclin B* (49). Similarly, the genes *Lut* and *Syp* interact with *Fest*, regulating post-transcriptional control of *Cyclin B* (50). We then investigated the cell type-specific expression of the genes *Cyclin B*, *Fest*, and *Syp* in our data set (Figure S6). *Lut* was excluded as it was not present as a one-to-one ortholog in our data set. The expression pattern of *Cyclin B* agrees with previously reported expression patterns in *D. melanogaster* (49), while also adopting a similarly specific pattern in *D. yakuba*. Interestingly, the expression pattern of *Cyclin B* appears to be non-specific in *D. ananassae*, showing a broad expression across all of spermatogenesis like *Rbp4*. While the *D. melanogaster* expression patterns of *Fest* and *Syp* were less specific than for *Rbp4* and *Cyclin B*, this low specificity decreased even further in *D. ananassae* (Figure S6). This result indicates that changes in cell type biased patterns of genes potentially reflect underlying functional and molecular changes.

Finally, *soti* was chosen as it was previously employed as a cell-type marker for *D. melanogaster* late spermatid cell types (1, 2, 31, 35). Our cross-species cell type classification predicted that transcript levels in *D. melanogaster* would both be higher and less specific than in *D. ananassae*. This prediction was validated using our *in-situ* imaging. For example, *D. melanogaster soti* expression appears to be broader than in *D. ananassae*, forming a broader, “smearier” expression pattern when compared to the tighter, more individualized puncta seen in *D. ananassae* (Figure 2). Note that these figures show maximal intensity projections on z-stacks.

### New spermatocyte cell cluster reveals a unique expression pattern

A striking result of the previous sections is the identification of a *D. ananassae*-specific cluster that appears in our cross-species UMAP clustering. We thus use our scRNA-Seq and RNA FISH data to further investigate this newly identified cluster. As mentioned in the previous section, this “*ananassae*-specific” cluster appears in the UMAP of *D. ananassae* cells but does not seem well-represented in *D. melanogaster* and *D. yakuba* projections. While the appearance of this cluster seems correlated with the inferred disappearance of spermatid cell types in *D. ananassae*, the new cluster also appears in close proximity with other known spermatocyte cell types in our unified cross-species projections (Figure 1D-F).

To resolve ambiguity and to further investigate the specificity of this new cluster, we identified genes that are differentially expressed (all spp.) in this cluster in comparison to all other clusters (see “*High differential expression across cell types in D. melanogaster spermatogenesis does not necessarily indicate differential expression across cell types in closely related species*”). The 20 most highly differentially expressed genes in the “*ananassae* spermatocyte” cluster are reported in Table S5.

While many of these differentially expressed genes are predicted to have expression in more than one tissue type, *Pkd2*, the 6^th^ most differentially expressed gene in this cluster, was also found to have relatively specific expression. In particular, *Pkd2* demonstrated elevated cluster-specific expression in *D. ananassae*. While specific, we note that *Pkd2* is still predicted to have residual expression from the GSC/Early spermatogenesis stage to the Early spermatid stages in both *D. melanogaster* as well as *D. ananassae* (Figure 2A).

A detailed examination of the RNA FISH data for *Pkd2* reveals surprisingly broad expression throughout spermatogenesis (Figure 2B-E), particularly in contrast to *B52* expression. *B52* was chosen because it is predicted to express in early spermatogonia as well as in our newly identified *ananassae*-specific cluster. As predicted, *B52* showed high specificity in germline stem cells and spermatogonia, with a residual expression detected in our “*ananassae* spermatocyte” cell cluster. These results confirm that cells with the “*ananassae* spermatocyte” expression pattern are a type of spermatocyte.

Expression of *Pkd2* in *D. melanogaster* and *D. ananassae* was found to overlap with GSC/Early spermatogonia as indicated by the expression pattern of *B52* (both species) and *Rbp4* (*D. melanogaster* only, “*Rbp4-mel*”). The expression of *Pkd2* subsequently decreases in concert with *B52* and *Rbp4*-*mel* as we continue along the germline differentiation process. Interestingly, *Pkd2* remains relatively highly expressed even in late-stage spermatocytes and onion-stage spermatids preceding spermatid elongation. This late-stage expression is most interestingly highlighted by the occasional presence of round cells that express *Pkd2* well into the spermatid stages of differentiation as represented by *soti* expression (Figure 2B, E). This combination of scRNA-Seq and RNA FISH data suggests that the “*ananassae* spermatocyte” cell cluster has a biological component and is not the sole result of technical artefacts. However, we stress that this cluster does not likely represent a cell type that is functionally different than previously identified spermatocytes, but rather a shift in the expression pattern of a set of these cells. This is further discussed in the Supplementary Materials section ‘*ananassae’ spermatocyte cluster*.

### Transcriptional bursting in spermatogenesis is controlled via opposing mechanisms during meiotic transition

We used our validated cell type assignments to study the evolution of transcriptional control by examining how the transcriptional bursting properties of different genes change during development. General transcriptional activity first increases, then dramatically decreases during the stage of spermatogenesis between the 16-spermatocyte stage and the second round of meiosis. These divisions subsequently result in 64 round spermatids in all three species (11). Importantly, the total pool of remaining transcripts is slowly degraded as they are translated in subsequent stages, supplemented only by a small number of genes that are transcribed post-meiotically. The increased resolution of scRNA-Seq analysis (26, 51) and MS2-MS2 coat protein system imaging (24, 25) has brought significant advances in understanding the fundamental molecular mechanisms underlying transcription, shedding light on the chemical kinetics that drive stochastic gene expression. However, global control of the pre- to post-meiotic transition during *Drosophila* spermatogenesis has remained understudied.

Aside from higher-order eukaryotic regulation via nucleosomes and histones, sequence-based transcriptional control is exerted by genomic segments that are broadly classified as promoter and enhancer sequences. Prior results have mechanistically demonstrated how this transcriptional control of promoters alters burst size via pol II availability, while control of enhancers mechanistically alters burst frequency via BRD4 and the Mediator complex ((20, 26, 52–54)). To investigate how transcriptional control of germline differentiation is controlled in *Drosophila*, we begin by calculating burst size and frequency for all annotated genes by cell type using ‘txburst,’ using our three-species data (Figure 3)(26). After applying default txburst quality control criteria, the number of genes that were analyzed per cell type varied by cell type and species (Figure S7, Table S6). Interestingly, we find that in the early, pre-meiotic stages of spermatogenesis (i.e., germline stem cells, early spermatogonia, late spermatogonia, and early spermatocytes), transcriptional bursting is primarily controlled via alterations in burst size. While burst frequency also increases during these stages, overall burst size appears to reach a maximum during the late spermatogonia stage before declining during spermatocyte stages (Figure 3). Interestingly, in *D. melanogaster*, burst size drops sharply between the early and late spermatocyte stages, while in *D. yakuba* and *D. ananassae*, burst size decreases between the spermatogonia and spermatocyte stages, remaining relatively stable from early to late spermatocyte stages (Figure 3A-F). This genome-wide transition from burst size-dominant control (reflecting promoter-based regulation) to burst frequency-dominated control (reflecting enhancer-based regulation) suggests that phenotypic divergence during spermatogenesis in *Drosophila* may be evolving via global, genome-level alterations of transcriptional control via pol II initiation (‘k_ini_’) in alignment with prior observations (20, 52, 53, 55), rather than through alterations in a smaller subset of genes.

**Figure 3:**
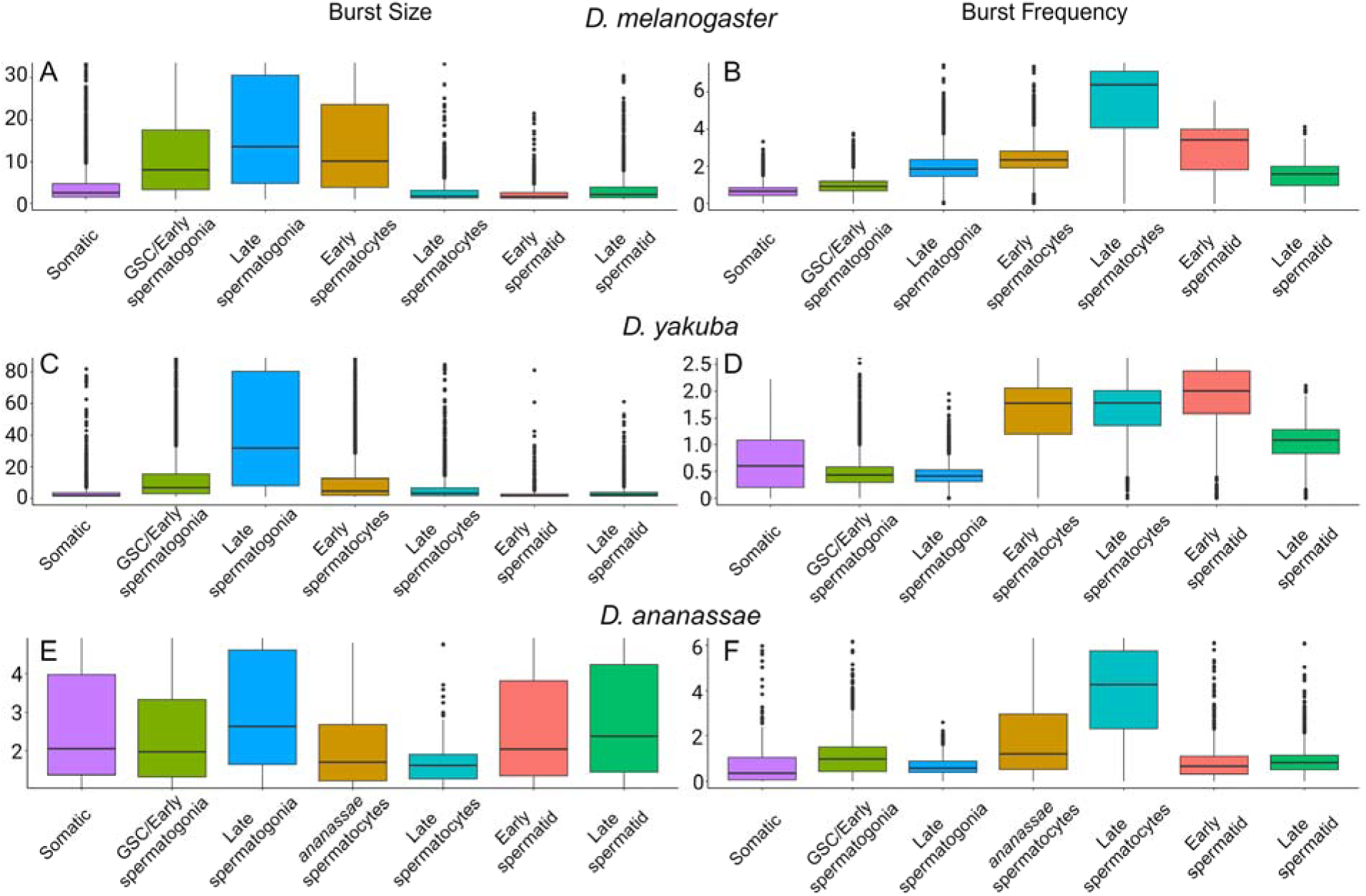
Pre- and post-meiotic cell types in spermatogenesis are controlled by opposing mechanisms. The burst size and burst frequency distributions for **(A, B)** *D. melanogaster*, **(C, D)** *D. yakuba*, and **(E, F)** *D. ananassae* reveal that transcription in the earlier stages of spermatogenesis are primarily controlled via burst size, while transcription in later stages of spermatogenesis is primarily controlled via burst frequency. This transition between control mechanisms correlates with the transition between mitosis and meiosis.

The control of bursting frequency appears to become the more dominant mechanism during post-meiotic stages of spermatogenesis (i.e., late spermatocyte, and spermatid stages, Figure 3B). An increase in bursting frequency in *D. melanogaster* is observed in the late spermatocyte stage simultaneously with a correlated decrease in burst size. A similar change is observed between late spermatogonia and early spermatocyte in *D. yakuba*, where a decrease in burst size occurs simultaneously with an increase in bursting frequency (Figure 3C, D). In *D. ananassae*, an increase in burst frequency is observed in the early spermatocyte stage followed by an even larger increase in the late spermatocyte stage (Figure 3E, F). While the increase in burst frequency during the transition between late spermatogonia and early spermatocyte stages is correlated with a decrease in burst size in the same stages, the second increase of burst frequency during the late spermatocyte stage does not show a correlated change in burst size. This increase in burst frequency is potentially an outlier, as not only did a very low number of annotated genes pass quality control measures (202 genes, Figure S7C), but also the variance in the observed distribution is also quite high. These observations could also reflect a deeper evolutionary divergence in transcriptomic control or cell type, resulting from a weaker cell type assignment as revealed by its sparse representation in UMAP and PCA projections (Figure 1E). If the inferred distribution is closer in value to distributions found for the early spermatocytes and *D. ananassae* spermatocyte cell types, it would appear that *D. melanogaster* may have undergone a species-specific alteration where the size-to-frequency control transition has been delayed from the early spermatocyte stage to the late spermatocyte stage.

Overall, these results are consistent with a model of transcriptional control where an abundance of pol II is present (20, 26, 52, 53, 55) during earlier stages of spermatogenesis and becomes depleted during the meiotic transition. During the period of high overall pol II availability and activity, general levels of transcription remain high (20, 26, 52, 53, 55), and bursting kinetics are dominated by large burst size. Subsequent to this depletion, transcription then decreases significantly, shifting global transcriptional control into a more burst frequency-dominated mode of activity. It is likely that this modulation of burst frequency is achieved by enhancer-related changes in transcription factor stoichiometry (23, 25, 56).

### Bursting kinetics of X-linked vs autosomal genes reveals that partial dosage compensation likely occurs in somatic and germline tissue

To help elucidate the timing of dosage compensation during spermatogenesis, we examined the transcriptional bursting kinetics for both X-linked (N=2198) and autosomal (N=11749) protein-coding genes. Interestingly, our data set shows a highly consistent pattern where transcriptional burst size and frequency are significantly lower for X-linked genes than for autosomal genes across cell types and species (Mann-Whitney U test, Benjamini-Hochberg correction, FDR < 0.05) (Figure 4). Significant differences between X-linked and autosomal bursting kinetics remain consistent throughout spermatogenesis. Perhaps most surprisingly, this difference is detected in *D. melanogaster* somatic tissue. While X-linked genes typically show lower activity than autosomal genes (even in statistically insignificant comparisons), the transcriptional bursting frequency for X-linked genes is higher than for autosomal genes in *D. yakuba* early spermatocytes. Additionally, while nearly every cell type in *D. melanogaster* and *D. yakuba* produces a significant difference between X- and autosomal-linked genes’ transcriptional kinetics, the observed differences in *D. ananassae* are typically not significant after multiple hypothesis correction. This is likely because the number of genes that pass quality control in txburst is much lower for *D. ananassae* than for *D. melanogaster* and *D. yakuba*, resulting from the relatively poor reference genome quality and read depth for *D. ananassae* vs *D. melanogaster* and *D. yakuba* (Table S1).

**Figure 4:**
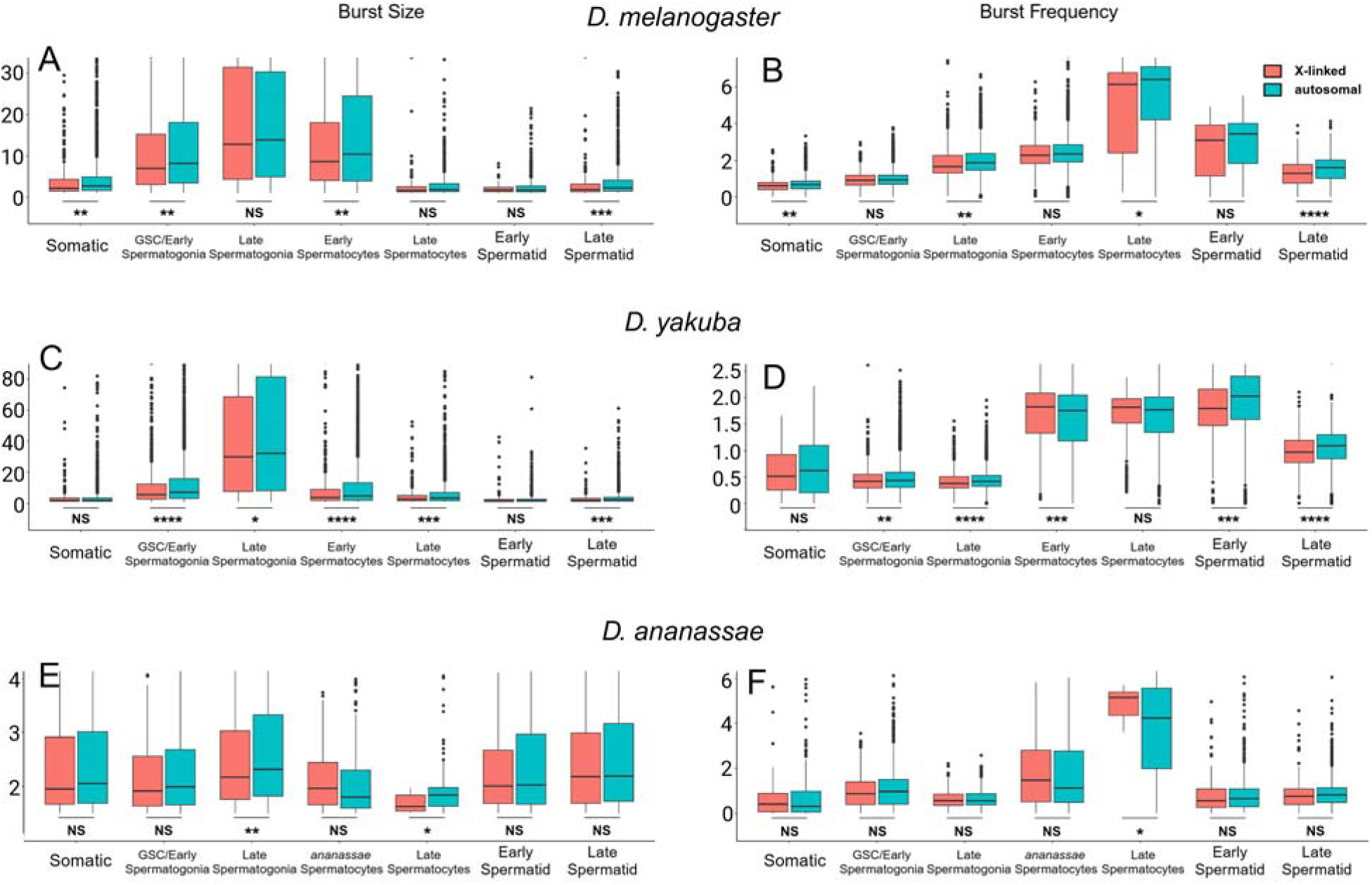
X-linked genes show lower bursting kinetics than autosomal genes prior to spermatogenesis. The burst size and burst frequency distributions for **(A, B)** *D. melanogaster*, **(C, D)** *D. yakuba*, and **(E, F)** *D. ananassae* reveal a conserved pattern of smaller average burst sizes and lower bursting frequencies for X-linked than autosomal genes across cell types (* = p < 0.05, ** = p < 0.01, *** = p < 0.001, **** = p < 0.0001. Note these are p-values before Benjamini-Hochberg correction, FDR < 0.05). No stage of spermatogenesis shows lower burst kinetics for X-linked genes than autosomal genes. The significant repression of X-linked genes in somatic cells suggests that decreased meiotic X-chromosome expression occurs in a developmental stage preceding testis maturation.

Despite these differences, the ratio between X-linked and autosomal genes passing quality-control remains relatively constant across all stages and species (26) (Figure 4, Figure S7), reflecting both the number of genes that are expressed in each stage as well as read depth. The consistent ratio of X to autosomal genes expressed does not reveal a particular stage of spermatogenesis during which the entire X chromosome is entirely inactivated. After either the early spermatocyte stage in *D. melanogaster* or the late spermatogonia stage in *D. yakuba* and *D. ananassae*, there is a drop off in the number of genes that pass QC (Figure S7). Despite this change, the number of genes expressed on the X vs autosomal chromosomes remains at a relatively constant ratio. As this ratio remains constant even when comparing somatic cell types to germline cell types (Figure 4), it is possible that the process of X-downregulation occurs prior to the establishment of somatic vs. germline identity during the differentiation of testis tissue (57). This pattern could potentially be further tested by comparing the transcriptional bursting characteristics of X-linked to autosomal genes in earlier stages of testis organogenesis, e.g., in larval testis tissue.

### Origination of new transcripts alters regulation of neighboring genes by increasing burst size during meiotic transition

The cultivator model of *de novo* gene origination hypothesizes that the most evolvable method of altering the expression of pre-existing genes is the appearance of a new proximal promoter element (42). Unlike changes to existing transcription-factor binding sites, as enhancer sequences typically act on longer range distances (≥ ∼1.5 kb), the cultivator model posits that the appearance of a promoter would act on relatively short ranges (∼100-500 bp), likely through enhancer-mediated promoter interference (24) or supercoiling-mediated transcriptional coupling (58). One key prediction of this model would be that ncRNAs are found to be closer to their protein-coding neighbors than expected by chance – this is indeed the case (42). Another key prediction of this model is that the origination of a *de novo* transcript would alter the regulation of its immediate neighboring genes, but not more distant neighbors as might be expected by enhancers. To test this prediction, we examined the transcriptional bursting characteristics of protein-coding genes that are neighboring newly activated *de novo* transcripts.

We begin by identifying four neighboring protein-coding genes for all 170 transcripts identified through bulk RNA-Seq analysis (Supplementary Methods, Data S3), two upstream and two downstream (“L2,” “L1,” “R1,” and “R2” respectively, Figure 5). Note that the “L2” and “L1” genes (or “5’-2” and “5’-1”) are defined to be to the 5’ of the *de novo* transcript, regardless of the sense or anti-sense orientation of the transcript. Similarly, “R2” and “R1” genes (or “3’-1” and “3’-2”) are defined to be to the 3’ direction. Using these neighboring genes’ orthologous genes in *D. yakuba* and *D. ananassae*, we compared the burst size and frequency for each class of neighbors to genome-wide distributions. Unlike the 170 evolutionarily young *de novo* transcripts identified through our comparative analysis of bulk RNA-Seq data, the neighboring protein-coding genes are not expected to be biased in an evolutionary age.

**Figure 5:**
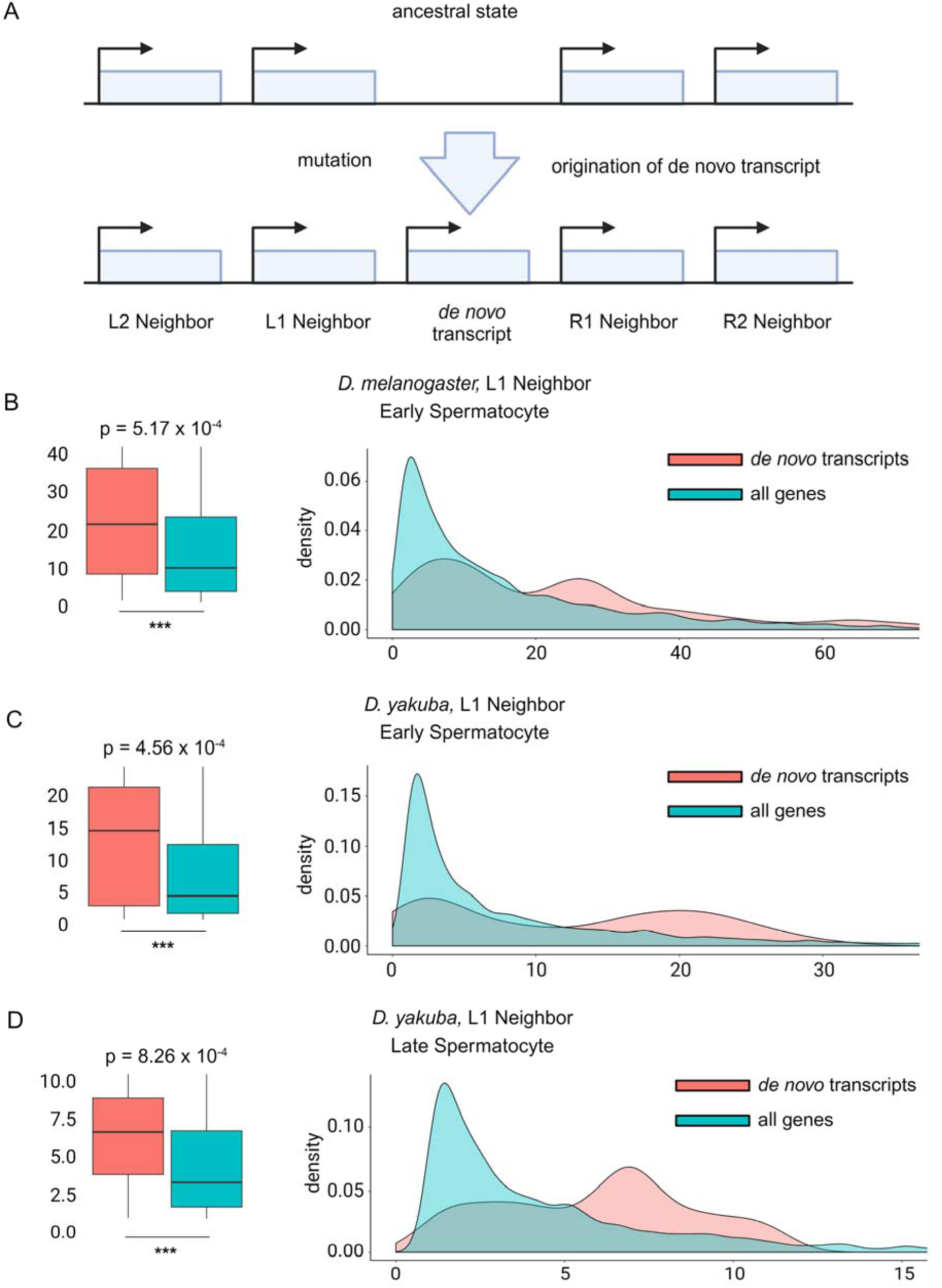
Higher burst sizes in L1 neighboring genes of *de novo* transcripts support a cultivator model of *de novo* gene origination. **(A)** The initial steps of the model suggests that a new promoter element near a pre-existing gene may serve to regulate nearby genes in *cis*. We tested this by comparing the bursting kinetics of genes immediately neighboring new *de novo* transcripts to further neighboring genes. Note that in our study, *de novo* transcripts include those unique to *D. melanogaster* and those shared across the *melanogaster-simulans* or the *melanogaster-yakuba* clades. **(B-D)** L1 neighboring genes just upstream of *de novo* transcripts show significantly higher burst sizes than the distribution of all genes during the early and late spermatocyte stages, aligning with the meiosis onsite. These differences remain significant after Benjamini-Hochberg correction (FDR < 0.05). No other significant differences were observed in further neighboring genes or within spermatocytes preceding the *de novo* transcript origination (Figure S8).

Consistent with the cultivator model, the origination of these *de novo* transcripts is associated with a significant increase in transcriptional burst size in their 1^st^ upstream neighboring genes, but not further neighboring genes. These differences are also found concurrently with the meiotic transition, suggesting that these genes may be functionally important in later stages of spermatogenesis. More specifically, significant differences are found either in the early spermatocyte (*D. melanogaster*, p=5.17 x 10^-4^ and *D. yakuba*, p=4.56 x 10^-4^, Mann-Whitney U-test) and late spermatocyte (*D. yakuba*, p=8.25 x 10^-4^) stages. These results remain significant after correction for multiple hypothesis testing using a Benjamini-Hochberg procedure (4 neighbors x 8 clusters x 3 species = 96 hypotheses, FDR < 0.05; Figure 5, Figure S8). Importantly, the cultivator model also predicts that there is no significant difference in transcriptional bursting kinetics prior to the origination of these *de novo* transcripts as these neighboring genes pre-date the appearance of the new *de novo* transcripts in the outgroup species, *D. ananassae*. Consistent with this hypothesis, no significant differences in burst size are detected in any neighboring genes. Furthermore, no significant differences are detected in burst frequency in any neighbors at any stage in any species. Given an increase in burst size but no alteration of burst frequency, these results suggest that a promoter-based effect, rather than an enhancer-based mechanism, drives the *cis*-regulatory effect of new promoter birth. We note that while it is possible that *de novo* transcripts appear next to older genes that already have large burst size kinetics, this does not explain why these neighboring genes show significantly elevated burst size after, but not before, *de novo* transcript origination.

To further validate this finding, we compared the genomic distribution of mean read count per cell (i.e. expression) with the same distributions calculated for L2, L1, R1, and R2 neighboring gene groups (Figure S9, Table S7). While many significant differences in mean read count per cell (cell type-specific expression) were found, most of these significant differences persisted between in-group (*D. melanogaster* and *D. yakuba*) and out-group (*D. ananassae*) comparisons or had opposing directions between species (e.g., significantly increased in *D. melanogaster* and decreased in *D. yakuba*). Only three neighboring group/cell type combinations showed significant differences within in-group species (*D. melanogaster* and *D. yakuba*) while also being directionally consistent: L1 neighbors in early spermatocytes (increase), R1 neighbors in early spermatocytes (decrease), and R1 neighbors in late spermatids (decrease).

In alignment with our previous finding of increased burst size, we find significantly elevated expression of L1 neighboring genes in early spermatocytes in *D. melanogaster* and *D. yakuba* but not *D. ananassae* (Figure S9). Interestingly, we also find significantly decreased expression of R1 neighboring genes in both early spermatocytes and early spermatids when compared to genomic expression. The combination of increased L1 neighboring gene expression as well as decreased R1 neighboring gene expression is particularly striking as it is consistent with supercoiling-mediated coupling (58). The appearance of a new promoter element in the tandem upstream/downstream conformations as shown in Figure 5A would be expected to generate under-winding just upstream of and over-winding just downstream of the new promoter. This would result in both increased transcription in the upstream gene (in this case, the L1 neighbor), and decreased transcription of the downstream gene (in this case, the R1 neighbor).

## Discussion

### Feature selection through ANOVA allows for the identification of conserved cell type-specific genes

Our work demonstrates how the identification of the evolutionarily conserved 198-gene list allowed for cell type classifications from *D. melanogaster* to be extended to a panel of closely related *Drosophila* species. While it was utilized in this study to overcome cell type-specific evolution, the methodology used here should be generalizable to a large range of problems where conditions and transcriptome expression vary in a correlated manner. For example, strong cell type-specific transcriptome-wide responses to external treatment (e.g., heat shock) could obscure cell type assignments. The application of an ANOVA-based methodology could be utilized to identify genes that differ strongly between cell types but remain robust to treatment.

Interestingly, the 198-gene list that we identified did not show a large overlap with more traditionally utilized cell type markers in *D. melanogaster* while still being significantly enriched for previously identified spermatogenesis-related function. One potential interpretation of this observation is that while the markers in the 198-gene list do have known cell type-specific function, it is possible that they do not perform as well as other, more classic markers within *D. melanogaster*. Alternatively, it is also possible that the list of classic marker genes may reflect the historical discovery process regarding *D. melanogaster* spermatogenesis alone rather than underlying biological functions within the larger *melanogaster* species group. Regardless of the reason, our work has shown that the utilization of these marker genes in *D. melanogaster* does not necessarily imply that they are the most optimal markers when examining multiple species’ genomes.

### Biological differences in cell type classification

When we examined the “*ananassae* spermatocyte” cluster, we found biological evidence that there may be meaningful transcriptional differences in these cells compared to other cell types. For example, while these cells are poorly represented in *D. melanogaster* and *D. yakuba*, we found unusual patterns in our differential expression analysis in our full data set. Recall that we found significant differential expression of *His2Av*, an early spermatogenesis marker in *D. melanogaster*, as well as *p-cup*, a late spermatogenesis marker in *D. melanogaster*. Consistent with this observation, we also note the early expression of *Pkd2* in GSC/Early spermatogonia, as well as the occasional presence of cells expressing *Pkd2* well into spermatid differentiation (Figure 2C, E). Notably, neither *His2Av* nor *p-cup* are found in our 198-gene list used for classification. Such transcriptional differences during the spermatocyte stage, combined with the detection of the “*ananassae*-specific spermatocyte” cluster, and unique *D. ananassae* reproductive biology during meiosis (59), suggests that the rapid evolution of the testis may be dominated by changes during this stage, reminiscent of earlier observations that *de novo* genes tend to express most highly in spermatocytes (35).

We also found that *Rpb4* has a surprisingly broad expression pattern in *D. ananassae*. This provokes the interesting question of whether this broad expression pattern is reflective of the ancestral state or evolved in an ancestor of *D. ananassae* subsequent to *D. melanogaster* speciation. It is likely that investigation into the upstream transcriptional differences that drive the alteration of *Rpb4* expression would lead to more general insight into the evolution of transcriptional programming in the testis. Similarly, investigation into the downstream effects of broad *Rpb4* expression would likely be fruitful. *Rpb4*, in conjunction with *fest* and *Syp*, has been shown to regulate *Cyclin B* during meiosis in *D. melanogaster* (49, 50). However, *Rbp4* and *Cyclin B* have much higher cell type specificity in *D. melanogaster* than *D. ananassae*, but in slightly different tissue types. How did these expression patterns evolve? Is the evolutionary appearance of this specificity coupled between the two genes? Alternatively, did this loss of specificity occur only in the *D. ananassae* lineage? These questions require further investigation in the future, particularly with the utilization of additional outgroup species.

We also note that our cross-species categorization detected an absence of spermatids in our scRNA-Seq data. However, our *in-situ* results showed highly specific expression of *soti*, while our brightfield microscopy revealed cells that can clearly be identified as late spermatids based on morphology alone. Interestingly, the gross morphology of these *D. ananassae* spermatids appeared to diverge from *D. melanogaster* as well, as they appeared to be “stringier” and “fuzzier” in addition to downstream morphological differences observed in the accessory gland, where tissue appeared to be larger and whiter. As the total amount of transcripts becomes increasingly depleted as spermatids fully mature, the transcriptional programming controlling these phenotypic differences likely occurs in earlier stages, e.g. “*anannassae* spermatocyte.” Consequently, the downstream biological effects of this altered transcriptional programming will likely be opaque to transcription-based techniques, highlighting the need for proteomic analysis or the application of traditional genetic tools beyond *D. melanogaster*.

### Transcriptional bursting kinetics offers insight into male sex chromosome regulation

Many unanswered questions remain regarding the mechanisms that drive decreased X-chromosome expression during spermatogenesis, leading to a large variety of proposed models. For example, differential polymerase elongation rates have been suggested as a mechanism underlying dosage compensation in males (60). Alternatively, a more canonical pathway involving the lncRNAs *roX1*, *roX2*, and the male-specific lethal complex (MSL) is possible (61, 62). One attractive aspect of the canonical MSL pathway hypothesis is that recent results have shown that the intrinsically disordered C-terminal region of *MSL2* and the *roX* lncRNAs induce X chromosome compartmentalization that functions similarly to phase separation (63). This has led to the suggestion that this X chromosome compartment could be enriched for pol II, resulting in increased initiation of transcription and an increase in transcriptional burst size (27). However, it has also been suggested that elements of the MSL complex, such as *maleless*, do not associate with the X chromosome during spermatogenesis (64).

Our results demonstrate a clear transition from a burst size-dominated mode of regulation to a burst frequency-dominated mode during the meiotic transition, suggesting that this transition may be driven by a global decrease in polymerase availability. However, in alignment with a prior analysis of previous results (31), our analysis of transcriptional bursting kinetics demonstrates that significantly decreased burst size and burst frequency for X-linked genes precedes not only the meiotic transition, but possibly also the establishment of germline stem cells. This decreased burst size is consistent with a study demonstrating decreased pol II binding on X chromosomes derived from both isolated spermatocytes and dissociated testis in comparison to wing imaginal discs (65). While these results are consistent with some manner of incomplete dosage compensation (31, 66), both decreased burst size and burst frequency suggests that a depletion of pol II localized to the X chromosome alone is not a complete explanation for the observed X-linked decrease in expression. More specifically, a local depletion of pol II would decrease transcriptional burst size without altering burst frequency as observed in the developed embryo (27). Instead, the as-yet unknown mechanism for decreased X chromosome expression should affect both the binding rate and the initiation rate of pol II to X chromosome-linked promoter sequences.

Another interesting result from our analyses is species-specific differences in kinetic bursting parameter values. The ‘txburst’ software performs inference over raw read counts, and so it is likely that these differences are related to differences in read depth (Figure S9). Differences in transcript decay rates across species may also be a contributing factor (52). These differences are unlikely to reflect true biological differences in transcriptional burst kinetics, as these differences appear to be unbiased in regard to transcript identity (e.g., housekeeping genes). Importantly, we do not compare kinetic parameters across species, so such differences do not affect our conclusions; further experimental evidence, e.g., single molecule studies, would be needed for proper cross-species comparisons.

## Methods and Materials

Information on methods and materials used are available in the Supplementary Materials. Briefly, we generated scRNA-Seq data from the testes of *D. melanogaster, D. yakuba,* and *D. ananassae*. ANOVA was used to transfer cell type classifications across species and applicability was validated by simulation and HCR RNA FISH. *De novo* transcripts were computationally identified, and bursting kinetics were inferred using maximum likelihood.

## Data and code availability

scRNA-Seq reads were upload to NCBI under the project number PRJNA995212. Scripts for analyses may be found at https://github.com/LiZhaoLab/denovo_bursting_2025.

## Acknowledgments

We thank Helen Duan and Connie Zhao from The Rockefeller University Genomics Resource Center for the help with scRNA-Seq library preparation, RRID:SCR_020986. Confocal microscopy was performed in the Rockefeller University’s Bio-Imaging Resource Center, RRID:SCR_017791. We thank the Zhao lab members for helpful discussion and Sasha Mills for editing help. UL would like to thank Huang Suqi for helpful discussions regarding statistics.

## Author contribution

U.L., N.S., and L.Z. conceived the study, designed the experiments, and formulated the analyses. U.L., C.L., C.B.L, and N.S. generated the data. U.L. conducted all computational analyses. U.L. and L.Z. wrote the paper.

## Funding

This work was supported by National Institutes of Health (NIH) MIRA R35GM133780, the Robertson Foundation, and an Allen Distinguished Investigator Award from Paul G. Allen Family Foundation to L.Z, and National Science Foundation postdoctoral fellowship 2410289 to U.L. The content of this study is solely the responsibility of the authors and does not necessarily represent the official views of the funders.

## Notes

### Competing Interest Statement

The authors have declared no competing interest.

### Summary of Updates

Revised texts and added supplemental figures

