## Supplemental information for "Comparative single cell analysis of transcriptional bursting reveals the role of genome organization on *de novo* transcript origination"

### Supplementary Information

#### 1 Statistical model

##### 1.1 Data matrix

Suppose we have a data matrix, $D_{ij} \in \mathbb{W}$ consisting of raw read counts from a scRNA-Seq experiment, where $i \in \{1, 2, 3, ..., N_{genes}\}$ and $j \in \{1, 2, 3, ..., N_{cells}\}$. The cells are expected to have known cell types $u \in \{1, 2, 3, ..., N_{types}\}$ and species of origin $v \in \{1, 2, 3, ..., N_{species}\}$. We assume that the data matrix $\mathbf{D}$ contains the direct sum of two data matrices $C_{ij}$ and $S_{ij}$ where

$$D_{ij} = \beta_{Ci}\, C_{ij} + \beta_{Si}\, S_{ij} + \epsilon_{ij}\ . \tag{1}$$

Here, $C_{ij}$ is a data matrix containing cell type-related information, $S_{ij}$ contains species-related information, $\beta_{Ci}$ and $\beta_{Si}$ are the relative contribution of cell type and species on a given gene's read counts respectively, and $\epsilon_{ij}$ is an error term. Note that as multiple species are being considered, $i$ generally consists of the set of genes with one-to-one orthologs across all species; $i$ is each species' ortholog.

##### 1.2 Cell-type data matrix

We define the cell-type data matrix $C_{ij}$ as the linear composition of two matrices, $A_{iu}$ and $B_{uj}$, where $\mathbf{A}$ is a $i \times u$ matrix containing information relating gene expression to cell type (i.e. how active or inactive each gene is in a given cell type), and $\mathbf{B}$ is a $u \times j$ matrix containing information relating cell type to sequenced cells (i.e. the classification map of each sequenced cell's correct cell type identity). That is to say,

$$C_{ij} = A_{iu} \times B_{uj}. \tag{2}$$

In general, $A_{ij} \in \mathbb{W}$, where $A_{ij}$ is the read count distribution (expression levels) associated with a given gene in a particular cell type. In the case of binary activation states where genes are either "*on*" (i.e. 1) or "*off*" (i.e. 0), $A_{iu}$ assumes the form

$$\mathbf{A} = \begin{array}{r} \\ gene1 \\ gene2 \\ gene3 \\ gene4 \\ \vdots \\ geneN_{i-1} \\ geneN_i \end{array} \begin{array}{c} \begin{array}{cccccc} type1 & type2 & type3 & type4 & ... & typeN_u \end{array} \\ \left[\begin{array}{cccccc} 1 & 0 & 0 & 0 & \dots & 0 \\ 1 & 0 & 0 & 0 & \dots & 0 \\ 0 & 1 & 0 & 0 & \dots & 0 \\ 0 & 1 & 0 & 0 & \dots & 0 \\ \vdots & \vdots & \vdots & \vdots & \ddots & \vdots \\ 0 & 0 & 0 & 0 & \dots & 1 \\ 0 & 0 & 0 & 0 & \dots & 1 \end{array}\right] \end{array}, \tag{3}$$

when genes 1 and 2 are activated only in cell type 1, genes 3 and 4 are activated only in cell type 2, etc.

We similarly assume that detected cells may have only a single cell type designation, so $B_{uj} \in \{0, 1\}$ assumes the form

$$\mathbf{B} = \begin{array}{c} \\ \textit{type}1 \\ \textit{type}2 \\ \vdots \\ \textit{type}N_u \end{array} \begin{array}{c} \begin{array}{ccccccc} \textit{cell}1 & \textit{cell}2 & \textit{cell}3 & \textit{cell}4 & \ldots & \textit{cell}N_{j-1} & \textit{cell}N_j \end{array} \\ \left[\begin{array}{ccccccc} 1 & 1 & 0 & 0 & \ldots & 0 & 0 \\ 0 & 0 & 1 & 1 & \ldots & 0 & 0 \\ \vdots & \vdots & \vdots & \vdots & \ddots & \vdots & \vdots \\ 0 & 0 & 0 & 0 & \ldots & 1 & 1 \end{array}\right] \end{array}, \tag{4}$$

when cell type 1 is represented by experimentally detected cells 1 and 2, cell type 2 is represented by experimentally detected cells 3 and 4, etc.

#### 1.3 Species data matrix

We also construct a species data matrix, $S_{ij}$ as the linear composition of two matrices, $Q_{iv} \in \mathbb{Z}$ and $R_{vj} \in \{0, 1\}$, where $\mathbf{Q}$ is a $i \times v$ matrix containing information relating evolutionary changes in gene expression to species-of-origin (i.e. how active or inactive each gene is in a species), and $\mathbf{R}$ is a $v \times j$ matrix containing information relating species to sequenced cells (i.e. the classification map of each sequenced cell's species of origin). That is to say,

$$S_{ij} = Q_{iv} \times R_{vj}. \tag{5}$$

Note that unlike $A_{iu} \in \mathbb{W}$, $Q_{iv} \in \mathbb{Z}$ as a given gene's expression may have decreased in a particular species. Again, assuming a binary activation states where genes are either "*on*" (i.e. 1) or "*off*" (i.e. 0), $Q_{iu}$ assumes the form

$$\mathbf{Q} = \begin{array}{c} \\ \textit{gene}1 \\ \textit{gene}2 \\ \textit{gene}3 \\ \textit{gene}4 \\ \vdots \\ \textit{gene}N_{i-1} \\ \textit{gene}N_i \end{array} \begin{array}{c} \begin{array}{cccccc} \textit{species}1 & \textit{species}2 & \textit{species}3 & \textit{species}4 & \ldots & \textit{species}N_v \end{array} \\ \left[\begin{array}{cccccc} 1 & 0 & 0 & 0 & \ldots & 0 \\ 1 & 0 & 0 & 0 & \ldots & 0 \\ 0 & 1 & 0 & 0 & \ldots & 0 \\ 0 & 1 & 0 & 0 & \ldots & 0 \\ \vdots & \vdots & \vdots & \vdots & \ddots & \vdots \\ 0 & 0 & 0 & 0 & \ldots & 1 \\ 0 & 0 & 0 & 0 & \ldots & 1 \end{array}\right] \end{array}, \tag{6}$$

when genes 1 and 2 show elevated expression in species 1 (when compared to other species), genes 3 and 4 show elevated expression in species 2, etc. Note that such species-specific alteration of gene expression may be correlated to the evolution of different cell types. For example, it is possible that the expression of a gene, which is generally expressed in only a single tissue, has altered in one or more species.

As experimentally detected cells may have only a single species-of-origin, $R_{vj}$ assumes the form

$$\mathbf{R} = \begin{array}{c} \\ \textit{species}1 \\ \textit{species}2 \\ \vdots \\ \textit{species}N_v \end{array} \begin{array}{c} \begin{array}{ccccccc} \textit{cell}1 & \textit{cell}2 & \textit{cell}3 & \textit{cell}4 & \ldots & \textit{cell}N_{j-1} & \textit{cell}N_j \end{array} \\ \left[\begin{array}{ccccccc} 1 & 1 & 0 & 0 & \ldots & 0 & 0 \\ 0 & 0 & 1 & 1 & \ldots & 0 & 0 \\ \vdots & \vdots & \vdots & \vdots & \ddots & \vdots & \vdots \\ 0 & 0 & 0 & 0 & \ldots & 1 & 1 \end{array}\right] \end{array}, \tag{7}$$

indicating that in species 1, the experimentally detected cells 1 and 2 have elevated expression; in species 2, the experimentally detected cells 3 and 4 have elevated expression, etc.

#### 1.4 Statement of problem

The issue of cell type assignment may be formally defined as a recovery of the cell type assignment matrix $B_{uj}$ from Eqn. (1). This is performed using some procedure $\mathcal{G}$ such that

$$\mathcal{G} : h, D_{ij} \to \hat{B}_{hj}\,, \tag{8}$$

where $h$ is a pre-determined number of cell types and $\hat{B}_{hj}$ is the estimated cell type assignment matrix.

When the effect of species matrix $S_{ij}$ is sufficiently small and $h$ is appropriately chosen, $\hat{B}_{hj}$ approximates $B_{uj}$. This is often true in many biological scenarios; $S_{ij}$ is always 0 when no species, while the effect of $S_{ij}$ is small in evolutionarily well-conserved tissues. In such cases, $\mathcal{G}$ typically involves dimensional reduction (often a mapping procedure over a few principal components, e.g. UMAP), some manner of unsupervised clustering (e.g. $k$-means clustering where $k = h$), followed by batch correction for species (if applicable). However, when $S_{ij}$ appears to produce a significant effect, particularly in the case where there is a strong cross-correlation between $\mathbf{B}$ and $\mathbf{R}$, $\hat{B}_{hj}$ no longer appears to approximate $B_{uj}$ even after batch correction (c.f. Figure S1). Biologically, this can occur when there is a high degree of species-specific cell type evolution.

We propose a feature-selection criteria $\mathcal{H}_{SQ}$ such that

$$\mathcal{H}_{SQ} : i, D_{i,j} \; \rightarrow \; i', D_{i'j} \tag{9}$$

identifies a gene set $i' \subset i$. We hypothesize that utilization of this gene set $i'$ and associated data sub-matrix $D_{i'j}$ will allow for $\mathcal{G} : h, D_{i'j} \rightarrow \hat{B}_{hj}$ to approximate $\mathbf{B}$ despite a large species effect $\mathbf{R}$ in $\mathbf{D}$. Equivalently, application of the feature-selection criteria $\mathcal{H}_{SQ}$ reduces the effect of $\mathbf{R}$ on $\hat{B}_{hj}$.

### 2 Classification framework

We begin our proposed feature-selection criteria $\mathcal{H}_{SQ}$ with the standard ANOVA F-statistic: $F$. We define $F$ as being calculated for each gene $i$ using a generic cell-label matrix $\mathbf{W}$ corresponding to observed data matrix $D_{ij}$. We will refer to this as

$$F_{i,\mathbf{W}} = F(i, D_{ij}, \mathbf{W}). \tag{10}$$

For each gene $i$, we thus calculate the following two values:

$$F_{i,\,celltype} = F(i, D_{ij}, B_{uj'}) \tag{11}$$

and

$$F_{i,\,species} = F(i, D_{ij}, R_{vj}). \tag{12}$$

where $j' \subset j$ such that $B_{uj'}$ is known. While species-of-origin labeling $R_{vj}$ is fully known for each cell $j$ via barcoding, cell type information is often incomplete. As a result, we generally only calculate $F_{celltype}$ for a subset of cells $j'$ originating from a single reference species with known cell type labeling $B_{uj'}$.

The feature selection criteria $\mathcal{H}_{SQ}$ uses the calculated values $F_{i,\,celltype}$ and $F_{i,\,species}$ to identify $i' \subset i$ such that

$$F_{i',\,celltype} \geq \; cdf^{-1}(p_{celltype}, F_{i,\,celltype}), \tag{13}$$

$$F_{i',\,species} \leq \; cdf^{-1}(p_{species}, F_{i,\,species}). \tag{14}$$

Here, $cdf^{-1}$ represents the inverse cumulative distribution function for $F_{i,\mathbf{W}}$ and $p_{celltype}$ and $p_{species}$ are probability thresholds for the $F$-statistic distributions calculated over cell type and species labels respectively. In practice and in this manuscript, we set $p_{celltype} = p_{species} \equiv p$.

### 3 Simulation

#### 3.1 Simulation model

To validate whether the application of the $\mathcal{H}_{SQ}$ criteria produces $\hat{B}_{hj}$ that can approximate $B_{uj}$ in spite of a large species effect $\mathbf{R}$, we utilized a simulation strategy that allows us to reconstruct $B_{uj}$. Specifically, we utilized a variation of Eqn. (1) to simulate a data set **D'** such that

$$D'_{ij} = (1 - \beta_i)C'_{ij} + \beta_i S'_{ij} + \epsilon_{ij} \,, \tag{15}$$

where $\beta_i \in [0, 1]$ denotes the relative contribution of $C'_{ij}$ and $S'_{ij}$ to $D'_{ij}$. In our simulations, $\beta_i \sim \beta(7,\, 7)$ where $\beta$ represents the classic $\beta$ distribution. In our simulations, we have $i \in \{1, 2, 3, ..., 9999\}$ genes detected in $j \in \{1, 2, 3, ..., 2700\}$ simulated cells. These 2700 simulated cells fall into $u \in \{1, 2, 3, ..., 9\}$ distinct cell types and are sourced from the same organ collected from $v \in \{1, 2, 3\}$ simulated species. We make the simplifying assumption that the phylogeny of the three species constitutes a 3-way tree with equal divergence times.

We simulate the scenario where cells $1 - 900$ are sourced from species 1, cells $901 - 1800$ are sourced from species 2, and cells $1801 - 2700$ are sourced from species 3. Of cells from species 1 (i.e. cells $1 - 900$), cells $1 - 100$ correspond to cell type 1, cells $101 - 200$ correspond to cell type 2, cells $201 - 300$ correspond to cell type 3, ..., and cells $801 - 900$ correspond to cell type 9. Cells from species 2 and 3 also adopt a similar pattern (i.e. cells $901 - 1000$, $1801 - 1900$ correspond to cell type 1, cells $1001 - 1101$, $1901 - 2000$ correspond to cell type 2, etc.).

#### 3.2 Sampling procedure

To simulate read counts in our experimental structure, we define a sampling procedure

$$\mathcal{F} : X_{ij} \to X'_{ij} \,, \tag{16}$$

where $X_{ij} \in \{0, 1\}$ is an $i \times j$ matrix representing the binary activation state of gene $i$ in simulated scRNA-Seq cell $j$ and $X'_{ij} \in \mathbb{W}$ is a $i \times j$ matrix representing the simulated read count of gene $i$ in simulated scRNA-Seq cell $j$.

We then implement our sampling procedure $\mathcal{F}$ such that

$$\mathcal{F} : \begin{cases} X'_{ij} \sim y_{off} & \text{iff} \quad X_{ij} = 0 \\ X'_{ij} \sim y_{on} & \text{iff} \quad X_{ij} = 1 \end{cases} \,, \tag{17}$$

where $y_{off}$ and $y_{on}$ are empirical distributions derived from our real *D. melanogaster* testis scRNA-Seq data. $y_{off}$ is the read count distribution of Fbgn0038531 (as its mean expression value is in the 80th percentile of all genes in our data set), while $y_{on}$ is the read count distribution of Fbgn0037345 (as its mean expression value is 95th percentile of all genes in our data set). The 80th and 95th percentiles were chosen because genes with lower mean expression values tended to have read counts of 0 in the vast majority of cells. Thus, to generate our simulated data set **D'**, we now construct our cell type matrix **C** and species matrix **S** to generate **C'** and **S'** respectively using our sampling procedure $\mathcal{F}$.

#### 3.3 Cell type identity

To construct $C_{ij}$, we now specify the structure of its compositional matrices $A_{iu}$ and $B_{uj}$. Recall that $\mathbf{A}$ is the matrix relating gene expression to cell type. Without loss of generality, we make the simplifying assumption that a gene is "*on*" in only a single tissue type and "*off*" in all other tissue types; mapping onto $\mathbb{W}$ for more realistic read count distributions will occur via $\mathcal{F}$. In this case, we define

$$
\mathbf{A} = \begin{array}{r} \\ gene1 \\ gene2 \\ \vdots \\ gene1111 \\ gene1112 \\ gene1113 \\ \vdots \\ gene2222 \\ gene2223 \\ gene2224 \\ \vdots \\ gene9998 \\ gene9999 \end{array}
\begin{array}{c}
\begin{array}{ccccc} type1 & type2 & type3 & \dots & type9 \end{array} \\
\begin{bmatrix}
1 & 0 & 0 & \dots & 0 \\
1 & 0 & 0 & \dots & 0 \\
\vdots & \vdots & \vdots & \ddots & \vdots \\
1 & 0 & 0 & \dots & 0 \\
0 & 1 & 0 & \dots & 0 \\
0 & 1 & 0 & \dots & 0 \\
\vdots & \vdots & \vdots & \ddots & \vdots \\
0 & 1 & 0 & \dots & 0 \\
0 & 0 & 1 & \dots & 0 \\
0 & 0 & 1 & \dots & 0 \\
\vdots & \vdots & \vdots & \ddots & \vdots \\
0 & 0 & 0 & \dots & 1 \\
0 & 0 & 0 & \dots & 1
\end{bmatrix}
\end{array}. \tag{18}
$$

Similarly, recall that **B** is the matrix relating cell type to simulated sequenced cells, where a simulated cell may only posses a single cell type. In this case, we define

$$
\mathbf{B} =
$$

$$
\begin{array}{r} \\ type1 \\ type2 \\ \vdots \\ type9 \end{array}
\begin{array}{c}
\begin{array}{ccccccccccc} cell1 & \dots & cell100 & cell101 & \dots & cell900 & cell901 & \dots & cell1800 & cell1801 & \dots & cell2700 \end{array} \\
\begin{bmatrix}
1 & \dots & 1 & 0 & \dots & 0 & 1 & \dots & 0 & 1 & \dots & 0 \\
0 & \dots & 0 & 1 & \dots & 0 & 0 & \dots & 0 & 0 & \dots & 0 \\
\vdots & \ddots & \vdots & \vdots & \ddots & \vdots & \vdots & \ddots & \vdots & \vdots & \ddots & \vdots \\
0 & \dots & 0 & 0 & \dots & 1 & 0 & \dots & 1 & 0 & \dots & 1
\end{bmatrix}
\end{array}. \tag{19}
$$

The resulting matrix **C** is thus

$$
\mathbf{C} =
$$

$$
\begin{array}{r} \\ gene1 \\ gene2 \\ \vdots \\ gene1111 \\ gene1112 \\ \vdots \\ gene2222 \\ gene2223 \\ \vdots \\ gene8888 \\ gene8889 \\ \vdots \\ gene9999 \end{array}
\begin{array}{c}
\begin{array}{cccccccccccc} cell1 & \dots & cell100 & cell101 & \dots & cell900 & cell901 & \dots & cell1800 & cell1801 & \dots & cell2700 \end{array} \\
\begin{bmatrix}
1 & \dots & 1 & 0 & \dots & 0 & 1 & \dots & 0 & 1 & \dots & 0 \\
1 & \dots & 1 & 0 & \dots & 0 & 1 & \dots & 0 & 1 & \dots & 0 \\
\vdots & \ddots & \vdots & \vdots & \ddots & \vdots & \vdots & \ddots & \vdots & \vdots & \ddots & \vdots \\
1 & \dots & 1 & 0 & \dots & 0 & 1 & \dots & 0 & 1 & \dots & 0 \\
0 & \dots & 0 & 1 & \dots & 0 & 0 & \dots & 0 & 0 & \dots & 0 \\
\vdots & \ddots & \vdots & \vdots & \ddots & \vdots & \vdots & \ddots & \vdots & \vdots & \ddots & \vdots \\
0 & \dots & 0 & 1 & \dots & 0 & 0 & \dots & 0 & 0 & \dots & 0 \\
0 & \dots & 0 & 0 & \dots & 0 & 0 & \dots & 0 & 0 & \dots & 0 \\
\vdots & \ddots & \vdots & \vdots & \ddots & \vdots & \vdots & \ddots & \vdots & \vdots & \ddots & \vdots \\
0 & \dots & 0 & 0 & \dots & 0 & 0 & \dots & 0 & 0 & \dots & 0 \\
0 & \dots & 0 & 0 & \dots & 1 & 0 & \dots & 1 & 0 & \dots & 1 \\
\vdots & \ddots & \vdots & \vdots & \ddots & \vdots & \vdots & \ddots & \vdots & \vdots & \ddots & \vdots \\
0 & \dots & 0 & 0 & \dots & 1 & 0 & \dots & 1 & 0 & \dots & 1
\end{bmatrix}
\end{array}. \tag{20}
$$

#### 3.4 Species-specific cell type evolution

We also construct $S_{ij}$ by specifying the structure of its compositional matrices $Q_{iv}$ and $R_{vj}$. Recall that **Q** is the matrix relating evolutionary gene expression changes to species. We make the simplifying assumption that gene expression has only increased in a binary fashion; as before, mapping onto $\mathbb{W}$ for more realistic read count distributions will occur via $\mathcal{F}$. In this case, we define

$$\mathbf{Q} = \begin{array}{c} \\ gene1 \\ gene2 \\ \vdots \\ gene3333 \\ gene3334 \\ gene3335 \\ \vdots \\ gene6666 \\ gene6667 \\ gene6668 \\ \vdots \\ gene9998 \\ gene9999 \end{array} \begin{array}{c} \begin{array}{ccc} species1 & species2 & species3 \end{array} \\ \left[\begin{array}{ccc} 1 & 0 & 0 \\ 1 & 0 & 0 \\ \vdots & \vdots & \vdots \\ 1 & 0 & 0 \\ 0 & 1 & 0 \\ 0 & 1 & 0 \\ \vdots & \vdots & \vdots \\ 0 & 1 & 0 \\ 0 & 0 & 1 \\ 0 & 0 & 1 \\ \vdots & \vdots & \vdots \\ 0 & 0 & 1 \\ 0 & 0 & 1 \end{array}\right] \end{array} . \tag{21}$$

This definition of $\mathbf{Q}$ states that genes $1 - 3333$ have increased their expression levels exclusively in species 1, genes $3334 - 6666$ have increased their expression levels in exclusively in species 2, and genes $6667 - 9999$ have increased their expression exclusively in species 3. Note that these genes are expressed in a tissue-specific manner. That is to say that in all 3 species, genes $1 - 1111$ are expressed solely in cell type 1, genes $1112 - 2222$ are expressed solely in cell type 2, and genes $2223 - 3333$ are expressed solely in cell type 3. As a result, the definition of $\mathbf{Q}$ simulates a scenario in which species 1 has increased expression of these cell type-related genes in a species-specific manner, i.e. the genes denoting cell type $1 - 3$ have higher expression in species 1 than in species 2 or species 3. Similarly, genes expressed in cell types $4 - 6$ have higher expression in species 2 than in species 1 or species 3; genes expressed in cell types $7 - 9$ have higher expression in species 3 than in species 1 or species 2.

Specifically, the matrix $\mathbf{Y} \equiv \mathbf{A^T} \times \mathbf{Q}$, representing species-specific cell type evolution, becomes

$$\mathbf{Y} =$$

$$\begin{array}{c} \\ type1 \\ type2 \\ type3 \\ type4 \\ type5 \\ type6 \\ type7 \\ type8 \\ type9 \end{array} \begin{array}{c} \begin{array}{ccc} species1 & species2 & species3 \end{array} \\ \left[\begin{array}{ccc} 1111 & 0 & 0 \\ 1111 & 0 & 0 \\ 1111 & 0 & 0 \\ 0 & 1111 & 0 \\ 0 & 1111 & 0 \\ 0 & 1111 & 0 \\ 0 & 0 & 1111 \\ 0 & 0 & 1111 \\ 0 & 0 & 1111 \end{array}\right] \end{array} , \tag{22}$$

indicating that cell types 1, 2, and 3 have increased expression in species 1, cell types 4, 5, and 6 have increased expession in species 2, and cell types 7, 8, and 9 have increased expression in species 3.

Recall that matrix $\mathbf{R}$ is the matrix relating species of origin to simulated cells. We define

$$\mathbf{R} =$$

$$\begin{array}{c} \\ species1 \\ species2 \\ species3 \end{array} \begin{array}{c} \begin{array}{ccccccccc} cell1 & cell2 & \dots & cell900 & cell901 & \dots & cell1800 & cell1801 & \dots & cell2700 \end{array} \\ \left[\begin{array}{cccccccccc} 1 & 1 & \dots & 1 & 0 & \dots & 0 & 0 & \dots & 0 \\ 0 & 0 & \dots & 0 & 1 & \dots & 1 & 0 & \dots & 0 \\ 0 & 0 & \dots & 0 & 0 & \dots & 0 & 1 & \dots & 1 \end{array}\right] \end{array} . \tag{23}$$

The resulting matrix **S** is thus

$$\mathbf{S} = \begin{array}{c} \\ gene1 \\ gene2 \\ \vdots \\ gene3333 \\ gene3334 \\ \vdots \\ gene6666 \\ gene6667 \\ \vdots \\ gene9999 \end{array} \begin{array}{c} \begin{array}{cccccccccc} cell1 & cell2 & ... & cell900 & cell901 & ... & cell1800 & cell1801 & ... & cell2700 \end{array} \\ \begin{bmatrix} 1 & 1 & \dots & 1 & 0 & \dots & 0 & 0 & \dots & 0 \\ 1 & 1 & \dots & 1 & 0 & \dots & 0 & 0 & \dots & 0 \\ \vdots & \vdots & \ddots & \vdots & \vdots & \ddots & \vdots & \vdots & \ddots & \vdots \\ 1 & 1 & \dots & 1 & 0 & \dots & 0 & 0 & \dots & 0 \\ 0 & 0 & \dots & 0 & 1 & \dots & 1 & 0 & \dots & 0 \\ \vdots & \vdots & \ddots & \vdots & \vdots & \ddots & \vdots & \vdots & \ddots & \vdots \\ 0 & 0 & \dots & 0 & 1 & \dots & 1 & 0 & \dots & 0 \\ 0 & 0 & \dots & 0 & 0 & \dots & 0 & 1 & \dots & 1 \\ \vdots & \vdots & \ddots & \vdots & \vdots & \ddots & \vdots & \vdots & \ddots & \vdots \\ 0 & 0 & \dots & 0 & 0 & \dots & 0 & 1 & \dots & 1 \end{bmatrix} \end{array}. \quad (24)$$

##### 3.5 Simulated data set and assessment of accuracy

Now that we have **C** and **S**, we apply our sampling procedure $\mathcal{F}$ to obtain **C'** and **S'**. We also generate $\beta_i$ by sampling $\beta(7,7)$ for each gene index $i$. Finally, we generate an error term $\epsilon_i$ such that $\epsilon_i \sim y_{off} \cup y_{on}$. We then combine these terms using Eqn. 15 to obtain **D'**. A heatmap of the resulting simulated data matrix **D'** is found in Figure S2A.

We then apply the feature selection criteria $\mathcal{H}_{SQ}$ to identify gene set $i'$. In our application of $\mathcal{H}_{SQ}$, we began with $p = 0.5$ and gradually increased it until the cardinality of gene set $|i'| = 200$. This was an arbitrary choice to reflect the observation that $|i'| = 198$ when we chose $p = 0.5$ in our scRNA-Seq data set. As such, we aimed to assess the accuracy of $\hat{B}_{hj}$ when $|i'| \approx 198$.

When our feature selection criteria $\mathcal{H}_{SQ}$ is applied, it is helpful to visualize $F_{i,\,celltype}$ and $F_{i,\,species}$ on a per-cell basis. This is shown in Figure S2B. The resulting high degree of correlation between $F_{i,\,celltype}$ and $F_{i,\,species}$ indicates that **R** has a large effect on $\hat{B}_{hj}$.

To assess the accuracy of $\hat{B}_{hj}$ when compared to $B_{uj}$, we use two different methodologies:

First, as the $\mathcal{H}_{SQ}$ feature selection criteria is non-parametric, we may compare the distribution of $\beta_{i'}$ to $\beta_i$. If $\hat{B}_{hj}$ is a good estimator of $B_{uj}$, we expect $\beta_{i'}$ to be distributed close to 0, indicating that $D'_{i'}$ reflects a large cell type effect $C'_{i'}$ and a low species-specific effect $S'_{i'}$. Alternatively, if $\hat{B}_{hj}$ is a poor estimator of $B_{uj}$, we expect $\beta_{i'}$ to not be distributed close to 0. Interestingly, if $\hat{B}_{hj}$ is a good estimator of $R_{vj}$, and not $B_{uj}$, we expect $\beta_{i'}$ to distributed close to 1. The resulting distributions of $\beta_{i'}$ and $\beta_i$ are found in Figure S2C.

We also perform UMAP on both the full simulated data set $D'_{ij}$ as well as the subsetted data set $D'_{i'j}$ to visually estimate $\hat{h}$. If $\hat{B}_{hj}$ is a poor estimator of $B_{uj}$, we expect $\hat{h} = N_u \times N_v = 27$. That is to say that we expect to see 27 individual clusters. In such a scenario, it is impossible to provide proper cell type assignment across different species, as the cluster representing a given cell type in one species is separate from the cluster representing the same cell type in a different species (e.g. species 1's cell type 1 is a distinct cluster from species 2's cell type 1). This can be seen in the UMAP visualization of the full data set $D'_{ij}$ provided in Figure S2D. Alternatively, if $\hat{B}_{hj}$ is a good estimator of $B_{uj}$, we expect $\hat{h} = N_u = 9$. This can be seen in the UMAP visualization of the subsetted data set $D'_{i'j}$ provided in Figures S2E-G. Note that if $\hat{B}_{hj}$ is a good estimator of $R_{vj}$ and not $B_{uj}$, we expect $\hat{h} = N_v = 3$.

### 4 Performance of $\mathcal{H}_{SQ}$ criteria in asymmetric phylogenies

While the symmetric phylogenetic structure implemented in Section 3 allow for a simpler simulation strategy, such structures are not applicable in comparative studies that utilize a known out-group species. To determine whether application of the $\mathcal{H}_{SQ}$ criteria is able to recover $B_{uj}$ in asymmetric phylogenies, we simulate two different scenarios that result in species 1 and species 2 sharing more similarities than with species 3.

We note that there is no explicit parameterization of divergence time or phylogenetic structure in section 3. There is also no clear methodology for assessing divergence time using single cell transcriptomes. In the absence of these two facets, we may intuit transcriptomic similarity may be used as a proxy for divergence. Specifically, we mean that species with similar transcriptomes are more related than two species with differing transcriptomes. What does it mean, however, to have a similar transcriptome when considering multiple cell types? If one applies the simplifying assumption that overall transcriptomic "similarity" is a linear function each cell types' "similarity", that is to say that species with a higher number of similar/common cell types are more related than species with fewer similar or common cell types, we may utilize the number of similar cell types as a proxy for evolutionary divergence. That is say that diverged species have fewer cell types in common with less diverged species.

Given this assumption, we then simulate scenarios with asymmetric phylogenies by varying the number of common cell types shared between a set of three species. Specifically, we simulate scenarios where either two cell types (scenario A) or all nine cell types (scenario B) expression patterns are shared between species 1 and 2 but not species 3. While it is likely that naturally occurring species diverge in a much more complex manner than what is reflected in these simulations, this strategy illustrates a simple, effective, and easily interpretable set of simulations in which we can test the reliability our methodology with asymmetric phylogenies.

We may understand this simulation strategy more easily through visualization of both the simulated raw count matrices as well as UMAP projections of these data set. The simulated raw count matrix of scenario A (Figure S3A) shows how the expression pattern of cell types 3 and 6 are identical within species 1 and species, but is altered in species 3. This is readily demonstrated in the UMAP visualization of the full data set, where two cell types from species 1 and species 2 overlap (Figure S3B). Similarly, the simulated raw count matrix of scenario B (Figure S3C) shows how the expression patterns of cell types 1 through 9 are identical between species 1 and species 2, but is altered in species 3. The UMAP visualization of the full data set reveals that the cell types from species 3 remain distinct from the cell types from species 1 and species 2 (Figure S3D). We then took the top 200 genes as described in Section 3.5. In both scenarios A and B, cell type assignments can clearly be transferred accurately across all species (Figure S3E-F, G-H).

The simulations presented here represent only one of many methods by which asymmetric phylogenies may be represented. However, given the assumption that these structures reflect asymmetric phylogenies, we stress these results span the entire range of possible asymmetries in 3-way trees. The simulations from the previous sections reflect a three-way phylogeny where all branch lengths are equal, i.e. zero asymmetry. Scenario A represents some intermediate situation between maximal and minimal asymmetries, while Scenario B represents the maximally asymmetric 3-way phylogeny. If we assume that the effectiveness of our methodology either monotonically increases or decreases as asymmetry is varied, our results therefore demonstrate that our methodology is effective across the range of all possible asymmetries so far as asymmetry is defined in this set of simulations.

### 5 scRNA-Seq library preparation and sequencing

Testis from D. melanogaster, D. yakuba, and D. ananassae were dissected in ice-cold PBS before being transferred to ice. Dissociation was performed via desheathing using a lysis buffer consisting of 2% collagenase (100mg/ml stock) in 1X TrypLE buffer. Samples were centrifuged, then mildly vortexed, and then incubated at room temperature for 10 minutes. This procedure was repeated three times (30 minutes total), then passed through 35 µm cell strainers. The resulting samples were centrifuged for 7 minutes at 1200 rpm at 4°C. The resulting pellet was washed with 200 µL of cold HBSS and centrifuged again for 7 minutes at 1200 rpm at 4°C. The resulting pellet was suspended in 35 µL of cold HBSS. 5 µL of the resulting single cell suspension was stained with 5 µL of trypan blue for cell counting and the remaining 30 µL of suspension was used for library preparation and sequenced using the 10X platform by the Rockefeller University Genomics Resource Center.

#### 6 scRNA-Seq pipeline and analysis

Raw data was processed using cellranger (v7.1.0) and aligned to FlyBase genomes (dmel-r6.44, dyak-r1.05, and dana-r1.06). Resulting count matrices were loaded into Monocle3, where only "melanogasterized" genes with one-to-one orthologs reported by FlyBase were retained. Ribosomal genes, tRNA genes, and mitochondrial genes were also excluded from further analysis. These data sets were preprocessed and aligned using Monocle3. Differential expression analyses were performed in Seurat (v 4.1.3) using cell type assignments performed in Monocle3. Further information may be found in the Supplementary Information.

Count matrices were combined and cell type labels from a previous study [1] were imported, and species labels were applied. ANOVA F-statistics were calculated for each gene as described in Statistical procedure and the resulting 198-gene list was retained. Raw counts were then extracted for the 198-gene list in Monocle3 and then subsequently combined, preprocessed, and aligned using default settings. Note that these default settings apply a batch-correction step by species within our 198-gene list. Dimensional reduction was performed and leiden clustering was performed using default settings with 3x10-4 resolution. This resulted in 9 clusters, where two clusters overlapping with early spermatocytes were combined and two other clusters overlapping with early spermatids were also combined. Differential expression analyses were performed using Seurat's 'FindMarker' function. Unless otherwise noted, differential analyses between assigned cell types was performed without respect to species identity. For example, we have taken all cells labeled as GSC/Early spermatocyte from all species, and compared them to all other cells from all species.

#### 7 In situ hybridization chain reaction (HCR RNA FISH)

Dissection of testis tissue followed [2] and hybridization chain reaction protocols from [3] were adapted for Drosophila testis. Oligo pools/probes were generated using HCRProbeMakerCL v2.0 (https://github.com/rwnull/HCRProbeMakerCL). Specific probe sequences paired with amplifier information are provided in Table S8.

Mixed-aged Drosophila males (w1118 for D. melanogaster, and D. ananassae received from Vanessa Ruta lab) were dissected in untreated welled slides containing droplets of ice-cold testis isolation buffer [2]. Tissues were dissected for no longer than 10 minutes to preserve tissue freshness. Note, all subsequent steps were performed in welled slides while being observed in a microscope to prevent tissue loss. Fresh tissue was then transferred to room temperature and remaining isolation buffer was replaced with 3.7% formaldehyde suspended in 1X PBS buffer at room temperature. After a 30-minute fixation, tissue was washed three times in 1X PBS buffer. Fixed tissue was then processed using [3]. As subsequent steps were performed under observation in a welled slide, reaction volumes were adjusted such that all tissues were submerged in relevant solutions and buffers (generally 50-70 uL). Probes were synthesized from IDT while necessary buffers and amplifiers were ordered from Molecular Instruments. B1, B3, B4, and B5 amplifiers were combined with Alexa-647, Alexa-594, Alexa-546, and Alex-488 respectively. Due to significant cross-talk between Alexa-594 and Alexa-546 channels in microscopy, samples were prepared using combinations of either (B1, B3, and B5) or (B1, B4, and B5). Processed tissue was then transferred to poly-L-lysine coated microscopy slides using 70% glycerol in 1X PBS as mounting media, covered using hydrophobic cover slips, and sealed with nail polish. The resulting slides were stored at 4C until imaging was completed.

Confocal imaging was performed using a Zeiss LSM 980 microscope with Airyscan 2 running Zeiss Zen Blue acquisition software. All subsequent image processing was also performed in Zeiss Zen Blue. Z-stacks were generated across the entire tissue and automated stitching was performed. Subsequent z-stacks were flattened using orthogonal projection using maximal intensity and resulting intensity profiles were adjusted using recommended automatic settings on a per-channel basis. Finally, the overall intensity profiles of the merged figure were manually adjusted to decrease saturation.

### 8 Bulk RNA-Seq pipeline and identification of de novo transcripts

We used both HALPER [4], a HAL-based [5] methodology that relies on ProgressiveCactus [6] multi-genome alignments, and LiftOff [7], a minimap2-based [8] methodology which does not require multi-genome alignments and thus does not rely on synteny. We then obtained a panel of publicly available bulk testis RNA-Seq data sets from these species (SRR1024003, SRR1024002, SRR1617567, SRR486109, SRR5839446, SRR16213299, SRR16213300, SRR6667440, SRR6667439, SRR931549) and calculated TPM values for both annotations generated through HALPER and LiftOff. This approach thus allows us to identify the homologous regions in outgroup species for transcripts, allowing us to positively identify non-detection events in outgroup species.

### 9 Inference of transcriptional bursting properties

Raw cell counts were exported into .csv files for each cell type. Each cell count file was then input into 'txburstML.py' from [9]. Resulting .pkl files were then converted and then imported into R. Genes failing quality-control from 'txburstML.py' were discarded, and orthology calls were made using FlyBase orthology information ("dmel_orthologs_in_drosophila_species_fb_2022_01"). Inferred burst size and burst frequency values were further analyzed in R.

### 10 '*ananassae*' spermatocyte cluster

An important aspect of our scRNA-Seq analyses is the detection of a new "*ananassae* spermatocyte" cluster and correlated loss of a spermatid cluster in our *D. ananassae* data set. This inevitably leads to questions regarding whether these represent new cell types or are instead the result of technical artefacts like differing cell recovery during scRNA-Seq library preparation. This is a critical question that needs to be directly addressed. We also expect that the continued development of scRNA-Seq technology will likely result in a correlated increase in the utilization of scRNA-Seq analyses for comparative studies. As such, providing a comprehensive and thorough answer to these questions is of the utmost importance. In the following section, we will first succinctly provide our answer to this question, followed by a longer discussion of our reasoning for that answer.

We believe that our detection of these new clusters is probably not the result of the evolutionary appearance of a new, distinct cell type. We also do not believe that it is the result of technical differences in cell type recovery within our final scRNA-Seq libraries. We further assert that the detection of new UMAP clusters in our data is not artefactual in nature, but is instead reflective of changes in the underlying biology of evolutionarily conserved cell types. In alignment with the most prevalent interpretive framework for scRNA-Seq data, we utilize our distinct UMAP cluster as an "*ananassae* spermatocyte" cell type in our figures as we believe that a departure from this framework would currently be unnecessarily distracting. The rationale for this choice is discussed later in this section. However, we believe that future studies will likely need to more firmly address certain discrepancies resulting from this framework.

We must first reconsider the commonly-held framework for cell type assignments in scRNA-Seq data sets. Typically, distinct clusterings in UMAP projections are assumed to be representative of distinct cell types. For example, two distinct clusters in a UMAP projection is generally assumed to represent two distinct cell types. We will refer to this framework as a "cell type-focused" framework. Implicit within the cell type-focused framework is the assumption that differing expression profiles result from intrinsic functional differences that are encapsulated and represented by a given cell's cell type assignment. Differences in expression patterns are similarly assumed to be the result of differences in cell type identities.

We wish to challenge this framework instead by describing what we refer to as an "expression pattern-focused" framework. We begin with the following observation: clusterings in UMAP projections strictly represent expression profile similarities within distinct sets of detected cells. While differences in cell type identity represent one reason why expression profiles may vary, there remain many other reasons why expression profiles may vary. For example, environmental effects, such as temperature changes or drug treatment, or simple evolutionary divergence represent two of many reasons why expression profiles may vary. This is to say that any given UMAP cluster can not only reflect cell type differences, but may also represent a number of other factors. We thus assert that expression pattern-focused frameworks may be more applicable than a cell type-focused framework in certain classes of comparative studies. An important distinction resulting from this shift in framework is that a given UMAP cluster does not necessarily reflect a difference in cell type assignment. Instead, an expression-pattern focused framework would imply that a given UMAP cluster is representative of a given stereotypical expression pattern, not necessarily a distinct cell type. These stereotypical expression patterns may, indeed, correlate with different cell type identities, but may also correlate with a large number of other factors as described above.

Re-examination of our scRNA-Seq data under an expression pattern-focused framework thus alters our interpretation of our UMAP clusterings. The UMAP cluster that we refer to as the "*ananassae* spermatocyte" is, more precisely, a set of cells that demonstrate a particular stereotypical expression pattern. This particular expression pattern, at least as assessed over our subset of 198 genes, is not shared by a significant number of cells in the *D. melanogaster* and *D. yakuba* data sets. Recall as well that these 198 genes were selected to have a small species-specific signal. This means that each individual gene in the 198-gene set is similarly expressed across species, at least when considered in bulk. However, when these genes are considered in *combination*, a new stereotypical expression pattern emerges that is shared by a large number of *D. ananssae* cells, but is shared by a very low number of cells in *D. melanogaster* and *D. yakuba*. Assuming that these changes are driven by genomic changes, not environmental or epigenetic factors, we may further conclude that the *ananassae* spermatocyte expression pattern is the result of a set of regulatory differences upstream of these 198 genes. This is not to say that there may not also be expression differences downstream of these 198 genes, but insofar as the stereotypical *ananassae* spermatocyte expression pattern *in the 198-gene set* is concerned, expression differences may be attributed to upstream regulatory changes either in *cis-* or in *trans-*.

While it is unclear to what degree the appearance of the *ananassae* spermatocyte expression pattern reflects a true alteration of cell type identity, we believe that it is likely that not reflective of a difference in cell type assignment. Cell type assignments in the testis have been classically defined through a combination of mitotic stage and gross morphological changes that far precede the development of scRNA-Seq data. The initial challenge of aligning previously unalignable whole-transcriptome scRNA-Seq data relies on this "ground truth" of cell type identity. If the fundamental assumption of the existence of cross-species cell types is discarded, the inability to align cross-species data is not a barrier to subsequent analysis. For example, if scRNA-Seq data sourced from *D. melanogaster* testis tissue and *D. ananassae* head tissue were combined, the ability to transfer cell type assignments from *D. melanogaster* to *D. ananassae* would be highly troubling. Similarly, the inability to transfer cell type assignments from *D. melanogaster* testis tissue to *D. ananassae* testis tissue prompted us to develop a new methodology to allow cross-species cell type assignments.

The existence of this "ground truth" allows us to assume that cell identity supercedes global expression pattern differences. We may thus conclude that global transcriptomic differences between *D. melanogaster* and *D. ananassae* expression patterns is not reflective of underlying differences in cell type. For example, the function of a germline stem cell remains conserved regardless of species-of-origin, even if scRNA-Seq derived from these cells defies alignment. Similarly, we may assume that the "*ananassae* spermatocyte" clustering does not truly reflect a previously unobserved cell type. We already accept the assumption that cell type identity is immutable when considering the full transcriptome. It should also be fair to assume that cell type identity remains immutable when considering any subset of the transcriptome as well (i.e. our 198-gene set). Given this assumption, then, we may then conclude that the shifts in cell type representation are instead shifts in the type and degree of stereotypical expression patterns represented in our collection of testis-derived cells. The detection of an additional "*ananassae* spermatocyte" cluster and the loss of spermatid cells in the *D. ananassae* data set are not representative of true gains and losses of different cell types. They are instead simply reflective of expression pattern differences in the 198-gene set. We note, however, that the existence of this "ground truth" of cell identity is an assumption. In the case that a "truly new" cell type with an evolutionarily novel function appears, this a very poor assumption to make, subject to the ability to define a "truly new" cell type outside of the context of scRNA-Seq clusterings.

We also highlight that the previous conclusion is parsimonious even outside of the context of a "ground truth" assumption, as true differences cell type representation requires additional complications. For example, we need to invoke additional cell division or death events if there are true biological differences in spermatocyte and spermatid production. Similarly, we would need to invoke a separate biological mechanism for explaining why collagenase treatment of *D. ananassae* testis tissue would release a different number of spermatocytes or spermatids in comparison to *D. melanogaster*. While it is certainly possible that these mechanisms may exist, such events have yet to be described in the literature.

A new expression pattern-focused framework, in contrast to a cell type-focused framework, is also powerful in that it allows us to further consider the functional changes that drive the expression divergence. We have demonstrated that the functions associated with our 198 genes are enriched for sperm mobility, protein maturation/degradation, insulin signaling, and more. This reflects the observation that differences between earlier and later germline differentiation stages (e.g. GSC/Early spermatogonia vs Late spermatid) are generally related to such functions. Similarly, we can also conclude that some of the most evolutionarily important differences between *D. melanogaster* and *D. ananassae* are also likely related to changes to these functions.

To further understand how these functional changes are manifested, we again consider the set of stereotypical expression patterns between D. melanogaster and D. ananassae. Assuming there are significant differences between the developmental trajectories of D. melanogaster and D. ananassae, we now consider the following question: at what stage do differences in D. melanogaster and D. ananassae begin to manifest?

We address this question by first examining the stereotypical "*ananassae* spermatocyte" expression pattern that appears in exclusively in our *D. ananassae* data set (and not in the *D. melanogaster* data set.) Visual inspection of the cross-species UMAP clearly indicates that there is a set of cells that most represent this expression pattern. To further explore this cell type, we identified a gene that is highly and most specifically expressed in cells that follow this "*ananassae* spermatocyte" pattern: *Pkd2* (Figure 2). Our RNA FISH results for *Pkd2* show that the cells most closely following this "*ananassae* spermatocyte" pattern are clearly both early- and late-stage spermatocyte cells.

The importance of the expression pattern-focused framework is also highlighted when considering *Pkd2* expression in *D. melanogaster*. It is clear that there are few cells in *D. melanogaster* (as defined by *Pkd2* expression relative to morphological/mitotic staging) that do not also appear within the "*ananassae* spermatocyte" cluster. Why, then, is *Pkd2* expression robustly detected in *D. melanogaster*, particularly when there are very few "*ananassae* spermatocyte cells" in *D. melanogaster* scRNA-Seq data? A cell type-focused framework implies that *Pkd2* is a marker gene for "*ananassae* spermatocytes," a cell type that is "unique" to *D. ananassae*. The RNA FISH data would also paradoxically imply that a large number of these *D. ananassae*-specific cells is also detected in *D. melanogaster*. The observation that these cells in *D. melanogaster* are also clearly spermatocytes based on morphological and mitotic staging, leading to further confusion.

This apparent contradiction is resolved under the lens of an expression pattern-focused framework. First, this framework implies that there are simply fewer cells in *D. melanogaster* whose expression patterns (within the 198-gene set) are similar to the stereotypical "*ananassae* spermatocyte" expression pattern. Secondly, this does not exclude the possibility that spermatocytes from both *D. melanogaster* and *D. ananassae* express *Pkd2* - this is indeed what is observed.

We now return to the original discussion of whether these cells represent a distinct cell type. We note that an extended discussion of what is meant by a "cell type" and "stereotypical expression pattern" would likely be distracting and confusing to readers that are familiar with scRNA-Seq data and are not particularly concern with such distinctions. Ultimately, the semantics of referring a set of cells possessing a particular expression pattern vs. a cell type may not be particularly relevant outside of the results mentioned in this section. As such, we have been careful to refer to the "*ananassae* spermatocyte" clustering as merely that: a distinct cluster. However, we do additionally treat it as a distinct "cell type" in our figures and analyses despite differences in representation between *D. melanogaster* and *D. ananassae*. We do, however, provide an extended discussion of the matter as we believe that this is an important issue that will become increasing relevant with an increasing number of comparative scRNA-Seq experiments addressing factors like treatment effects or evolutionary divergence. Regardless, this discussion highlights the importance of combining multiple sources of evidence, e.g. gross morphology or RNA FISH data, for cell type classification when considering scRNA-Seq data.

### 11 Supplementary Tables

#### Table S1: Summary Sequencing QC Statistics

| | *D. melanogaster* | *D. yakuba* | *D. ananassae* |
|---|---|---|---|
| reference genome | dmel_r6.44 | dyak_r1.05 | dana_r1.06 |
| Estimated Number of Cells | 5,000 | 5,000 | 5,000 |
| Mean Reads per Cell | 85,313 | 94,561 | 78,520 |
| Median Genes per Cell | 4,422 | 2,458 | 2,206 |
| Number of Reads | 426,563,073 | 472,805,835 | 392,602,394 |
| Valid Barcodes | 97.50% | 97.90% | 97.10% |
| Sequencing Saturation | 14.80% | 28.60% | 17.80% |
| Q30 Bases in Barcode | 97.70% | 98.00% | 96.90% |
| Q30 Bases in RNA Read | 91.40% | 93.50% | 91.20% |
| Q30 Bases in UMI | 97.60% | 98.00% | 94.20% |
| Reads Mapped to Genome | 93.80% | 96.90% | 89.10% |
| Reads Mapped Confidently to Genome | 91.90% | 92.00% | 86.80% |
| Reads Mapped Confidently to Intergenic Regions | 2.20% | 4.60% | 7.40% |
| Reads Mapped Confidently to Intronic Regions | 2.60% | 3.80% | 5.90% |
| Reads Mapped Confidently to Exonic Regions | 87.10% | 83.70% | 73.50% |
| Reads Mapped Confidently to Transcriptome | 82.20% | 82.00% | 73.00% |
| Reads Mapped Antisense to Gene | 4.40% | 4.10% | 6.00% |
| Fraction Reads in Cells | 67.70% | 58.00% | 16.70% |
| Total Genes Detected | 15,849 | 14,471 | 13,386 |
| Median UMI Counts per Cell | 29,169 | 9,406 | 5,314 |

#### Table S2: List of 198 Genes Used for Classification

| symbol | FlyBase | symbol | FlyBase | symbol | FlyBase | symbol | FlyBase |
|---|---|---|---|---|---|---|---|
| Dip-B | FBgn0000454 | eEF1gamma | FBgn0029176 | CG9861 | FBgn0034844 | CG31029 | FBgn0051029 |
| Hsc70-3 | FBgn0001218 | Tsp42Ee | FBgn0029506 | levy | FBgn0034877 | HEATR2 | FBgn0051320 |
| Hsp83 | FBgn0001233 | CG16781 | FBgn0029661 | CG13585 | FBgn0035020 | CG31642 | FBgn0051642 |
| spir | FBgn0003475 | CG12680 | FBgn0029740 | mri | FBgn0035107 | COX6CL | FBgn0051644 |
| sta | FBgn0003517 | MCTS1 | FBgn0029833 | CG13901 | FBgn0035164 | CG31784 | FBgn0051784 |
| mts | FBgn0004177 | CG3566 | FBgn0029854 | CG2211 | FBgn0035211 | SCaMC | FBgn0052103 |
| HmgD | FBgn0004362 | CG11369 | FBgn0029951 | CG12091 | FBgn0035228 | CG32181 | FBgn0052181 |
| aly | FBgn0004372 | CG15330 | FBgn0029987 | CG13926 | FBgn0035243 | Naa30B | FBgn0052319 |
| B52 | FBgn0004587 | CG15337 | FBgn0030014 | Vta1 | FBgn0035251 | SMSr | FBgn0052380 |
| drk | FBgn0004638 | fh | FBgn0030092 | PAN3 | FBgn0035397 | CG32720 | FBgn0052720 |
| Polr2I | FBgn0004855 | CG2533 | FBgn0030319 | CG1309 | FBgn0035519 | CG31275 | FBgn0063261 |
| Calr | FBgn0005585 | CG12096 | FBgn0030457 | CG10674 | FBgn0035592 | Cby | FBgn0067317 |
| PpD5 | FBgn0005778 | CG10996 | FBgn0030525 | CG6610 | FBgn0035675 | CG34173 | FBgn0085202 |
| Sod2 | FBgn0010213 | Slc25A46a | FBgn0030717 | Sh3beta | FBgn0035772 | Not1 | FBgn0085436 |
| GstS1 | FBgn0010226 | rngo | FBgn0030753 | CG8111 | FBgn0035825 | CG34450 | FBgn0085479 |
| heph | FBgn0011224 | stas | FBgn0030850 | CG5021 | FBgn0035944 | 26-29-p | FBgn0250848 |
| Trp1 | FBgn0011584 | ND-24 | FBgn0030853 | CG4483 | FBgn0035970 | CG42302 | FBgn0259198 |
| La | FBgn0011638 | CG14229 | FBgn0031059 | CG4080 | FBgn0035983 | gwl | FBgn0260399 |
| lark | FBgn0011640 | CG15450 | FBgn0031132 | CG10748 | FBgn0036327 | Ero1L | FBgn0261274 |
| Snr1 | FBgn0011715 | CG15880 | FBgn0031283 | Best4 | FBgn0036491 | slim | FBgn0261477 |
| tsr | FBgn0011726 | CG7295 | FBgn0031372 | QIL1 | FBgn0036726 | SmD2 | FBgn0261789 |
| TfIIA-S | FBgn0013347 | CG8851 | FBgn0031546 | CG8004 | FBgn0036920 | Vha16-4 | FBgn0262513 |
| alien | FBgn0013746 | Art2 | FBgn0031592 | CG3288 | FBgn0037030 | fzr | FBgn0262699 |
| Cp1 | FBgn0013770 | CG15432 | FBgn0031603 | Vps37B | FBgn0037299 | twr | FBgn0262801 |
| Fkbp12 | FBgn0013954 | CG3792 | FBgn0031662 | CG11999 | FBgn0037312 | CG43188 | FBgn0262817 |
| CkIalpha | FBgn0015024 | tomb | FBgn0031715 | CG14668 | FBgn0037320 | CG43317 | FBgn0263022 |
| Rpt2 | FBgn0015282 | cdc14 | FBgn0031952 | Hpr1 | FBgn0037382 | Rcd5 | FBgn0263832 |
| Ssb-c31a | FBgn0015299 | Argl | FBgn0032076 | wa-cup | FBgn0037502 | Sgt1 | FBgn0265101 |
| BEAF-32 | FBgn0015602 | CG9586 | FBgn0032101 | Son | FBgn0037716 | FeCH | FBgn0266268 |
| nrv2 | FBgn0015777 | w-cup | FBgn0032269 | CG8478 | FBgn0037746 | Zmynd10 | FBgn0266709 |
| apt | FBgn0015903 | CG5421 | FBgn0032434 | topi | FBgn0037751 | CG45263 | FBgn0266801 |
| Nurf-38 | FBgn0016687 | CCT4 | FBgn0032444 | CG12817 | FBgn0037798 | SelR | FBgn0267376 |
| Usp47 | FBgn0016756 | CG15482 | FBgn0032483 | fabp | FBgn0037913 | vib | FBgn0267975 |
| CG5989 | FBgn0017429 | Tpr2 | FBgn0032586 | CG14864 | FBgn0038311 | eEF1alpha1 | FBgn0284245 |
| Lk6 | FBgn0017581 | Prosbeta4 | FBgn0032596 | CG5478 | FBgn0038386 | Pdi | FBgn0286818 |
| ATPsyngamma | FBgn0020235 | CG6380 | FBgn0032632 | blp | FBgn0038387 | dbf | FBgn0287630 |
| Rack1 | FBgn0020618 | CG15142 | FBgn0032645 | m-cup | FBgn0038488 | | |
| eIF4B | FBgn0020660 | CG9328 | FBgn0032886 | Prx3 | FBgn0038519 | | |
| Pka-R2 | FBgn0022382 | CG13751 | FBgn0033340 | CG14315 | FBgn0038568 | | |
| EloB | FBgn0023212 | CG2063 | FBgn0033400 | CG10887 | FBgn0038773 | | |
| Rtca | FBgn0025630 | Urod | FBgn0033428 | VhaAC39-2 | FBgn0039058 | | |
| SkpA | FBgn0025637 | wuc | FBgn0033770 | Spase22-23 | FBgn0039172 | | |
| scf | FBgn0025682 | Pex13 | FBgn0033812 | CG6980 | FBgn0039228 | | |
| tna | FBgn0026160 | mars | FBgn0033845 | CG14546 | FBgn0039395 | | |
| tacc | FBgn0026620 | CG13334 | FBgn0033856 | Gp93 | FBgn0039562 | | |
| Kap-alpha3 | FBgn0027338 | CG6220 | FBgn0033865 | Nph | FBgn0039735 | | |
| CG7115 | FBgn0027515 | Tfb1 | FBgn0033929 | ATPsynC | FBgn0039830 | | |
| ATPsyndelta | FBgn0028342 | CG11807 | FBgn0033996 | CG12061 | FBgn0040031 | | |
| Rpt5 | FBgn0028684 | Parp16 | FBgn0034129 | schlank | FBgn0040918 | | |
| Rpn2 | FBgn0028692 | CG8963 | FBgn0034181 | wrd | FBgn0042693 | | |
| Fmr1 | FBgn0028734 | Dnaaf3 | FBgn0034352 | HBS1 | FBgn0042712 | | |
| CSN6 | FBgn0028837 | CG18605 | FBgn0034411 | AP-2sigma | FBgn0043012 | | |
| cathD | FBgn0029093 | CG13526 | FBgn0034774 | Cka | FBgn0044323 | | |
| Prosbeta5 | FBgn0029134 | LS2 | FBgn0034834 | CG30398 | FBgn0050398 | | |

#### Table S3: Gene Ontology Analysis for 198-Gene List

| GO biological process | minor category | # | # | expected | Fold | sign | raw P value | FDR |
|---|---|---|---|---|---|---|---|---|
| motile cilium assembly | | 27 | 5 | 0.39 | 12.93 | + | 7.89E-05 | 2.95E-02 |
| | cellular process | 7050 | 131 | 100.98 | 1.3 | + | 2.15E-05 | 2.68E-02 |
| axonemal dynein complex assembly | | 30 | 5 | 0.43 | 11.64 | + | 1.23E-04 | 3.83E-02 |
| | axoneme assembly | 60 | 7 | 0.86 | 8.14 | + | 4.28E-05 | 2.00E-02 |
| | microtubule bundle formation | 72 | 7 | 1.03 | 6.79 | + | 1.24E-04 | 3.70E-02 |
| | microtubule cytoskeleton organization | 350 | 20 | 5.01 | 3.99 | + | 2.88E-07 | 7.19E-04 |
| | cytoskeleton organization | 588 | 22 | 8.42 | 2.61 | + | 5.16E-05 | 2.27E-02 |
| | microtubule-based process | 477 | 24 | 6.83 | 3.51 | + | 1.66E-07 | 1.24E-03 |
| | protein-containing complex organization | 731 | 26 | 10.47 | 2.48 | + | 2.38E-05 | 2.55E-02 |
| regulation of cellular response to insulin stimulus | | 46 | 6 | 0.66 | 9.11 | + | 8.72E-05 | 2.96E-02 |
| regulation of insulin receptor signaling pathway | | 48 | 6 | 0.69 | 8.73 | + | 1.08E-04 | 3.51E-02 |
| centrosome cycle | | 74 | 7 | 1.06 | 6.6 | + | 1.45E-04 | 3.87E-02 |
| | microtubule organizing center organization | 79 | 7 | 1.13 | 6.19 | + | 2.11E-04 | 4.93E-02 |
| cilium movement | | 78 | 7 | 1.12 | 6.27 | + | 1.96E-04 | 4.73E-02 |
| positive regulation of protein metabolic process | | 189 | 12 | 2.71 | 4.43 | + | 2.85E-05 | 2.37E-02 |
| | regulation of protein metabolic process | 419 | 22 | 6 | 3.67 | + | 2.84E-07 | 1.06E-03 |
| | regulation of primary metabolic process | 1708 | 44 | 24.47 | 1.8 | + | 1.24E-04 | 3.57E-02 |
| protein maturation | | 248 | 13 | 3.55 | 3.66 | + | 8.68E-05 | 3.09E-02 |
| | protein metabolic process | 2213 | 55 | 31.7 | 1.74 | + | 3.57E-05 | 2.23E-02 |
| | macromolecule metabolic process | 3241 | 73 | 46.42 | 1.57 | + | 3.09E-05 | 2.10E-02 |
| | organic substance metabolic process | 4341 | 90 | 62.18 | 1.45 | + | 4.08E-05 | 2.18E-02 |
| | metabolic process | 4623 | 96 | 66.22 | 1.45 | + | 1.42E-05 | 2.66E-02 |
| | organonitrogen compound metabolic process | 2705 | 64 | 38.75 | 1.65 | + | 2.90E-05 | 2.17E-02 |
| | nitrogen compound metabolic process | 3687 | 81 | 52.81 | 1.53 | + | 1.63E-05 | 2.45E-02 |
| | primary metabolic process | 4094 | 84 | 58.64 | 1.43 | + | 1.58E-04 | 4.08E-02 |
| spermatogenesis | | 273 | 13 | 3.91 | 3.32 | + | 2.15E-04 | 4.88E-02 |
| | male gamete generation | 315 | 15 | 4.51 | 3.32 | + | 7.04E-05 | 2.77E-02 |
| | gamete generation | 872 | 29 | 12.49 | 2.32 | + | 2.77E-05 | 2.59E-02 |
| | multicellular organismal reproductive process | 895 | 29 | 12.82 | 2.26 | + | 6.00E-05 | 2.49E-02 |
| proteolysis involved in protein catabolic process | | 334 | 15 | 4.78 | 3.14 | + | 1.31E-04 | 3.64E-02 |
| | protein catabolic process | 346 | 15 | 4.96 | 3.03 | + | 1.90E-04 | 4.74E-02 |
| cellular process involved in reproduction in multicellular organism | | 846 | 28 | 12.12 | 2.31 | + | 4.21E-05 | 2.10E-02 |

#### Table S4: Differential Expression Analysis and Ranking for *D. melanogaster* Marker Genes

| Gene | Somatic | GSC/E. -gonia | L. -gonia | *ananassae* -cyte | E. -cyte | L. -cyte | E. -tid | L. -tid |
|---|---|---|---|---|---|---|---|---|
| aly | N/A | 701/1326 (+) | 576/960 (+) | N/A | N/A | N/A | 137/713 (-) | 128/639 (-) |
| zfh1 | N/A | N/A | N/A | N/A | N/A | N/A | N/A | N/A |
| Fas1 | N/A | N/A | N/A | N/A | N/A | N/A | N/A | N/A |
| Hsp23 | 7/827 (+) | 1198/1326 (-) | N/A | N/A | 254/254 (-) | 429/439 (-) | 675/913 (-) | 547/639 (-) |
| MtnA | 4/827 (+) | N/A | N/A | N/A | N/A | 435/439 (-) | 713/713 (-) | 639/639 (-) |
| His2Av | 517/827 (-) | 375/1326 (+) | 455/960 (+) | 245/366 (+) | N/A | N/A | 293/713 (-) | 380/639 (-) |
| aub | N/A | 600/1326 (+) | N/A | N/A | N/A | N/A | N/A | N/A |
| bam | N/A | 766/1326 (+) | N/A | N/A | N/A | N/A | N/A | N/A |
| fzo | N/A | N/A | N/A | N/A | N/A | N/A | N/A | N/A |
| twe | 657/827 (-) | N/A | N/A | N/A | N/A | N/A | 163/713 (-) | 254/639 (-) |
| soti | 810/827 (-) | 1308/1326 (-) | N/A | N/A | N/A | N/A | 634/713 (-) | 24/639 (+) |
| Dpy-30L2 | N/A | N/A | N/A | N/A | N/A | N/A | N/A | N/A |
| p-cup | 724/827 (-) | 1183/1326 (-) | N/A | 73/366 (+) | 169/254 (-) | N/A | N/A | 18/639 (+) |

E. = Early, L. = Late
"-" = "spermato-"
N/A = gene was not significantly differentially expressed in given cell type
(+) = upregulated, (-) = downregulated

#### Table S5: Differentially Expressed Genes in "*ananassae* spermatocyte"

| p_val | avg_log2FC | pct.1 | pct.2 | p_val_adj | cluster | gene |
|---|---|---|---|---|---|---|
| 4.50E-53 | 1.409943754 | 0.985 | 0.939 | 9.00E-50 | "ananassae" spermatocyte | Pzl |
| 0.000270958 | 1.358382077 | 0.483 | 0.187 | 0.541915229 | "ananassae" spermatocyte | CG1674 |
| 4.50E-63 | 1.210942688 | 0.73 | 0.565 | 9.00E-60 | "ananassae" spermatocyte | Gp93 |
| 1.53E-86 | 1.201585427 | 0.99 | 0.773 | 3.06E-83 | "ananassae" spermatocyte | Prosbeta2R2 |
| 1.01E-84 | 1.185955458 | 0.995 | 0.868 | 2.02E-81 | "ananassae" spermatocyte | Kr-h2 |
| 4.77E-53 | 1.167950305 | 0.655 | 0.175 | 9.55E-50 | "ananassae" spermatocyte | Pkd2 |
| 1.24E-88 | 1.152438016 | 0.923 | 0.887 | 2.49E-85 | "ananassae" spermatocyte | Tob |
| 8.82E-49 | 1.145990174 | 0.682 | 0.309 | 1.76E-45 | "ananassae" spermatocyte | CG5421 |
| 1.17E-20 | 1.122626694 | 0.523 | 0.197 | 2.35E-17 | "ananassae" spermatocyte | CG31999 |
| 2.92E-12 | 1.112658973 | 0.565 | 0.508 | 5.83E-09 | "ananassae" spermatocyte | pigs |
| 5.25E-14 | 1.064575532 | 0.653 | 0.686 | 1.05E-10 | "ananassae" spermatocyte | CG1428 |
| 0.000359456 | 1.052631057 | 0.63 | 0.778 | 0.718911381 | "ananassae" spermatocyte | sls |
| 5.76E-125 | 1.047015777 | 0.92 | 0.759 | 1.15E-121 | "ananassae" spermatocyte | Hsc70-3 |
| 1.44E-71 | 0.979132548 | 1 | 0.964 | 2.88E-68 | "ananassae" spermatocyte | CG31609 |
| 3.86E-72 | 0.931736647 | 0.897 | 0.641 | 7.71E-69 | "ananassae" spermatocyte | Best3 |
| 5.11E-60 | 0.915974674 | 0.993 | 0.775 | 1.02E-56 | "ananassae" spermatocyte | VhaPPA1-2 |
| 1.58E-66 | 0.908834704 | 0.938 | 0.715 | 3.16E-63 | "ananassae" spermatocyte | CG31465 |
| 2.91E-20 | 0.905885198 | 0.668 | 0.514 | 5.81E-17 | "ananassae" spermatocyte | Best4 |
| 2.63E-43 | 0.894541377 | 0.745 | 0.598 | 5.26E-40 | "ananassae" spermatocyte | Sc2 |

#### Table S6: Number of Genes by Gene Set

| Gene Set | Somatic | GSC/E. -gonia | L. -gonia | ananassae -cyte | E. -cyte | L. -cyte | E. -tid | L. -tid |
|---|---|---|---|---|---|---|---|---|
| mel_L2 | 38 | 57 | 57 | 45 | 11 | 10 | 26 | 32 |
| mel_L1 | 48 | 55 | 56 | 52 | 21 | 16 | 33 | 35 |
| mel_R1 | 36 | 43 | 50 | 42 | 10 | 8 | 21 | 25 |
| mel_R2 | 43 | 64 | 54 | 48 | 18 | 15 | 30 | 31 |
| yak_L2 | 35 | 64 | 82 | 67 | 36 | 14 | 33 | 30 |
| yak_L1 | 44 | 73 | 84 | 61 | 41 | 19 | 35 | 41 |
| yak_R1 | 23 | 54 | 71 | 52 | 35 | 11 | 21 | 23 |
| yak_R2 | 35 | 69 | 83 | 68 | 47 | 21 | 34 | 39 |
| ana_L2 | 9 | 22 | 39 | 11 | 4 | 17 | 20 | 7 |
| ana_L1 | 12 | 26 | 36 | 20 | 3 | 21 | 24 | 6 |
| ana_R1 | 12 | 11 | 27 | 10 | 1 | 10 | 18 | 3 |
| ana_R2 | 13 | 16 | 41 | 18 | 4 | 20 | 28 | 8 |
| mel_autosome | 2532 | 4247 | 3735 | 3214 | 933 | 695 | 1478 | 1859 |
| yak_autosome | 2317 | 5838 | 6263 | 4502 | 2717 | 903 | 1823 | 2209 |
| ana_autosome | 848 | 1625 | 3471 | 1029 | 178 | 1197 | 1606 | 373 |
| mel_chrx | 370 | 667 | 533 | 428 | 84 | 64 | 184 | 236 |
| yak_chrx | 344 | 1088 | 1062 | 716 | 398 | 120 | 276 | 345 |
| ana_chrx | 138 | 247 | 534 | 150 | 11 | 197 | 257 | 59 |

E. = Early, L. = Late
"-" = "spermato-"

#### Table S7: Raw P-Values for Differences in Mean Read Count per Cell

| Gene Set | Somatic | GSC/E. -gonia | L. -gonia | ananassae -cyte | E. -cyte | L. -cyte | E. -tid | L. -tid |
|---|---|---|---|---|---|---|---|---|
| mel_L2 | 0.12948854 | 0.89941216 | 2.64E-11 | 1.78E-12 | 1.01E-07 | 2.58E-10 | 0.15146514 | 0.08912165 |
| yak_L2 | 5.90E-10 | 0.847824101 | 0.182225266 | 1.06E-17 | 6.87E-08 | 6.23E-52 | 4.06E-07 | 0.603624623 |
| ana_L2 | 3.39E-11 | 3.82E-06 | 0.525018863 | 0.000230883 | 6.14E-10 | 0.767278592 | 0.365704743 | 1.02E-05 |
| mel_L1 | 0.189765282 | 0.444395722 | 2.27E-10 | 3.54E-13 | 7.19E-30 | 4.40E-29 | 0.011799289 | 0.816066834 |
| yak_L1 | 0.50720622 | 0.954235869 | 0.050956853 | 1.84E-17 | 1.33E-16 | 2.24E-09 | 0.083588278 | 0.855485187 |
| ana_L1 | 0.000308698 | 0.132501222 | 0.016986126 | 0.090125617 | 2.87E-209 | 0.292481269 | 0.319613352 | 6.57E-18 |
| mel_R1 | 0.007969019 | 1.02E-07 | 3.16E-119 | 7.20E-63 | 7.14E-68 | 9.36E-107 | 0.000976485 | 0.005802494 |
| yak_R1 | 5.07E-07 | 0.16399208 | 0.001346342 | 3.50E-16 | 1.90E-38 | 3.17E-81 | 1.37E-10 | 0.255292677 |
| ana_R1 | 2.92E-18 | 0.032799135 | 0.478570304 | 0.030790244 | 3.05E-39 | 0.632634517 | 0.356834788 | 1.46E-09 |
| mel_R2 | 0.896154945 | 0.002076184 | 1.20E-25 | 9.57E-92 | 0.060221471 | 1.78E-12 | 0.760707565 | 0.938006163 |
| yak_R2 | 1.52E-06 | 0.348065129 | 0.059120936 | 2.97E-07 | 2.99E-05 | 0.237679293 | 0.451404728 | 0.89398261 |
| ana_R2 | 0.00701831 | 0.137090851 | 0.009196743 | 4.18E-11 | 0.603993707 | 0.35187094 | 0.18668129 | 0.330831085 |

E. = Early, L. = Late
"-" = "spermato-"

#### Table S8: Hybridization Chain Reaction Probe Sequences

| spp. | gene | probe | sequence |
|---|---|---|---|
| mel | rbp4 | B1-Rbp4_RA_16_1A | GAGGAGGGCAGCAAACGGAAATGCTTTATTATTTATTAAATATCT |
| mel | rbp4 | B1-Rbp4_RA_16_2A | GAGGAGGGCAGCAAACGGAACTTAGATACCGTAGGGTGGTGTTGT |
| mel | rbp4 | B1-Rbp4_RA_16_3A | GAGGAGGGCAGCAAACGGAATTAATAGCTCCAACTCCCACGGCGG |
| mel | rbp4 | B1-Rbp4_RA_16_4A | GAGGAGGGCAGCAAACGGAAAGTAGCTGCCTGGAGTTCTTTCAAA |
| mel | rbp4 | B1-Rbp4_RA_16_5A | GAGGAGGGCAGCAAACGGAAACTCCTGGGCCGAGTAGATACCGGT |
| mel | rbp4 | B1-Rbp4_RA_16_6A | GAGGAGGGCAGCAAACGGAATTCTGTCCGGGTGTTGGCTTGGGTG |
| mel | rbp4 | B1-Rbp4_RA_16_7A | GAGGAGGGCAGCAAACGGAAATTGTTGTAATTGTAGCTATCAGCA |
| mel | rbp4 | B1-Rbp4_RA_16_8A | GAGGAGGGCAGCAAACGGAATCTTCACCTCCACCAATGTCTGAAG |
| mel | rbp4 | B1-Rbp4_RA_16_9A | GAGGAGGGCAGCAAACGGAAGTCTCCCTGTCCATGAGCAATTTCA |
| mel | rbp4 | B1-Rbp4_RA_16_10A | GAGGAGGGCAGCAAACGGAATTTCGAGTTCATGAAACCGGCTTCA |
| mel | rbp4 | B1-Rbp4_RA_16_11A | GAGGAGGGCAGCAAACGGAACCGCCACCCCTTTTGAAATCTTGAC |
| mel | rbp4 | B1-Rbp4_RA_16_12A | GAGGAGGGCAGCAAACGGAATATTATCAATGGTGTGGGGACGTGC |
| mel | rbp4 | B1-Rbp4_RA_16_13A | GAGGAGGGCAGCAAACGGAACACATAGGTGACGAAGCCAAAGCCC |
| mel | rbp4 | B1-Rbp4_RA_16_14A | GAGGAGGGCAGCAAACGGAAACCACCGCATCGGCCACGATACCGA |
| mel | rbp4 | B1-Rbp4_RA_16_15A | GAGGAGGGCAGCAAACGGAACGGTCGTCTGAGTGGATAGGCCGCC |
| mel | rbp4 | B1-Rbp4_RA_16_16A | GAGGAGGGCAGCAAACGGAATTCTTCCTTAATGGCCGTCTTTCCG |
| mel | rbp4 | B1-Rbp4_RA_16_1B | ACGATTTTAAGTCTCAAAAGGACCCTAGAAGAGTCTTCCTTTACG |
| mel | rbp4 | B1-Rbp4_RA_16_2B | GGTTTAACTGCGTTCCATCGTCCGCTAGAAGAGTCTTCCTTTACG |
| mel | rbp4 | B1-Rbp4_RA_16_3B | ACCTTATAGTCCTGGGTGGGCCACTTAGAAGAGTCTTCCTTTACG |
| mel | rbp4 | B1-Rbp4_RA_16_4B | GGCGCCGAAATCCAGCTTGTTATTGTAGAAGAGTCTTCCTTTACG |
| mel | rbp4 | B1-Rbp4_RA_16_5B | CCCATTCGGCCACTTTGCTGGAAGTTAGAAGAGTCTTCCTTTACG |
| mel | rbp4 | B1-Rbp4_RA_16_6B | TGCTGTGACGGCAACGAAGCAGCCATAGAAGAGTCTTCCTTTACG |
| mel | rbp4 | B1-Rbp4_RA_16_7B | TTGGGCCAGATAGGGATTGTAGTTGTAGAAGAGTCTTCCTTTACG |
| mel | rbp4 | B1-Rbp4_RA_16_8B | AACGCCTGTCGGCCTTCTGGGTGGATAGAAGAGTCTTCCTTTACG |
| mel | rbp4 | B1-Rbp4_RA_16_9B | AGGAAGCCAAATGCCCGCTGACGTCTAGAAGAGTCTTCCTTTACG |
| mel | rbp4 | B1-Rbp4_RA_16_10B | TTCCTTCAAGCCGCCCAGAAAGATCTAGAAGAGTCTTCCTTTACG |
| mel | rbp4 | B1-Rbp4_RA_16_11B | CCAAAACTTCCGACAACGGAACCAATAGAAGAGTCTTCCTTTACG |
| mel | rbp4 | B1-Rbp4_RA_16_12B | GCAGTGCGTGCTTGGTCTCCACAATTAGAAGAGTCTTCCTTTACG |
| mel | rbp4 | B1-Rbp4_RA_16_13B | TTGGACAATTTCCACGGACTTGGGATAGAAGAGTCTTCCTTTACG |
| mel | rbp4 | B1-Rbp4_RA_16_14B | GAATGGTTGCTCACCGGATCCCGCATAGAAGAGTCTTCCTTTACG |
| mel | rbp4 | B1-Rbp4_RA_16_15B | GACTAAAGAATCCGCGCAGCGTTTCTAGAAGAGTCTTCCTTTACG |
| mel | rbp4 | B1-Rbp4_RA_16_16B | GAAGATCTTGCGCAAATGCTCGTAGTAGAAGAGTCTTCCTTTACG |
| mel | soti | B3-soti_RA_16_1A | GTCCCTGCCTCTATATCTTTTATATGTGGTACTTATATATTATGT |
| mel | soti | B3-soti_RA_16_2A | GTCCCTGCCTCTATATCTTTAAAACAATAAGTTGAAAACCAAATA |
| mel | soti | B3-soti_RA_16_3A | GTCCCTGCCTCTATATCTTTTTATTGAACAAGATTGAACCGCAAA |
| mel | soti | B3-soti_RA_16_4A | GTCCCTGCCTCTATATCTTTGTCTTTTTTTCCTTTTTTGAGCTTG |
| mel | soti | B3-soti_RA_16_5A | GTCCCTGCCTCTATATCTTTATTTTTGTTTTGAGCAAGTTTTTAG |
| mel | soti | B3-soti_RA_16_6A | GTCCCTGCCTCTATATCTTTCTGGTTGGATATATGGACGTGTACA |
| mel | soti | B3-soti_RA_16_7A | GTCCCTGCCTCTATATCTTTTGCTCGATCTTCCGAGGTCCAAGAG |
| mel | soti | B3-soti_RA_16_8A | GTCCCTGCCTCTATATCTTTACTATGGCGAATCAGAACTGACCCC |
| mel | soti | B3-soti_RA_16_9A | GTCCCTGCCTCTATATCTTTAGTTCTGACTGGTCATATTCTCGAC |
| mel | soti | B3-soti_RA_16_10A | GTCCCTGCCTCTATATCTTTATCCTCGCGAACGTGACGCATTGCC |
| mel | soti | B3-soti_RA_16_11A | GTCCCTGCCTCTATATCTTTTCCGGCTCCTGACTTTGGCATGGCG |
| mel | soti | B3-soti_RA_16_12A | GTCCCTGCCTCTATATCTTTGTTTCTTCCGCCGACGTGGTGGTCC |
| mel | soti | B3-soti_RA_16_13A | GTCCCTGCCTCTATATCTTTCAACGGTGGCTCTTGAGGAGCGTCC |
| mel | soti | B3-soti_RA_16_14A | GTCCCTGCCTCTATATCTTTAGGCGCTGCTGCTGCTGTCCATCCT |
| mel | soti | B3-soti_RA_16_15A | GTCCCTGCCTCTATATCTTTCATTCACCGCGTCGCCGACTCGATC |
| mel | soti | B3-soti_RA_16_16A | GTCCCTGCCTCTATATCTTTGTGTAGTACGTCCATTCCAACTTAC |
| mel | soti | B3-soti_RA_16_1B | AGATAAATCATTCGGTTTTTATTAATTCCACTCAACTTTAACCCG |
| mel | soti | B3-soti_RA_16_2B | TTGTTAACAATTGTAGAATAAATTTTTCCACTCAACTTTAACCCG |
| mel | soti | B3-soti_RA_16_3B | AAATATAAGTTTGCTAAATCGTGTTTTCCACTCAACTTTAACCCG |
| mel | soti | B3-soti_RA_16_4B | TTTTCAACTGTAAGTCAGTAAAGTATTCCACTCAACTTTAACCCG |
| mel | soti | B3-soti_RA_16_5B | GAGCGGTAGATAACTATGGTGTAACTTCCACTCAACTTTAACCCG |
| mel | soti | B3-soti_RA_16_6B | AAAGATAACTAATGGGATGCATATCTTCCACTCAACTTTAACCCG |
| mel | soti | B3-soti_RA_16_7B | ATGTGTAGTATGGGATCCCAATGAGTTCCACTCAACTTTAACCCG |
| mel | soti | B3-soti_RA_16_8B | TTCTTCTAATTTACAATCTATGGGATTCCACTCAACTTTAACCCG |
| mel | soti | B3-soti_RA_16_9B | GCTGTGGCAGACCCATACCATTCGGTTCCACTCAACTTTAACCCG |
| mel | soti | B3-soti_RA_16_10B | GTAATTTGCAAAGTACTCGCCTCGCTTCCACTCAACTTTAACCCG |

| | | | |
|---|---|---|---|
| mel | soti | B3-soti_RA_16_11B | GTGACACTGGCCATTAGGAGGCACATTCCACTCAACTTTAACCCG |
| mel | soti | B3-soti_RA_16_12B | TGGGTTTGGTCATAGTGTAGAAACTTTCCACTCAACTTTAACCCG |
| mel | soti | B3-soti_RA_16_13B | TGGCCCTCCTTGCGCCGGAAAAACGTTCCACTCAACTTTAACCCG |
| mel | soti | B3-soti_RA_16_14B | AGTATGTCCATCTGACGATTGACTCTTCCACTCAACTTTAACCCG |
| mel | soti | B3-soti_RA_16_15B | ATGTGTCCAAGTCATCGCCAGCATCTTCCACTCAACTTTAACCCG |
| mel | soti | B3-soti_RA_16_16B | TAGCTGCTCATCGTACAGATCGTGTTTCCACTCAACTTTAACCCG |
| mel | B52 | B4-B52_RA_16_1A | CCTCAACCTACCTCCAACAAAAATTACTCATTGTATAGACAATTT |
| mel | B52 | B4-B52_RA_16_2A | CCTCAACCTACCTCCAACAATCATCTTCCAGCCAATGTCCCATGT |
| mel | B52 | B4-B52_RA_16_3A | CCTCAACCTACCTCCAACAAATTCATTAAGTTTTGGTTGTAGTCG |
| mel | B52 | B4-B52_RA_16_4A | CCTCAACCTACCTCCAACAATCTAATCGTCCATGCTCTCGTTATT |
| mel | B52 | B4-B52_RA_16_5A | CCTCAACCTACCTCCAACAACGATCGCGTGAACGGGAGTCGCGGT |
| mel | B52 | B4-B52_RA_16_6A | CCTCAACCTACCTCCAACAACGAACGGGAGCGGGTGCGGGAGTGG |
| mel | B52 | B4-B52_RA_16_7A | CCTCAACCTACCTCCAACAATGGAGCGTCCGCCACGAGATTTGGA |
| mel | B52 | B4-B52_RA_16_8A | CCTCAACCTACCTCCAACAAGAACGGGAGCGTCCACGTCCGCTGC |
| mel | B52 | B4-B52_RA_16_9A | CCTCAACCTACCTCCAACAAAATGGCCGTCTTCATGTCCGACAAC |
| mel | B52 | B4-B52_RA_16_10A | CCTCAACCTACCTCCAACAATGAGATCCTGCCAGCTAACGCGGCT |
| mel | B52 | B4-B52_RA_16_11A | CCTCAACCTACCTCCAACAACCGCCCCCCCGCCGACCACCATATC |
| mel | B52 | B4-B52_RA_16_12A | CCTCAACCTACCTCCAACAAGAGCGGTACCCCTGGCGGGTTCAAC |
| mel | B52 | B4-B52_RA_16_13A | CCTCAACCTACCTCCAACAAATTCAGTTCATAGACGGCATCGTCG |
| mel | B52 | B4-B52_RA_16_14A | CCTCAACCTACCTCCAACAACCGTAGCCATTTTTGATGAGGATGT |
| mel | B52 | B4-B52_RA_16_15A | CCTCAACCTACCTCCAACAAGCTCCAAATCGCGCTCGCGCACTCC |
| mel | B52 | B4-B52_RA_16_16A | CCTCAACCTACCTCCAACAATCCCACCATGATAACGGTTCCTTAC |
| mel | B52 | B4-B52_RA_16_1B | ATAGTTTAAATTTATCCGTATTTATATTCTCACCATATTCGCTTC |
| mel | B52 | B4-B52_RA_16_2B | CAAATCGGCACATTCAGCCGATCCCATTCTCACCATATTCGCTTC |
| mel | B52 | B4-B52_RA_16_3B | GACAGTGGTGGATTGGTATGATTTCATTCTCACCATATTCGCTTC |
| mel | B52 | B4-B52_RA_16_4B | TTTAATATTAATGGGACAGCTAACTATTCTCACCATATTCGCTTC |
| mel | B52 | B4-B52_RA_16_5B | CGCGACTTGTTTTCAGCCGATGCGGATTCTCACCATATTCGCTTC |
| mel | B52 | B4-B52_RA_16_6B | GGAACGGGAGTCACGCTCACGTTTGATTCTCACCATATTCGCTTC |
| mel | B52 | B4-B52_RA_16_7B | AACGAGACTTGACTGGCGACTTGGAATTCTCACCATATTCGCTTC |
| mel | B52 | B4-B52_RA_16_8B | GATCGGGAGCGCGATCGGGAGCTGGATTCTCACCATATTCGCTTC |
| mel | B52 | B4-B52_RA_16_9B | GTTTAGCTCGGTGTCATCCAACTTCATTCTCACCATATTCGCTTC |
| mel | B52 | B4-B52_RA_16_10B | CCTCGCCAGCCTGGCGCATGTAATCATTCTCACCATATTCGCTTC |
| mel | B52 | B4-B52_RA_16_11B | GATGATTTTTCGTTGTAACGACCGCATTCTCACCATATTCGCTTC |
| mel | B52 | B4-B52_RA_16_12B | CGTCGTAGCGGTCGCGGTTGCTGCCATTCTCACCATATTCGCTTC |
| mel | B52 | B4-B52_RA_16_13B | CACACGTTCGCCAAGCAGCTCTTTGATTCTCACCATATTCGCTTC |
| mel | B52 | B4-B52_RA_16_14B | TCACGATAGTCTTCGAATTCCACAAATTCTCACCATATTCGCTTC |
| mel | B52 | B4-B52_RA_16_15B | GTGTGCGGCCGTAGCCTTTGAAAAAATTCTCACCATATTCGCTTC |
| mel | B52 | B4-B52_RA_16_16B | GGGCAGACCGCCCACATACACTCGAATTCTCACCATATTCGCTTC |
| mel | pkd2 | B5-Pkd2_RA_16_1A | CTCACTCCCAATCTCTATAAGGTTCTGTTTCTATTTAGTACTATT |
| mel | pkd2 | B5-Pkd2_RA_16_2A | CTCACTCCCAATCTCTATAACTTTGTCGGTTTTGCTTTAGTTCTC |
| mel | pkd2 | B5-Pkd2_RA_16_3A | CTCACTCCCAATCTCTATAACCTCAAGGAGGCCCACACGGTTATT |
| mel | pkd2 | B5-Pkd2_RA_16_4A | CTCACTCCCAATCTCTATAACATGCCCGACAGTTTCCTGTAGATG |
| mel | pkd2 | B5-Pkd2_RA_16_5A | CTCACTCCCAATCTCTATAAATGGTCAAAATCGAGGTGATGAAGT |
| mel | pkd2 | B5-Pkd2_RA_16_6A | CTCACTCCCAATCTCTATAACCATCATATCGACATAAATAATATT |
| mel | pkd2 | B5-Pkd2_RA_16_7A | CTCACTCCCAATCTCTATAAATAGATGGTGTAGTAGATAACCATT |
| mel | pkd2 | B5-Pkd2_RA_16_8A | CTCACTCCCAATCTCTATAACGATCCAGCCAGTGAATATCCTTCA |
| mel | pkd2 | B5-Pkd2_RA_16_9A | CTCACTCCCAATCTCTATAAAACGGATGAAGGCGTCGTTTACGTA |
| mel | pkd2 | B5-Pkd2_RA_16_10A | CTCACTCCCAATCTCTATAAAGCTCTCCTGTGCCCCCTGAGGTCA |
| mel | pkd2 | B5-Pkd2_RA_16_11A | CTCACTCCCAATCTCTATAACTCAGGTGGCATTGACGTGAGATTC |
| mel | pkd2 | B5-Pkd2_RA_16_12A | CTCACTCCCAATCTCTATAACGTAAATAGTTTCTTCATCGTGTCA |
| mel | pkd2 | B5-Pkd2_RA_16_13A | CTCACTCCCAATCTCTATAACCTATTTTGCAGGGCTGAAGCCGCG |
| mel | pkd2 | B5-Pkd2_RA_16_14A | CTCACTCCCAATCTCTATAACACTTTTGCAGACGTCGATGGTGCT |
| mel | pkd2 | B5-Pkd2_RA_16_15A | CTCACTCCCAATCTCTATAACTTATCAGAAGTCGATTGGCCAGGA |
| mel | pkd2 | B5-Pkd2_RA_16_16A | CTCACTCCCAATCTCTATAACGCTGGCGGTGGCGGAGGTCCTCGA |
| mel | pkd2 | B5-Pkd2_RA_16_1B | ACATTTTATATTTAAATATTTTATAAACTACCCTACAAATCCAAT |
| mel | pkd2 | B5-Pkd2_RA_16_2B | CCAGAGATTAGCGCTCGGTTCTTGTAACTACCCTACAAATCCAAT |
| mel | pkd2 | B5-Pkd2_RA_16_3B | TATTGTTGATCAACTTTTCCAGAATAACTACCCTACAAATCCAAT |
| mel | pkd2 | B5-Pkd2_RA_16_4B | TCGCCCACAATGGGTGATCCAGTAGAACTACCCTACAAATCCAAT |
| mel | pkd2 | B5-Pkd2_RA_16_5B | TGAAAGTCGCCAAGAATCATCCGAAAACTACCCTACAAATCCAAT |
| mel | pkd2 | B5-Pkd2_RA_16_6B | TAATCCACACAAGAAAGGCCAGAATAACTACCCTACAAATCCAAT |
| mel | pkd2 | B5-Pkd2_RA_16_7B | TCCAGATTTGCGAATTTCCGTGATTAACTACCCTACAAATCCAAT |
| mel | pkd2 | B5-Pkd2_RA_16_8B | AACTCAACCAAACACAGACGCGAGCAACTACCCTACAAATCCAAT |

| | | | |
|---|---|---|---|
| mel | pkd2 | B5-Pkd2_RA_16_9B | AATATGCAGCATAGCAGGTGTTAAAAACTACCCTACAAATCCAAT |
| mel | pkd2 | B5-Pkd2_RA_16_10B | GTTCCCATCCACACCACTCTGTCGCAACTACCCTACAAATCCAAT |
| mel | pkd2 | B5-Pkd2_RA_16_11B | ACCCTGCTCACTACCCTGCTCACTAAACTACCCTACAAATCCAAT |
| mel | pkd2 | B5-Pkd2_RA_16_12B | CGAAGGAGCCACTACCATTTCCCGAAACTACCCTACAAATCCAAT |
| mel | pkd2 | B5-Pkd2_RA_16_13B | TGTGTAAAGGATGCGATTGGTTTTGAACTACCCTACAAATCCAAT |
| mel | pkd2 | B5-Pkd2_RA_16_14B | TCGAGGTTCAGATGCAGCTAGCTTGAACTACCCTACAAATCCAAT |
| mel | pkd2 | B5-Pkd2_RA_16_15B | AGTCGATGGCTTATCAGAAGTCGATAACTACCCTACAAATCCAAT |
| mel | pkd2 | B5-Pkd2_RA_16_16B | ACCCACTCCTATTGAAACGCGTTTGAACTACCCTACAAATCCAAT |
| ana | rbp4 | B1-ana_Rbp4_RA_GF17369_16_1A | GAGGAGGGCAGCAAACGGAATGGGTCGATTTATTTCTTCCAAGGA |
| ana | rbp4 | B1-ana_Rbp4_RA_GF17369_16_2A | GAGGAGGGCAGCAAACGGAATGAATTAAATATATATGTCATGTAT |
| ana | rbp4 | B1-ana_Rbp4_RA_GF17369_16_3A | GAGGAGGGCAGCAAACGGAACTATCATTTGTTATTTGAACGAAAA |
| ana | rbp4 | B1-ana_Rbp4_RA_GF17369_16_4A | GAGGAGGGCAGCAAACGGAATATCAAGGTGGTATCAGGGTGGTGT |
| ana | rbp4 | B1-ana_Rbp4_RA_GF17369_16_5A | GAGGAGGGCAGCAAACGGAAGGGCAAGGTGCCTCCGTTTCCCACT |
| ana | rbp4 | B1-ana_Rbp4_RA_GF17369_16_6A | GAGGAGGGCAGCAAACGGAAGCACAACAGTTCCTGCCCCATCTGC |
| ana | rbp4 | B1-ana_Rbp4_RA_GF17369_16_7A | GAGGAGGGCAGCAAACGGAACATTCACCCACATTGCTCGGAGTCC |
| ana | rbp4 | B1-ana_Rbp4_RA_GF17369_16_8A | GAGGAGGGCAGCAAACGGAACTGATGGAGATTCTTGGCGGCGTAG |
| ana | rbp4 | B1-ana_Rbp4_RA_GF17369_16_9A | GAGGAGGGCAGCAAACGGAAGCGCCAGGTAGGGATTGTAGGTGCT |
| ana | rbp4 | B1-ana_Rbp4_RA_GF17369_16_10A | GAGGAGGGCAGCAAACGGAACGTTTGCCGGGATCTGGTTTCTGGG |
| ana | rbp4 | B1-ana_Rbp4_RA_GF17369_16_11A | GAGGAGGGCAGCAAACGGAACTCGAGAAACCCGAATCCCCGCTTG |
| ana | rbp4 | B1-ana_Rbp4_RA_GF17369_16_12A | GAGGAGGGCAGCAAACGGAAGACAGTCCTTCAGTCCGCCCAGAAA |
| ana | rbp4 | B1-ana_Rbp4_RA_GF17369_16_13A | GAGGAGGGCAGCAAACGGAAGCCCGTTTGGTTTCCACGGTTTTGT |
| ana | rbp4 | B1-ana_Rbp4_RA_GF17369_16_14A | GAGGAGGGCAGCAAACGGAAATTGGTCATCGGATCTCGCATCACC |
| ana | rbp4 | B1-ana_Rbp4_RA_GF17369_16_15A | GAGGAGGGCAGCAAACGGAATTTTGCGCAAGTGCTCGTCTCCTTC |
| ana | rbp4 | B1-ana_Rbp4_RA_GF17369_16_16A | GAGGAGGGCAGCAAACGGAAAACTGGCATGACCCTCCGGCCGCAC |
| ana | rbp4 | B1-ana_Rbp4_RA_GF17369_16_1B | TTTTCATGCAAATTTCCTATGCAAATAGAAGAGTCTTCCTTTACG |
| ana | rbp4 | B1-ana_Rbp4_RA_GF17369_16_2B | TTCGAATAACTATGATTGCTTTTGCTAGAAGAGTCTTCCTTTACG |
| ana | rbp4 | B1-ana_Rbp4_RA_GF17369_16_3B | TAATTTTTTTTTTTAGATCTAAATATAGAAGAGTCTTCCTTTACG |
| ana | rbp4 | B1-ana_Rbp4_RA_GF17369_16_4B | TTGAAATTTTGAATGTTCTTAGAATTAGAAGAGTCTTCCTTTACG |
| ana | rbp4 | B1-ana_Rbp4_RA_GF17369_16_5B | CACCAATTTAAATATCGTACCGTAGTAGAAGAGTCTTCCTTTACG |
| ana | rbp4 | B1-ana_Rbp4_RA_GF17369_16_6B | TGGCCTTGCCGGTGGTCACGGCTACTAGAAGAGTCTTCCTTTACG |
| ana | rbp4 | B1-ana_Rbp4_RA_GF17369_16_7B | GCGTGTTTGTGTCCGGCCTTCGGTGTAGAAGAGTCTTCCTTTACG |
| ana | rbp4 | B1-ana_Rbp4_RA_GF17369_16_8B | ATATCCCAGATGCGGCGGCTGCGGCTAGAAGAGTCTTCCTTTACG |
| ana | rbp4 | B1-ana_Rbp4_RA_GF17369_16_9B | TGAAGGCCGATGGCGGGAGGACGGTTAGAAGAGTCTTCCTTTACG |
| ana | rbp4 | B1-ana_Rbp4_RA_GF17369_16_10B | GCGGCTCCGCCCACGGGGAAGCGAATAGAAGAGTCTTCCTTTACG |
| ana | rbp4 | B1-ana_Rbp4_RA_GF17369_16_11B | CTGGTCAGCGCTAGCAGAATCCTCATAGAAGAGTCTTCCTTTACG |
| ana | rbp4 | B1-ana_Rbp4_RA_GF17369_16_12B | AGTACTCGCGAATGGAGCTCTCGTCTAGAAGAGTCTTCCTTTACG |
| ana | rbp4 | B1-ana_Rbp4_RA_GF17369_16_13B | GGTTTCAAGAATTCGTGACGCGGCATAGAAGAGTCTTCCTTTACG |
| ana | rbp4 | B1-ana_Rbp4_RA_GF17369_16_14B | GGTGACGAAGCCGAAGCCCCGCGAATAGAAGAGTCTTCCTTTACG |
| ana | rbp4 | B1-ana_Rbp4_RA_GF17369_16_15B | TCTGTGTGGATAATCCCCCGATGAATAGAAGAGTCTTCCTTTACG |
| ana | rbp4 | B1-ana_Rbp4_RA_GF17369_16_16B | TTGTTACGGCTCGACGCTTTGCGCTTAGAAGAGTCTTCCTTTACG |
| ana | soti | B3-ana_soti_RA_GF17485_15_1A | GTCCCTGCCTCTATATCTTTATTCTATTCTTCTAATGAAATTAAG |
| ana | soti | B3-ana_soti_RA_GF17485_15_2A | GTCCCTGCCTCTATATCTTTATCTTGGGCCAAAAGAAGCAGCATG |
| ana | soti | B3-ana_soti_RA_GF17485_15_3A | GTCCCTGCCTCTATATCTTTTTCAGGATAGATTGGATCGAGGGGC |
| ana | soti | B3-ana_soti_RA_GF17485_15_4A | GTCCCTGCCTCTATATCTTTTGGAAGATGTGTTCTGGGCCCTGGA |
| ana | soti | B3-ana_soti_RA_GF17485_15_5A | GTCCCTGCCTCTATATCTTTCCCGGCCCGGGGTCCCCGAACCCAT |
| ana | soti | B3-ana_soti_RA_GF17485_15_6A | GTCCCTGCCTCTATATCTTTCAGGCCAACACCGTTGGGATAGTTC |
| ana | soti | B3-ana_soti_RA_GF17485_15_7A | GTCCCTGCCTCTATATCTTTGCGAAGAACTCGCCGCGCTGCTCCT |
| ana | soti | B3-ana_soti_RA_GF17485_15_8A | GTCCCTGCCTCTATATCTTTTCAGGAAGAGGGGGCACAGCTCGGG |
| ana | soti | B3-ana_soti_RA_GF17485_15_9A | GTCCCTGCCTCTATATCTTTAATCGTGAAGAAGCTGCGTTTCTTG |
| ana | soti | B3-ana_soti_RA_GF17485_15_10A | GTCCCTGCCTCTATATCTTTCCGCCCGCCCTCGCCCCCTCGAACA |
| ana | soti | B3-ana_soti_RA_GF17485_15_11A | GTCCCTGCCTCTATATCTTTCCAGCAGCATTGCCACGTGTCGGTT |
| ana | soti | B3-ana_soti_RA_GF17485_15_12A | GTCCCTGCCTCTATATCTTTATGAGCCAGCTGCTGTTCCAGGACC |
| ana | soti | B3-ana_soti_RA_GF17485_15_13A | GTCCCTGCCTCTATATCTTTGAGTCGTTGCCCGCGTCTCCCGCAA |
| ana | soti | B3-ana_soti_RA_GF17485_15_14A | GTCCCTGCCTCTATATCTTTTATCGTCGGGATCCCTGTCCTCGGG |
| ana | soti | B3-ana_soti_RA_GF17485_15_15A | GTCCCTGCCTCTATATCTTTGTGTAAGTGTAAATTGAAACAATAA |
| ana | soti | B3-ana_soti_RA_GF17485_15_1B | TACTTTTCGTTGGATTTCGCTGCTTTTCCACTCAACTTTAACCCG |
| ana | soti | B3-ana_soti_RA_GF17485_15_2B | TCGGGAATTCAACTCATTGGCCAACTTCCACTCAACTTTAACCCG |
| ana | soti | B3-ana_soti_RA_GF17485_15_3B | TAACAACCCTAGTCCCTACCTAGTTTTCCACTCAACTTTAACCCG |
| ana | soti | B3-ana_soti_RA_GF17485_15_4B | CCCAAAGGTGACCTGGAGAGGAAGGTTCCACTCAACTTTAACCCG |
| ana | soti | B3-ana_soti_RA_GF17485_15_5B | TCGGATGTCCCCTCGCTCCTGCGACTTCCACTCAACTTTAACCCG |
| ana | soti | B3-ana_soti_RA_GF17485_15_6B | CAATCAGAACTCACCCCAGTGATGCTTCCACTCAACTTTAACCCG |
| ana | soti | B3-ana_soti_RA_GF17485_15_7B | CTGGTCATGTTTTCGAAGAGGTACTTTCCACTCAACTTTAACCCG |

| | | | |
|---|---|---|---|
| ana | soti | B3-ana_soti_RA_GF17485_15_8B | GGACCTCGCCTATCGCTCTGGAGGCTTCCACTCAACTTTAACCCG |
| ana | soti | B3-ana_soti_RA_GF17485_15_9B | CTGATTCCGGAGCACCGTCGGCCGCTTCCACTCAACTTTAACCCG |
| ana | soti | B3-ana_soti_RA_GF17485_15_10B | CTCGCCGGAGGGCGTGGATGGGCAATTCCACTCAACTTTAACCCG |
| ana | soti | B3-ana_soti_RA_GF17485_15_11B | CCATGGGCGGAGGCTCCAGTGGCGCTTCCACTCAACTTTAACCCG |
| ana | soti | B3-ana_soti_RA_GF17485_15_12B | CCGCCTCAGTGGATGATGTGCCTGATTCCACTCAACTTTAACCCG |
| ana | soti | B3-ana_soti_RA_GF17485_15_13B | TGCTCCAGTTGCTCCTCCTGGGAGTTTCCACTCAACTTTAACCCG |
| ana | soti | B3-ana_soti_RA_GF17485_15_14B | CGTCGCCAATCCGATCGATCAGCAGTTCCACTCAACTTTAACCCG |
| ana | soti | B3-ana_soti_RA_GF17485_15_15B | CACTACCTCCATCTGGGCGAGTGGTTTCCACTCAACTTTAACCCG |
| ana | b52 | B4-ana_B52_RA_GF17608_16_1A | CCTCAACCTACCTCCAACAAAGTTAATCATATAGACAATTTTTGA |
| ana | b52 | B4-ana_B52_RA_GF17608_16_2A | CCTCAACCTACCTCCAACAATTTGGCCCAGCGGGTGGGTGGGACT |
| ana | b52 | B4-ana_B52_RA_GF17608_16_3A | CCTCAACCTACCTCCAACAATCCATTATAATTTTTCTTTTTCTTT |
| ana | b52 | B4-ana_B52_RA_GF17608_16_4A | CCTCAACCTACCTCCAACAAAGTCTAAAATCAAGTGCTTTAAAAT |
| ana | b52 | B4-ana_B52_RA_GF17608_16_5A | CCTCAACCTACCTCCAACAAAGCGGGAGCGGGAGCGCGAGCGGGA |
| ana | b52 | B4-ana_B52_RA_GF17608_16_6A | CCTCAACCTACCTCCAACAATTCGATGCGGAGCGGGAGCGGGAAC |
| ana | b52 | B4-ana_B52_RA_GF17608_16_7A | CCTCAACCTACCTCCAACAACGAACGCGAGCGGGAACGGGAACGT |
| ana | b52 | B4-ana_B52_RA_GF17608_16_8A | CCTCAACCTACCTCCAACAAGGGAGCGGGACCTCCTGCGTGAGCG |
| ana | b52 | B4-ana_B52_RA_GF17608_16_9A | CCTCAACCTACCTCCAACAATCCACCAAGTGGATGCGTCTGCCAT |
| ana | b52 | B4-ana_B52_RA_GF17608_16_10A | CCTCAACCTACCTCCAACAACTTGTGGGCATCGGCATAGGTGACC |
| ana | b52 | B4-ana_B52_RA_GF17608_16_12A | CCTCAACCTACCTCCAACAAGCAATGGCGGCCCATAGCGGGAGGA |
| ana | b52 | B4-ana_B52_RA_GF17608_16_13A | CCTCAACCTACCTCCAACAAGCGACCGCCGCCGCCTCGTCGACCA |
| ana | b52 | B4-ana_B52_RA_GF17608_16_14A | CCTCAACCTACCTCCAACAACTGCCACGAGCGGTTCCCCTGGCGG |
| ana | b52 | B4-ana_B52_RA_GF17608_16_15A | CCTCAACCTACCTCCAACAACCTTGCCATTCAACTCATAGACAGC |
| ana | b52 | B4-ana_B52_RA_GF17608_16_16A | CCTCAACCTACCTCCAACAACACGAAGCCGTATCCATTCTTGATG |
| ana | b52 | B4-ana_B52_RA_GF17608_16_1B | TTCACATAACTTTATCATTTTTATTATTCTCACCATATTCGCTTC |
| ana | b52 | B4-ana_B52_RA_GF17608_16_2B | CCATAGATTGCTTATTTTTTTTTAAATTCTCACCATATTCGCTTC |
| ana | b52 | B4-ana_B52_RA_GF17608_16_3B | CGACACCTATTTCTGGAGTTTCAAGATTCTCACCATATTCGCTTC |
| ana | b52 | B4-ana_B52_RA_GF17608_16_4B | TAATGTTTTGTAGAGTAGTAAAGTAATTCTCACCATATTCGCTTC |
| ana | b52 | B4-ana_B52_RA_GF17608_16_5B | AGGCATTTCCGTTTTTGGGCGAGGCATTCTCACCATATTCGCTTC |
| ana | b52 | B4-ana_B52_RA_GF17608_16_6B | GAACGGGAACGGGAGCGGGACTCTCATTCTCACCATATTCGCTTC |
| ana | b52 | B4-ana_B52_RA_GF17608_16_7B | GGACTTGGAAACGTCGCGCGATTTGATTCTCACCATATTCGCTTC |
| ana | b52 | B4-ana_B52_RA_GF17608_16_8B | ACTTAGAGCGAGAGTGCGACGAGCGATTCTCACCATATTCGCTTC |
| ana | b52 | B4-ana_B52_RA_GF17608_16_9B | CCACCGCTGCGTCCGCCGCGACGATATTCTCACCATATTCGCTTC |
| ana | b52 | B4-ana_B52_RA_GF17608_16_10B | CTCCACAACGCCCTCGTTGCGTCGCATTCTCACCATATTCGCTTC |
| ana | b52 | B4-ana_B52_RA_GF17608_16_11B | CCAGCCTGGCGCATGTAGTCTTTCAATTCTCACCATATTCGCTTC |
| ana | b52 | B4-ana_B52_RA_GF17608_16_12B | TCTCCACAATTAGACGGTACTCAGTATTCTCACCATATTCGCTTC |
| ana | b52 | B4-ana_B52_RA_GF17608_16_13B | TCTGGAATTTTTGTTTTTGTCGTTGATTCTCACCATATTCGCTTC |
| ana | b52 | B4-ana_B52_RA_GF17608_16_14B | TAACGATCGTCGTAGCGGTCACGGTATTCTCACCATATTCGCTTC |
| ana | b52 | B4-ana_B52_RA_GF17608_16_15B | CGACAACCACGCGCTCGCCCAGCAGATTCTCACCATATTCGCTTC |
| ana | b52 | B4-ana_B52_RA_GF17608_16_16B | GTCGGCATCGCGGTAATCTTCAAATATTCTCACCATATTCGCTTC |
| ana | pkd2 | B5-ana_Pkd2_RA_GF14194_16_1A | CTCACTCCCAATCTCTATAAATTCCGTAACGTAATTCCATAAACA |
| ana | pkd2 | B5-ana_Pkd2_RA_GF14194_16_2A | CTCACTCCCAATCTCTATAATGGGCTTGAGTCATGTTACTGCGCA |
| ana | pkd2 | B5-ana_Pkd2_RA_GF14194_16_3A | CTCACTCCCAATCTCTATAATTTCCATGATAATGGCCAAGAACAT |
| ana | pkd2 | B5-ana_Pkd2_RA_GF14194_16_4A | CTCACTCCCAATCTCTATAACAGGAATACAATTCCGAACATAAGA |
| ana | pkd2 | B5-ana_Pkd2_RA_GF14194_16_5A | CTCACTCCCAATCTCTATAAGTTACCGACTTGACCTTAAAGGTAT |
| ana | pkd2 | B5-ana_Pkd2_RA_GF14194_16_6A | CTCACTCCCAATCTCTATAAAGAATAGTTTACTGTTTGAAGTTG |
| ana | pkd2 | B5-ana_Pkd2_RA_GF14194_16_7A | CTCACTCCCAATCTCTATAAGCCGCCGCTGCGATAGAAAGCCAAT |
| ana | pkd2 | B5-ana_Pkd2_RA_GF14194_16_8A | CTCACTCCCAATCTCTATAAGCAGCTGTCCTTTTTCACCCGAATC |
| ana | pkd2 | B5-ana_Pkd2_RA_GF14194_16_9A | CTCACTCCCAATCTCTATAACTCTTCTTGTCGCACCTTCCTTCTG |
| ana | pkd2 | B5-ana_Pkd2_RA_GF14194_16_10A | CTCACTCCCAATCTCTATAAGGGTGGTGGTTCGGCAGGTTTAGTC |
| ana | pkd2 | B5-ana_Pkd2_RA_GF14194_16_11A | CTCACTCCCAATCTCTATAATGTGTCATTGAAATAGAACATGTAC |
| ana | pkd2 | B5-ana_Pkd2_RA_GF14194_16_12A | CTCACTCCCAATCTCTATAACGGCCCGACACCGGATCGCCGCTCC |
| ana | pkd2 | B5-ana_Pkd2_RA_GF14194_16_13A | CTCACTCCCAATCTCTATAAGCCGAAAATATTGAATCCGCTGGAT |
| ana | pkd2 | B5-ana_Pkd2_RA_GF14194_16_14A | CTCACTCCCAATCTCTATAAAAGTGGGGAAGTCGATCTGGCCACA |
| ana | pkd2 | B5-ana_Pkd2_RA_GF14194_16_15A | CTCACTCCCAATCTCTATAACTTGCCGCGTGGAGATCCATCGCCA |
| ana | pkd2 | B5-ana_Pkd2_RA_GF14194_16_16A | CTCACTCCCAATCTCTATAAAGTTGGTGGCGGTTTCCGGGGAGAT |
| ana | pkd2 | B5-ana_Pkd2_RA_GF14194_16_1B | TTCTGAAGAATCAAAACAGTTTATTAACTACCCTACAAATCCAAT |
| ana | pkd2 | B5-ana_Pkd2_RA_GF14194_16_2B | GGCGGAATTTCTTGGAAATACTTGGAACTACCCTACAAATCCAAT |
| ana | pkd2 | B5-ana_Pkd2_RA_GF14194_16_3B | TGATCTCACTCTTCACCGAGTTGTAAACTACCCTACAAATCCAAT |
| ana | pkd2 | B5-ana_Pkd2_RA_GF14194_16_4B | TAATAGGAGGCCCAGCTGAGCATAGAACTACCCTACAAATCCAAT |
| ana | pkd2 | B5-ana_Pkd2_RA_GF14194_16_5B | TAGGTCAACTGAGTCGTGGCCTTGTAACTACCCTACAAATCCAAT |
| ana | pkd2 | B5-ana_Pkd2_RA_GF14194_16_6B | TTAGTAGGCTGCGTTCCGTGAAGAAAACTACCCTACAAATCCAAT |
| ana | pkd2 | B5-ana_Pkd2_RA_GF14194_16_7B | CTTTTCGAATGTCAAGTCCATCACGAACTACCCTACAAATCCAAT |

| | | | |
|---|---|---|---|
| ana | pkd2 | B5-ana_Pkd2_RA_GF14194_16_8B | GTATCGGATGAAGGCGTCATTGACGAACTACCCTACAAATCCAAT |
| ana | pkd2 | B5-ana_Pkd2_RA_GF14194_16_9B | CGAGTCTCCCTCTTCTGCCTCGGGAAACTACCCTACAAATCCAAT |
| ana | pkd2 | B5-ana_Pkd2_RA_GF14194_16_10B | TGGAGGGGGCTGTGGAGGGGGCGATAACTACCCTACAAATCCAAT |
| ana | pkd2 | B5-ana_Pkd2_RA_GF14194_16_11B | TTCTCGGGTTGTAAAGAGCTTCTTCAACTACCCTACAAATCCAAT |
| ana | pkd2 | B5-ana_Pkd2_RA_GF14194_16_12B | GGCTTTGGTCTTATTTTGAACAGCTAACTACCCTACAAATCCAAT |
| ana | pkd2 | B5-ana_Pkd2_RA_GF14194_16_13B | CTTGGGGGGCTCTTTTGGCTTCTTTAACTACCCTACAAATCCAAT |
| ana | pkd2 | B5-ana_Pkd2_RA_GF14194_16_14B | TTGTGGCGACTGAGTAGCGGCCGCGAACTACCCTACAAATCCAAT |
| ana | pkd2 | B5-ana_Pkd2_RA_GF14194_16_15B | CCTTATTACTAGTCGGGGCGAAGCAAACTACCCTACAAATCCAAT |
| ana | pkd2 | B5-ana_Pkd2_RA_GF14194_16_16B | ACCTCTTATGACCCGCCTTGCGGGAAACTACCCTACAAATCCAAT |

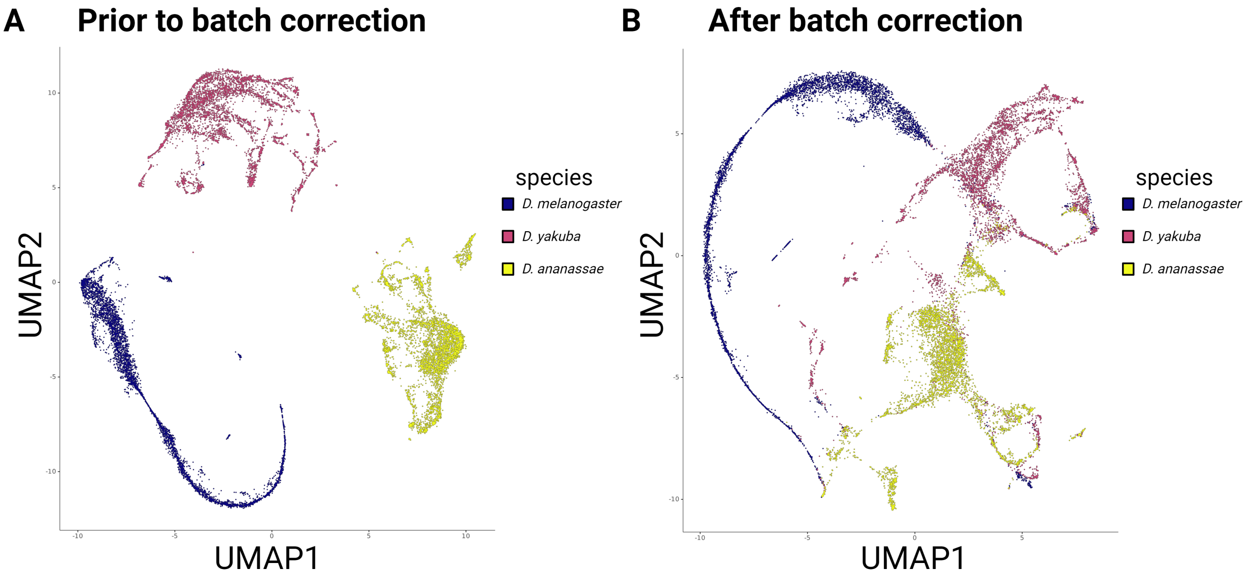

**Figure S1: Batch correction for species in testis data still shows significant species-specific differences. (A)** UMAP visualization of scRNA-Seq data from testis in *D. melanogaster*, *D. yakuba*, and *D. ananassae* shows clusters only correlating with species. **(B)** UMAP visualization of batch-corrected scRNA-Seq data shows persistent segregation by species.

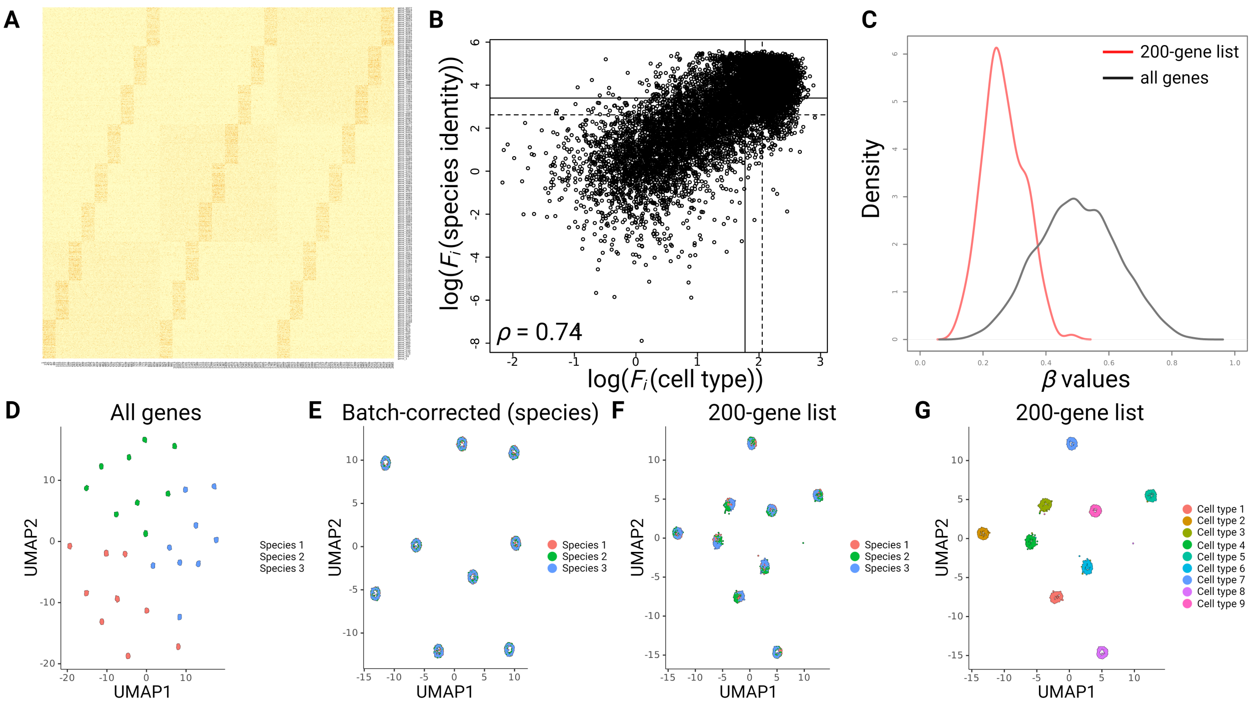

**Figure S2: Application of the methodology to simulated data suggests spermatogenesis undergoes strong species-specific cell type divergence**. **(A)** Heat map of simulated data. Each row represents the ortholog of a single-copy gene in its respective species, while each column represents a single cell. Darker color represents increased expression. Cells 1-300 are from species 1, cells 301-600 are from species 2, cells 601-900 are from species 3. Elevated expression of genes 1-1111 are indicative of cell type 1, 1112-2222 is indicative of cell type 2, 2223-3333 is indicative of cell type 3, etc. The genes reflecting cell types 1, 2, and 3 have increased their expression in species 1, the genes reflecting cell types 4, 5, and 6 have increased their expression in species 2, and the genes reflecting cell types 7, 8, and 9 have increased their expression in species 3. As a result, the heat map shows three sets of 9 blocks, representing cell types 1-9 in species 1-3. The larger three boxes reflect a species-specific increase in gene expression for cell types 1-3 (in species 1), cell types 4-6 (in species 2), and cell types 7-9 (in species 3). **(B)** ANOVA $F$-statistic distributions of simulated data show a high correlation between cell type and species identity resulting from the underlying data structure. Solid lines correspond with median marginal distributions, while dotted lines correspond to cut-off values used to generate the top-200 gene list. **(C)** Distributions of simulated $\beta$ values show that the top-200 gene list preferentially contains cell type information over species identity information. **(D)** UMAP visualization of simulated data shows clustering both by cell type and species identity. **(E)** Batch correction for species identity on simulated data recovers cell type information but ablates all species information. **(F-G)** Utilization of top-200 gene list recovers cell type clusters while still preserving species-specific differences in cell type.

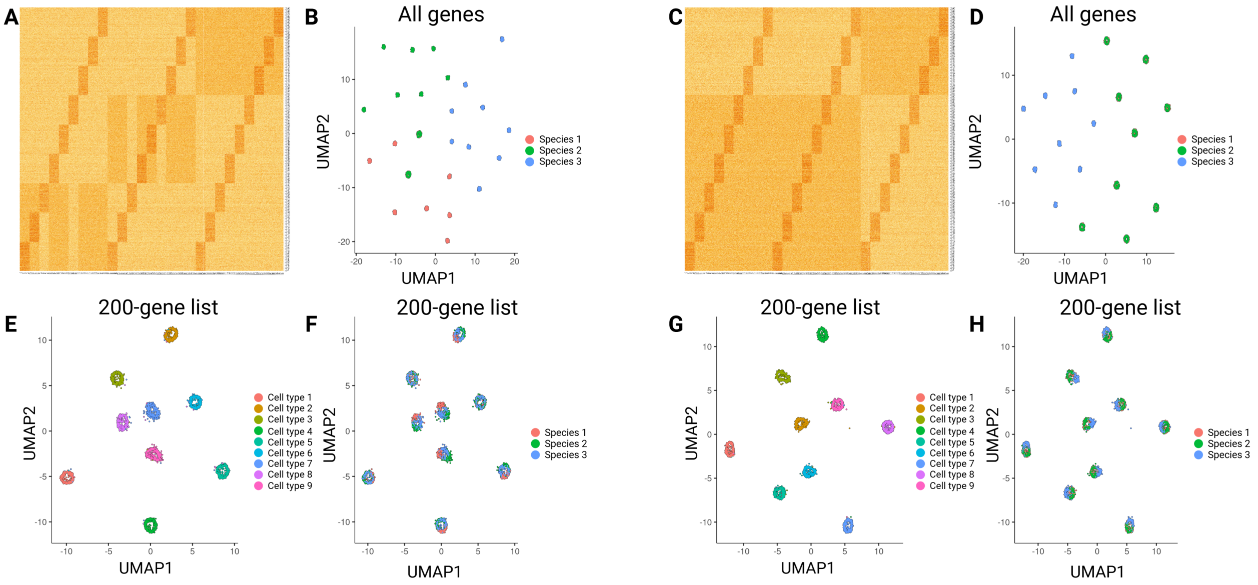

**Figure S3: Application of the methodology to data simulating asymmetric phylogenies demonstrates efficacy across differing divergence time ratios**. Two simulated data sets reflecting asymmetric phylogenies with either moderately (**A, B**) or maximally (**C, D**) asymmetric relatedness. Heat maps (**A, C**) and UMAP plots (**B, D**) are presented for these phylogenies. UMAP clustering resulting from utilizing the top 200 marker gene sets show that the effectiveness of the statistical procedure in moderately (**E, F**) and maximally (**G, H**) asymmetric scenarios.

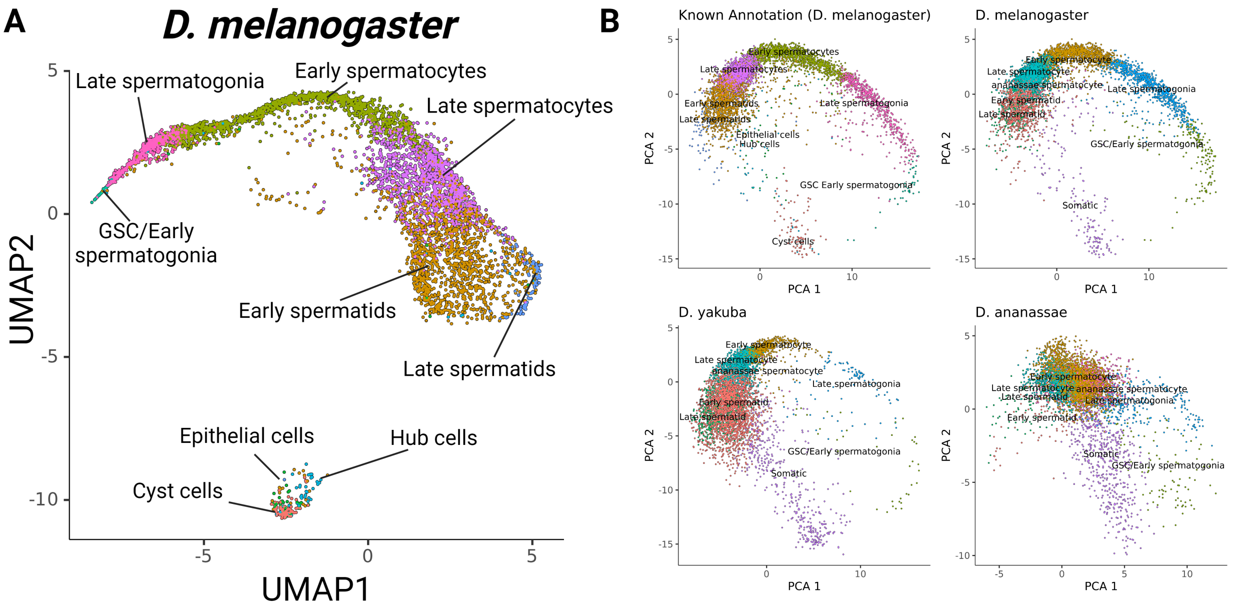

**Figure S4: UMAP and PCA projection over 198-gene cluster recovers known cell type information while showing evolutionarily conserved developmental transcriptomic divergence.** **(A)** Overlay of previously identified cell type classification on UMAP projection using the 198-gene list. **(B)** Overlay of newly assigned cell type classifications on PCA projections shows that developmental transcriptomic divergence is not an artefact of manifold project. This divergence is also evolutionarily conserved.

A

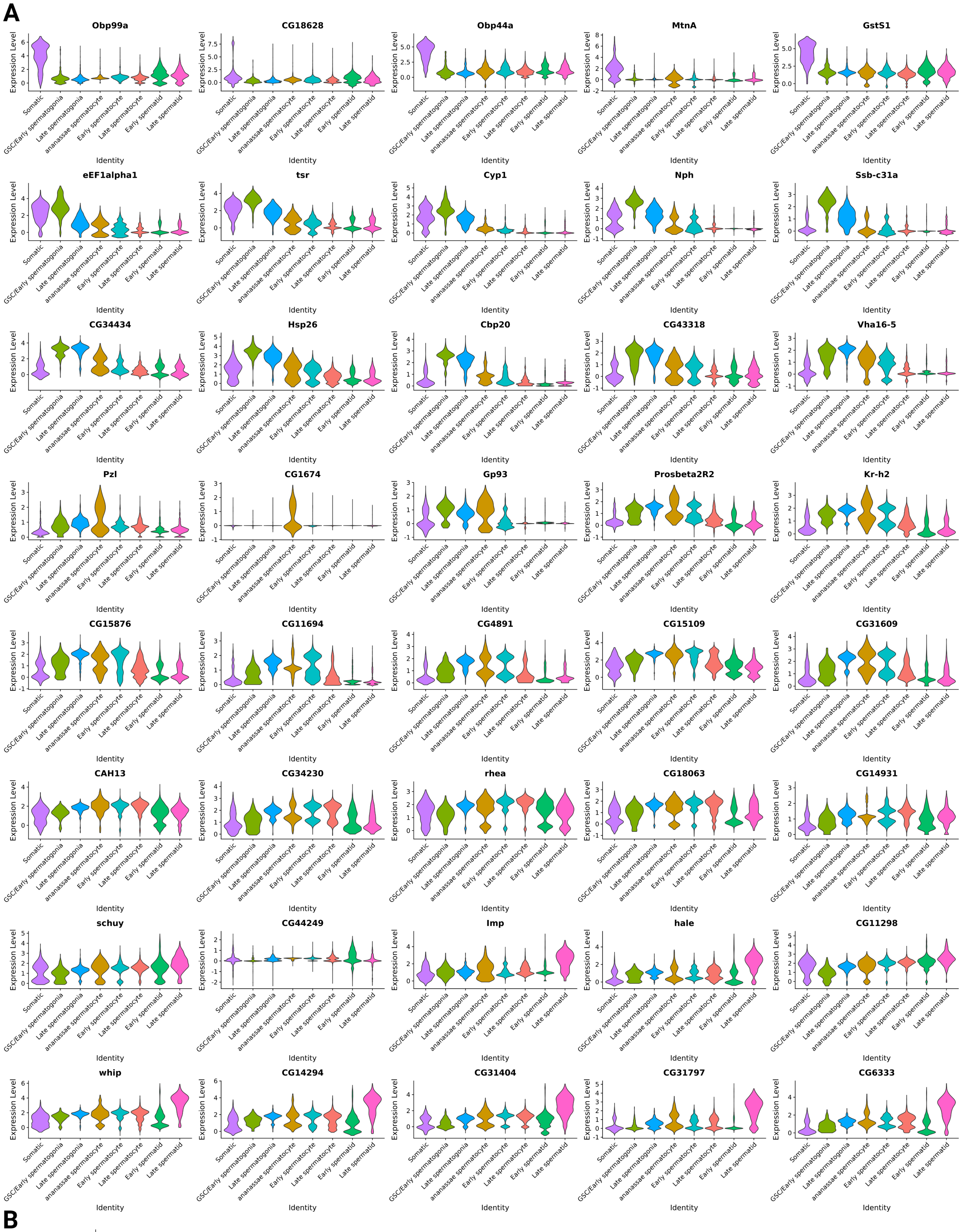
Obp99a
CG18628
Obp44a
MtnA
GstS1
eEF1alpha1
tsr
Cyp1
Nph
Ssb-c31a
CG34434
Hsp26
Cbp20
CG43318
Vha16-5
Pzl
CG1674
Gp93
Prosbeta2R2
Kr-h2
CG15876
CG11694
CG4891
CG15109
CG31609
CAH13
CG34230
rhea
CG18063
CG14931
schuy
CG44249
Imp
hale
CG11298
whip
CG14294
CG31404
CG31797
CG6333
Expression Level
Identity
Somatic
GSC/Early spermatogonia
Late spermatogonia
ananassae spermatocyte
Early spermatocyte
Late spermatocyte
Early spermatid
Late spermatid

B

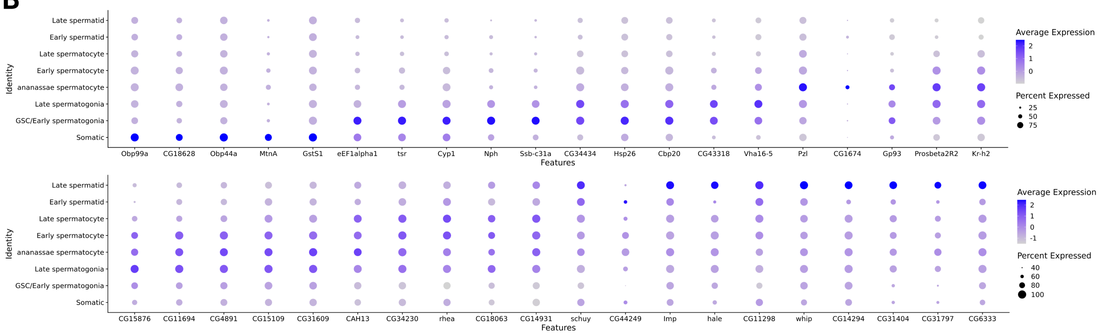
Identity
Late spermatid
Early spermatid
Late spermatocyte
Early spermatocyte
ananassae spermatocyte
Late spermatogonia
GSC/Early spermatogonia
Somatic
Obp99a
CG18628
Obp44a
MtnA
GstS1
eEF1alpha1
tsr
Cyp1
Nph
Ssb-c31a
CG34434
Hsp26
Cbp20
CG43318
Vha16-5
Pzl
CG1674
Gp93
Prosbeta2R2
Kr-h2
Features
Average Expression
Percent Expressed
CG15876
CG11694
CG4891
CG15109
CG31609
CAH13
CG34230
rhea
CG18063
CG14931
schuy
CG44249
Imp
hale
CG11298
whip
CG14294
CG31404
CG31797
CG6333

**Figure S5: Violin and dot plots for most differentially expressed genes between cluster reveal developmental trajectories of highly expressed genes**. **(A, B)** Violin and dot plots for most highly differentially expressed genes show a progression of expression across spermatogenesis.

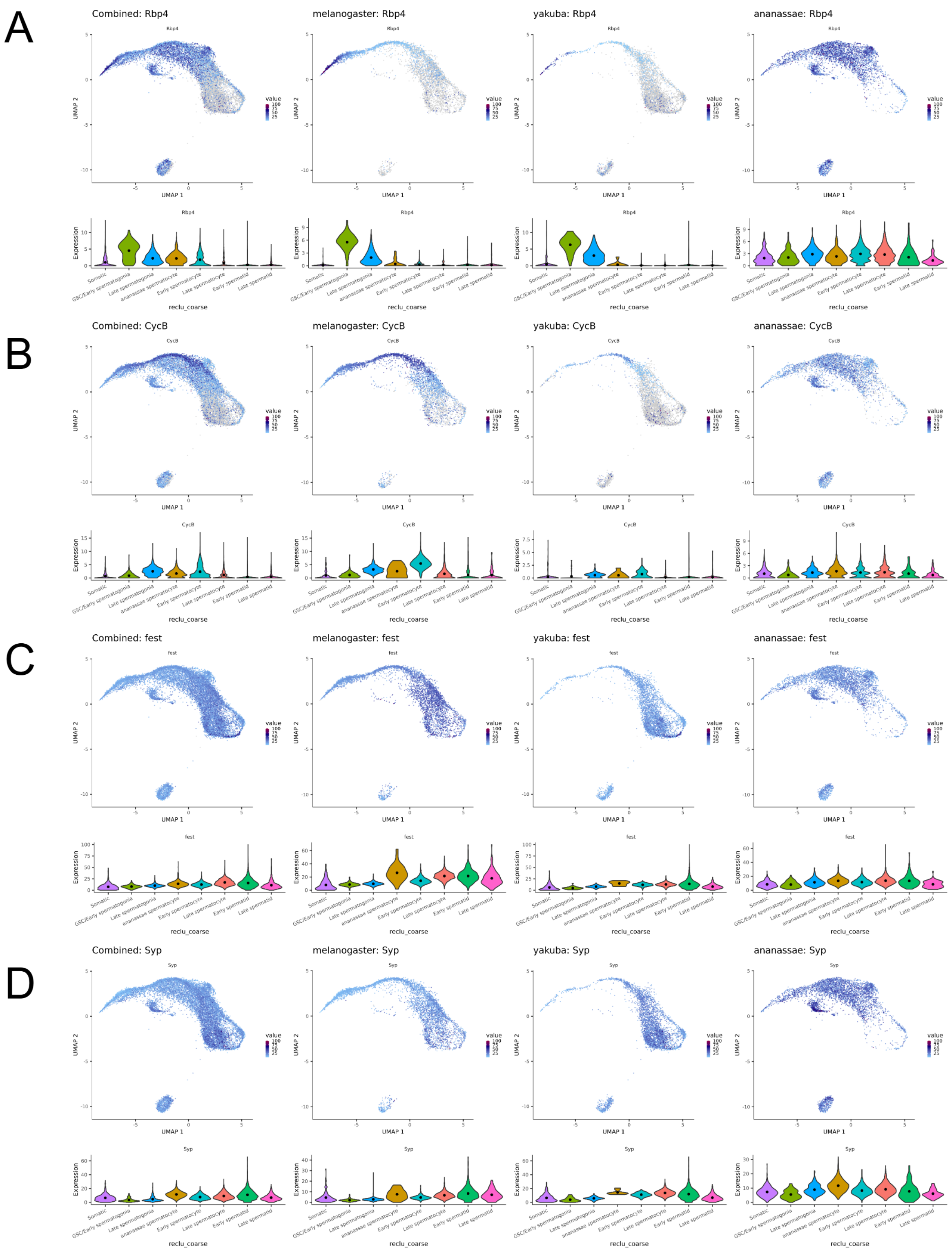

**Figure S6: Expression of *Rbp4*-related genes**. Cell type-specific expression patterns in *D. melanogaster*, *D. yakuba*, and *D. ananassae* for **(A)** *Rpb4* **(B)** *Cyclin B* **(C)** *Fest* and **(D)** *Syp*.

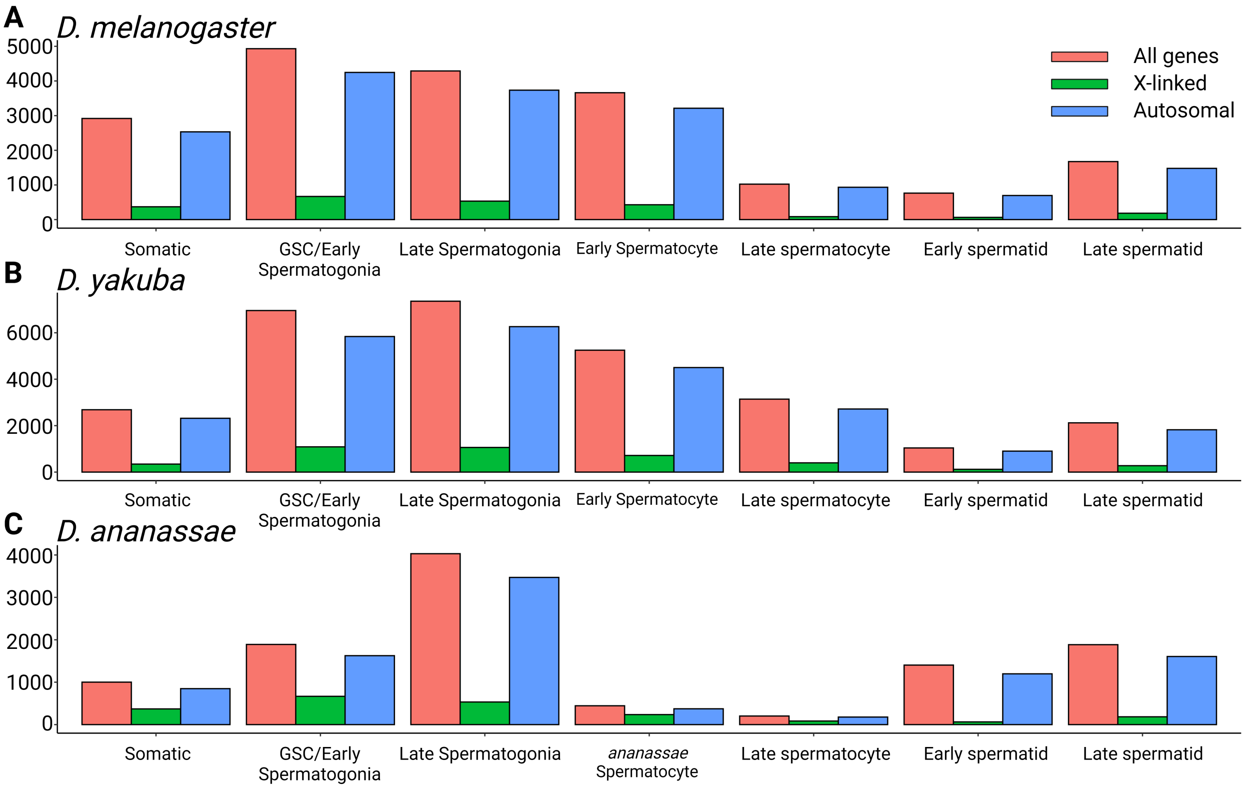

**Figure S7: Number of genes that pass quality control**. **(A-C)** Total number of genes that pass quality control by chromosome. The relative ratio of X-linked to autosomal genes is constant throughout spermatogenesis, indicating that there is not a specific stage during spermatogenesis at which X-inactivation begins.

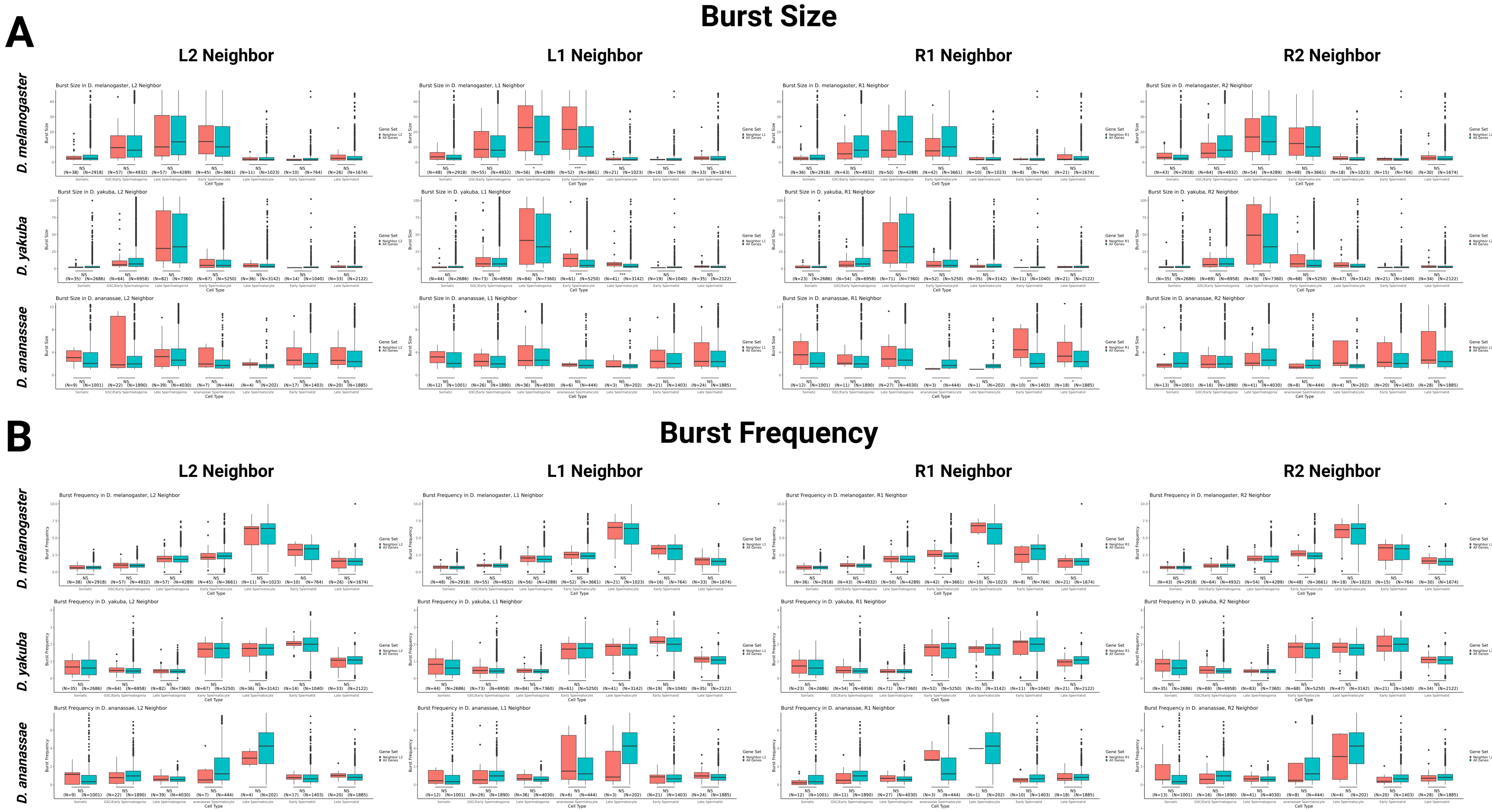

A
Burst Size
L2 Neighbor
L1 Neighbor
R1 Neighbor
R2 Neighbor
D. melanogaster
D. yakuba
D. ananassae
B
Burst Frequency
L2 Neighbor
L1 Neighbor
R1 Neighbor
R2 Neighbor
D. melanogaster
D. yakuba
D. ananassae

**Figure S8: Bursting kinetics for genes neighboring *de novo* transcripts across cell type and species**. (A) Burst size and (B) burst frequency for L2, L1, R1, and R2 neighbors and all genes are plotted above for *D. melanogaster*, *D. yakuba*, and *D. ananassae* (* = $p < 0.05$, ** = $p < 0.01$, *** = $p < 0.001$, **** = $p < 0.0001$. Note these are raw p-values prior to application of the Benjamini-Hochberg, FDR $< 0.05$). Here, *D. melanogaster* and *D. yakuba* are in-groups while *D. ananassae* is an out-group.

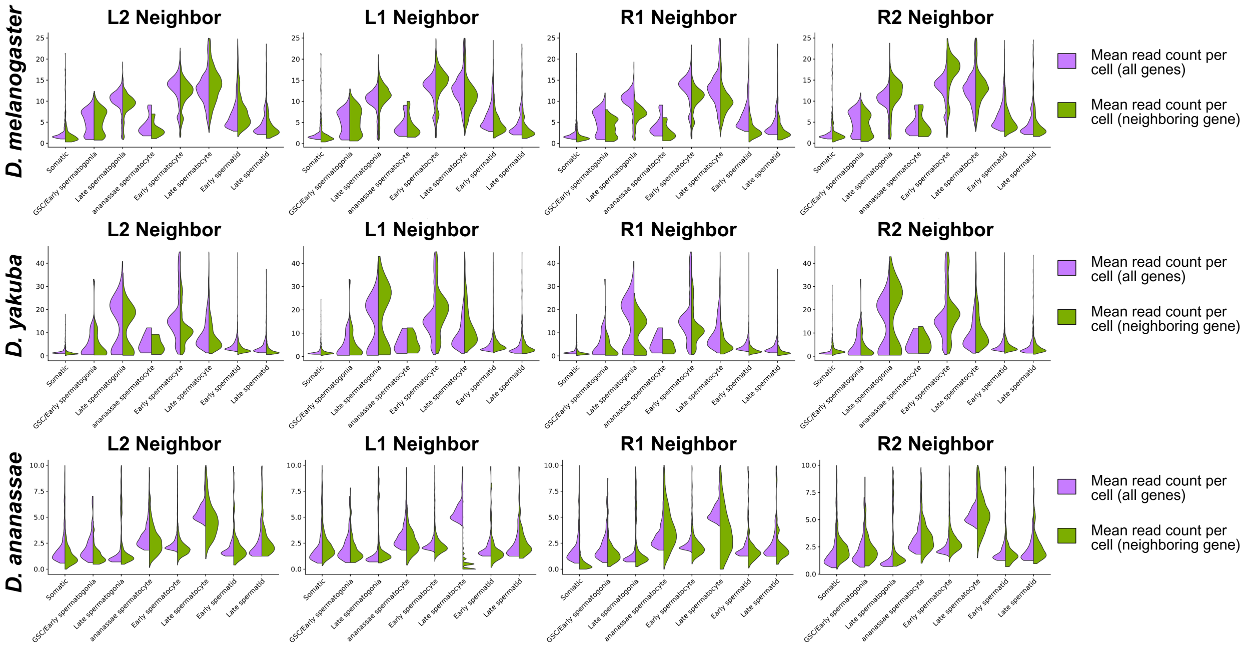

**Figure S9: Mean read count (expression) per cell for genes neighboring *de novo* transcripts across cell type and species**. Mean read count per cell for L2, L1, R1, and R2 neighbors and all genes are plotted above for *D. melanogaster*, *D. yakuba*, and *D. ananassae*. Note significantly increased expression per cell for L1 neighbors in early spermatocytes for *D. melanogaster* and *D. yakuba*, with no significant difference detected for the same cell type in *D. ananassae.* Similarly, expression is significantly decreased for R1 neighbors in early spermatocytes in *D. melanogaster* and *D. yakuba*, with no significant difference detected for the same cell type in *D. ananassae*. Significance is assessed using a t-test after multiple hypothesis correction - raw p-values for this analysis are found in Data S2.
